# Genomic Maxwell’s Demon Control of Cancer Cell Fates: Integrated Biophysical Mechanisms of Fate Commitment

**DOI:** 10.64898/2026.02.12.703632

**Authors:** Masa Tsuchiya, Kenichi Yoshikawa, Oleg Naimark

## Abstract

Molecular biology has revealed immense spatiotemporal complexity in cancer regulatory networks. A fundamental physical question remains: Does a physical principle exist that governs how this complexity self-organizes into genome-wide fate commitment and determines whether and when that commitment becomes irreversible?

We address this question at the coarse-grained level of genome-wide expression dynamics through integrated biophysical analyses of time-series transcriptomes from two cancer systems under four conditions: MCF-7 human breast cancer cells stimulated with heregulin (HRG) or epidermal growth factor (EGF), and HL-60 human leukemia cells stimulated with all-trans retinoic acid (atRA) or dimethyl sulfoxide (DMSO).

This study further develops our previous framework in which the critical point (CP) gene ensemble regulates self-organized criticality (SOC) in the whole expression system (WES), with CP-state changes triggering genome-wide expression avalanches during cell-fate transitions. Here, SOC control unifies information thermodynamics with genome-engine dynamics in an open, non-equilibrium WES, maintaining a critical dynamic balance between homeostatic stability and fate-guiding critical transitions.

The CP performs two integrated roles. As a genomic Maxwell’s demon (MD) operator, the CP links information acquisition to feedback-associated state rewriting; reconfiguration of the CP’s bimodal expression pattern provides operational rewritable memory. As an SOC controller, it synchronizes with the genome attractor and generates the dominant contribution to work in expression space, driving genome-wide reorganization. Thermodynamic phase synchronization between the CP and the peripheral expression system (PES; the WES excluding the CP), together with information-work coupling, integrates these information-thermodynamic and dynamical roles.

Across all four conditions, the CP guides both thermodynamic and dynamical phase synchronization to regulate fate commitment and subsequent genome-wide execution. The timing of information-work coupling distinguishes differentiation-inducing HRG, atRA, and DMSO from proliferative EGF. Based on these results, we propose four criteria for the irreversible arrow of cell-fate change in expression space and define time-gated rules for dynamic cancer-fate control. These criteria provide a testable framework for assessing physical thermodynamic irreversibility through entropy calibration and measurement of outward entropy export.

## 1. Introduction

Molecular biology has revealed immense spatiotemporal complexity across cancer-cell regulatory networks [Hanahan and Weinberg, 2011; Hanahan, 2026]. Recent cancer biology studies show that this complexity spans genetic alterations and chromatin-epigenetic regulation [Martínez-Jiménez et al., 2020; Esteller et al., 2024], signaling and transcriptional dynamics [Levchenko and Madsen, 2026; Davies et al., 2023], metabolic remodeling [Xu et al., 2023], and microenvironmental responses [de Visser and Joyce, 2023]. Pathway and gene-set resources reflecting this complexity include KEGG [Kanehisa et al., 2025], Reactome [Ragueneau et al., 2026], and MSigDB [Liberzon et al., 2015]. At the genome scale, chromosomal instability and structural rearrangements (reviewed in [Chen et al., 2025]), together with dynamic rewiring of higher-order chromatin architecture [Jang and Yoo, 2026], further contribute to this spatiotemporal complexity.

Together, these biological insights raise a fundamental physical question:

Does a system-level physical principle exist that controls how this complexity self-organizes into genome-wide fate commitment and determines whether and when that commitment establishes irreversible cancer cell fate?

We address this question by treating cancer cell fate as an emergent property of the whole expression system (WES) at a coarse-grained level. This WES exchanges matter, energy, and information with the environment and operates as an open, non-equilibrium, self-organizing thermodynamic network (Section 2.1).

Our previous studies identified the critical point (CP) gene ensemble as the regulator of self-organized criticality (SOC) in the whole expression system (WES), with CP-state changes triggering genome-wide expression avalanches during cell-fate transitions [Tsuchiya et al., 2014, 2015, 2016; Giuliani et al., 2018; Tsuchiya et al., 2020, Zimatore et al.,2021; Tsuchiya et al., 2023a]. We further demonstrated SOC-controlled genome dynamics across cell-fate transitions in cancer cell populations and T helper 17 differentiation, and at the single-cell level in mouse and human embryonic development [Tsuchiya et al., 2016, 2020]. The associated analytical methods have been progressively developed through these studies [Tsuchiya et al., 2023b].

One aim of the present study is to further advance our understanding of the SOC-control mechanism and to develop the associated analytical methods.

To identify the underlying non-equilibrium genomic mechanism, we integrate three previously developed analyses grounded in the same statistically validated convergence property of genome-wide expression (**Section 2.2**):

- **Information-thermodynamic analysis** (ITA) [Tsuchiya et al., 2025], which quantifies entropy, interdomain mutual information, and their temporal dynamics (Section 2.3);
- **Chromatin state analysis** (CSA) [Tsuchiya et al., 2025], which tracks chromatin structural change through a transcriptome-derived chromatin-remodeling proxy (Section 2.3); and
- **Expression flux analysis** (EFA) [Tsuchiya et al., 2020, 2024], which resolves genome-engine dynamics as an environment-coupled effective-force network among domain attractors. It also defines domain-specific effective mechanical work and reveals attractor synchronization dynamics (Section 2.4).

All three analyses build on coherent stochastic behavior (CSB) (Section 2.2), in which the averages of randomly selected gene ensembles converge to stable values as the number of genes increases, despite substantial fluctuations in individual genes [Tsuchiya et al., 2014, 2015, 2016]. This convergence enables macroscopic thermodynamic, chromatin-state, and dynamical analyses at a coarse-grained level despite single-gene noise.

We apply this integrated biophysical framework to time-series whole-transcriptome data from two cancer systems under four conditions. MCF-7 human breast cancer cells are stimulated with heregulin (HRG, cell differentiation) or epidermal growth factor (EGF, cell proliferation without cell-fate change). HL-60 human leukemia cells are stimulated with all-trans retinoic acid (atRA, cell differentiation) or dimethyl sulfoxide (DMSO, cell differentiation).

By comparing these four conditions with two additional replicate transcriptome datasets from MCF-7 cells, we develop a data-driven biophysical framework that unifies information thermodynamics [Tsuchiya et al., 2025] with genome-engine dynamics [Tsuchiya et al., 2016, 2020] through self-organized criticality (SOC) control of the WES (**Figure 1**). Importantly, this coarse-grained formulation averages over microscopic biological details while revealing emergent collective expression dynamics that underlie the unified biophysical principle.

**Figure 1.**
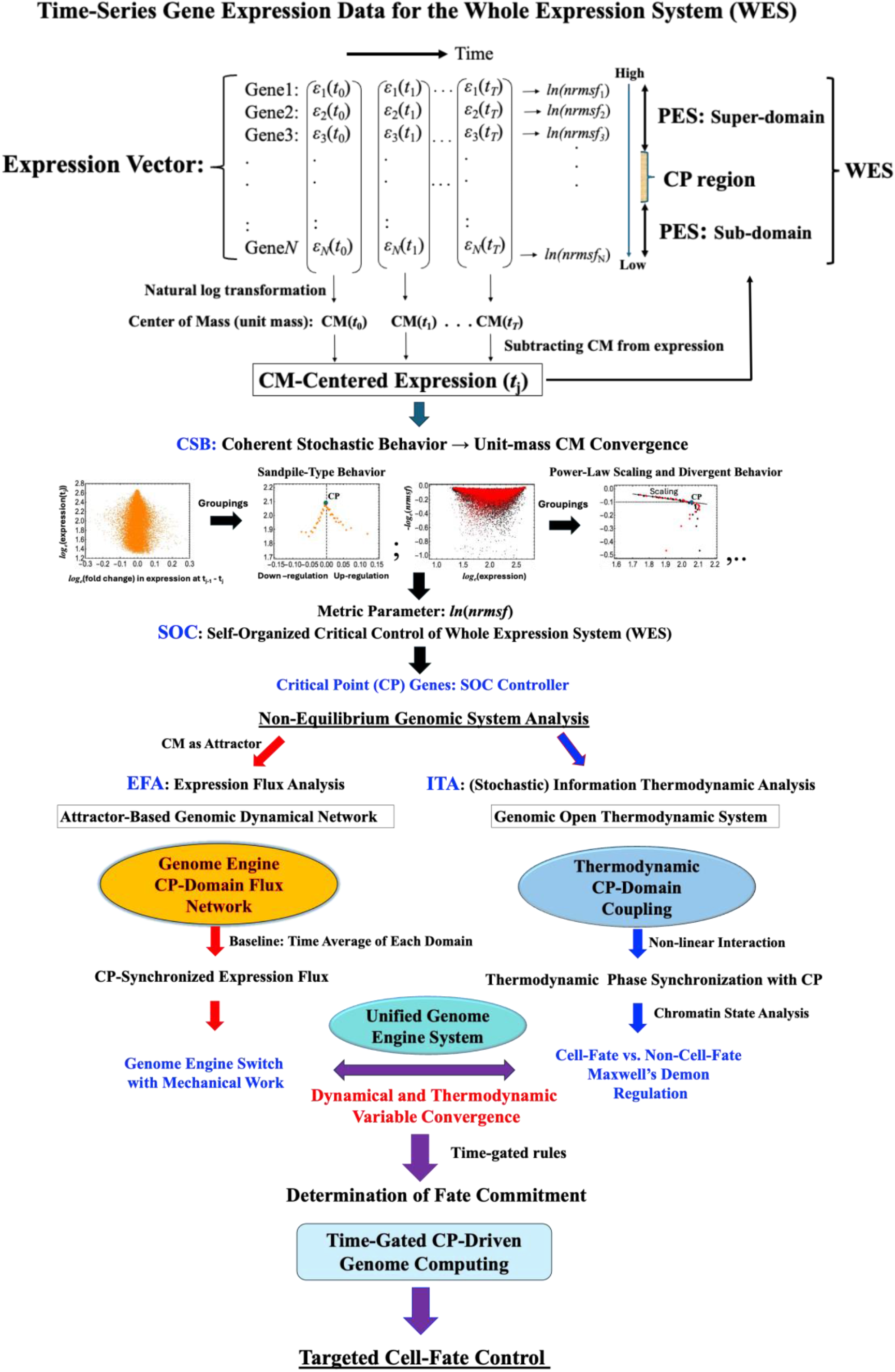
Unified scheme integrating dynamical-system and information-thermodynamic frameworks for genome regulation. Conceptual flowchart showing how the Critical Point (CP) drives self-organized criticality (SOC)-controlled genome dynamics and information-thermodynamic regulation of cell-fate commitment. **(Top)** Time-series gene expression vectors are log-transformed and centered on the unit-mass CM (ensemble average) to reveal coherent stochastic behavior. Convergence of the ensemble average establishes the macroscopic genome attractor (GA) of the whole expression system (WES). Its temporal trajectory provides the reference for subsequent analyses. Self-organized criticality (SOC) analysis along *ln*(nrmsf) identifies the CP region and partitions the WES into the Super-domain, CP region, and Sub-domain, with the Super- and Sub-domains together defining the peripheral expression system (PES). **(Middle)** Two complementary non-equilibrium stochastic analyses are applied in parallel. **The dynamical-system view (left**; **EFA)** reconstructs the effective expression-derived (EED) force interaction network among domain attractors. It identifies network reorganization associated with the genome-engine (GE) switch and quantifies domain-resolved EED work and CP-GA phase synchronization. **The stochastic information-thermodynamic view (right; ITA)**, with convergence-validated estimates, treats the whole expression system (WES) as an open, non-equilibrium thermodynamic system. It quantifies CP-PES information coupling, including pairwise and higher-order CP-PES information interactions that together drive thermodynamic phase synchronization. Chromatin state analysis bridges both views by relating the CP-driven GE switch to Maxwell’s demon (MD)-guided rewritable chromatin regulation, thereby linking synchronized dynamical and thermodynamic signatures to transcriptome-derived CP-guided chromatin state transitions. **(Bottom**) The two views converge into a unified genome engine system in which MD-cycle timing, the GE switch, CP-PES information coupling, and EED work define the progression from fate commitment to execution and provide the basis for CP-driven genome computing.

CSB convergence defines the temporal ensemble average of the WES trajectory as the genome attractor (GA), which provides a stable reference for temporal fluctuations and enables rigorous non-equilibrium dynamical and stochastic information-thermodynamic analyses. Furthermore, our previous study [Zimatore et al., 2021] demonstrated that chromatin remodeling serves as the material basis for SOC control of the WES (Section 2.3). Accordingly, the expression-variability coordinate, *ln*(*nrmsf*) (natural log of normalized root mean square fluctuation; see Section 2.1), provides a quantitative expression-derived proxy for chromatin remodeling at the coarse-grained level of collective genome-expression dynamics.

Our previous studies show that CSB reveals two hallmark SOC behaviors in gene-expression groups across the WES (**Figure 1**): sandpile-type critical behavior and scaling-divergent behavior. In the sandpile-type critical behavior, the summit corresponds to a group of genes exhibiting critical behavior, defined as the critical point (CP). Small changes in the CP state trigger genome-wide amplification [Tsuchiya et al., 2015, 2016], which propagates as avalanche-like events [Tsuchiya et al., 2023a] (Section 2.4). In the scaling-divergent behavior, collective power-law scaling, independent of group size, is characteristic of scale-free behavior, whereas near and beyond the CP boundary, the scaling pattern departs from the power-law relation and becomes divergent. These behaviors reflect collective genome-expression dynamics centered on the CP.

CSB further reveals collective behavior of the CP gene group, with coordinate-dependent critical dynamics (Fig. 2 in [Tsuchiya et al., 2023b]). Throughout this study, we analyze CP behavior along the *ln*(*nrmsf*) coordinate, where the temporal change in the ensemble average of the CP gene group exhibits bimodal singular behavior (Fig. 1 in [Tsuchiya et al., 2023b]).

**Figure 2.**
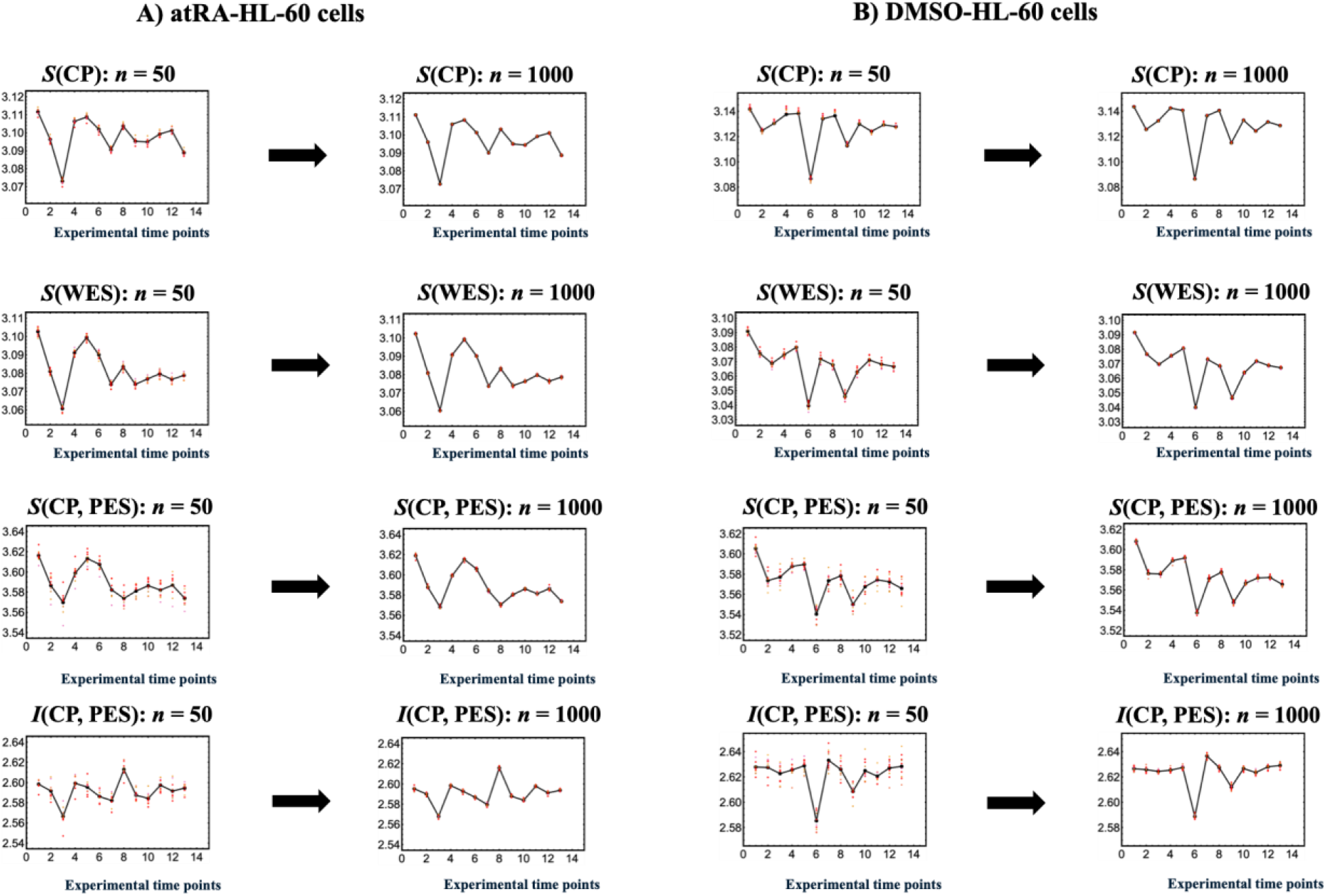
Convergence and robustness of information-thermodynamic metrics with respect to the number of bootstrap iterations. This figure compares temporal profiles of key information-thermodynamic metrics for **(A)** atRA-stimulated HL-60 cells in the first two columns and **(B)** DMSO-stimulated HL-60 cells in the last two columns. For HRG- and EGF-stimulated MCF-7 cells, see Fig. 4 in [Tsuchiya et al., 2025]. Bootstrapping is performed on the CP, PES, and WES expression vectors with the following fixed parameters: a random sample size of 1000 genes and 30 bins. Within each condition, *n*_iter_ = 50 is shown in the left column and *n*_iter_ = 1000 in the right column. For each condition, the full bootstrap procedure is repeated *n*_boot_ = 10 times. Red points show the scatter across these 10 runs, which is used to summarize run-to-run uncertainty (SD, see main text). Black lines show the mean across runs. The metrics plotted are *S*(CP), *S*(WES), *S*(CP,PES), and *I*(CP;PES), where *S*(WES) ≠ *S*(CP,PES) (see Section 2.3.1 for details). Convergence is confirmed when increasing *n*_iter_ from 50 to 1000 changes the run-averaged metric by only *O*(10^-3^) at each time point. The near overlap between the *n*_iter_ = 50 and *n*_ite*r*_ = 1000 ensemble-mean profiles, together with the clustering of the red points around the ensemble-mean profiles, indicates that all metrics have effectively converged, with differences between the ensemble-mean profiles remaining *O*(10^-3^).

Hereafter, this CP gene ensemble is referred to simply as the CP.

In this study, we identify two operational roles of the CP along *ln*(*nrmsf*):

**i) Dynamically**, the CP is the internal SOC regulator of the WES [Tsuchiya et al., 2016, 2020]. Unlike classical SOC models [Bak et al., 1987; Watkins et al., 2016], in which the critical state develops under slow driving and dissipation without a distinct internal regulatory subsystem, genomic SOC is dynamically maintained through internal regulation by the CP; the WES thereby operates as an autonomous dynamical system in which homeostasis and critical-transition dynamics coexist in a critical dynamic balance [Tsuchiya et al., 2023b]. This genomic SOC organization connects genomic regulation to the broader framework of criticality and dynamical scaling in living systems [Muñoz, 2018].

In Section 2.1, along *ln*(*nrmsf*), the WES is partitioned into three response domains: CP, Super (higher *ln*(*nrmsf*)), and Sub (lower *ln*(*nrmsf*)). In Section 2.2, we statistically validate CSB. In Section 2.4, this validation provides the statistical basis for treating the ensemble average of each domain as a one-dimensional dynamical macrovariable representing a domain attractor. EFA then constructs an interaction network of expression-derived (EED) forces among these attractors and defines domain-resolved EED work. The temporal reorganization of this interaction network near a critical transition defines the genome-engine (GE) switch [Tsuchiya et al., 2016, 2020, 2024].

Furthermore, the *ln*(*nrmsf*) coordinate serves as an SOC metric and chromatin-remodeling proxy [Zimatore et al., 2021; Tsuchiya et al., 2025], linking genome dynamics to chromatin structural dynamics. CP-guided fluctuations organize transcriptome-wide power-law scaling that is consistent with SOC and coordinated with chromatin structural transitions [Zimatore et al., 2021]. In HRG-stimulated MCF-7 cells, these transitions are validated by independent experimental evidence of pericentromere-associated domain (PAD) structural changes [Krigerts et al., 2021], whose timing and power-law scaling match the SOC analysis [Zimatore et al., 2021].

**ii) Thermodynamically,** the CP acts as a genomic Maxwell’s demon (MD) operator. In this role, the CP acquires information about the PES and uses this information to regulate genome-wide actuation through feedback. In Section 2.3, we further develop our previous information-thermodynamic framework and MD interpretation [Tsuchiya et al., 2025]. Information about the PES observed by the CP is defined by the net mutual information between the CP and PES, *I*_net_(CP;PES), which serves as the order parameter of CP-PES coupling [Tsuchiya et al., 2025]. Section 2.3.1 examines whether the temporal dynamics of *I*_net_(CP;PES) are phase synchronized with the CP entropy, *S*(CP), thereby testing CP-PES thermodynamic phase synchronization. Because CP and PES are distinct gene ensembles, their genes cannot be paired uniquely. This phase synchronization supports correspondence of their collective states at the same time points; expression-rank matching is then used to construct the CP-PES joint distribution and examine how *I*_net_(CP;PES) is decomposed into rank-matched pairwise and higher-order information contributions.

In Section 2.4, we examine how information gained by the CP is linked to CP EED work, connecting information acquisition to forward actuation across the critical transition. **Sections 2.4.4-4.5** connect the thermodynamic and dynamical roles of the CP by examining (1) how CP-PES information coupling is linked to CP-driven genome-engine actuation and (2) how their timing defines the progression from commitment to execution. Based on the four conditions, we propose four criteria for the irreversible arrow of cell-fate change in expression space. The conditional thermodynamic requirement can be tested through calibration of transcriptome-derived entropy to physical entropy and direct measurement of outward entropy export.

In the Discussion, **Section 3.1** examines the physical interpretation and scope of the genomic MD framework within SOC-controlled genome dynamics, and **Section 3.2** further considers chromatin and DNA structural dynamics as a possible physical basis for the transcriptome-derived critical transitions.

Together, ITA, CSA, and EFA provide a unified framework for identifying when genome-wide fate commitment occurs, distinguishing fate-guiding from non-fate-guiding responses, and defining experimentally testable time-gated transition windows. Phase-specific perturbation of CP-associated regulators would provide a direct test of CP-mediated causality.

All abbreviations are provided at the end of the manuscript.

## 2. Results

### 2.1 Whole-Expression Systems in Cultured Cell Lines

Cultured cells are considered open cellular systems that continuously exchange energy and matter with their culture environment via gas exchange (O_2_/CO_2_), heat flow, and net metabolite uptake and secretion in the culture medium. At the cell-population level, the whole-expression system is treated as an effective open genomic subsystem coupled to the cellular and culture environments. The cultures are maintained at a controlled temperature of (*T* = 37 °C = 310.15 K), while cellular metabolic heat is continuously dissipated to the temperature-controlled incubator, which functions as an external heat bath. CO_2_-regulated incubators and culture media impose relatively stable thermal and buffered extracellular conditions. Together with continuous metabolic conversion, nutrient and oxygen uptake, metabolite secretion, and heat dissipation, these conditions maintain the cells in a non-equilibrium state over the experimental window. These boundary conditions are consistent with nonequilibrium dissipative organization in open systems [Prigogine, 1967; De Bari et al., 2023]. This persistent non-equilibrium driving, imposed by the culture environment and mediated by cellular metabolism and signaling, reaches the nucleus as coordinated transcriptional and chromatin dynamics, motivating a genome-wide rather than gene-by-gene intranuclear description [Li, X. et al., 2018; Zimatore et al., 2021; Tsuchiya et al., 2025].

#### Coarse-Graining Coordinate Defining Macroscopic Variables

To expose genome-scale organization, we characterize each gene by its temporal fluctuation amplitude. For gene *i*, the root-mean-square fluctuation (*rmsf*) is

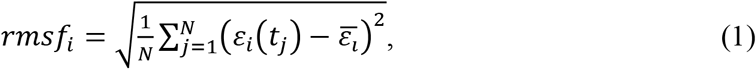

where *N* is the number of sampled time points and 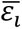 is the temporal average expression level of gene *i* over those time points. We then normalize this value by the maximum fluctuation observed in the dataset to obtain the normalized *rmsf*, *nrmsf*_i_ = (*rmsf*_i_/max(*rmsf*)). The final metric is the natural logarithm of this normalized value, *ln*(*nrmsf*), which is a time-independent gene-specific coordinate.

Because *nrmsf* measures the temporal variability of each gene, taking its logarithm transforms fluctuation magnitude into a scaling coordinate where power-law organization across the transcriptome emerges. We therefore adopt *ln*(*nrmsf*) as the self-organization coordinate of the whole-expression system (WES) [Tsuchiya et al., 2020; Zimatore et al., 2021]. Along the *ln*(*nrmsf*) coordinate, the WES is partitioned into three hierarchically coupled domains: the CP (critical point) region, the Super-domain (above the CP region along *ln*(*nrmsf*)), and the Sub-domain (below the CP region along *ln*(*nrmsf*)).

Coherent stochastic behavior (CSB) in grouped whole-expression dynamics identifies the CP region through bimodal singular critical behavior in the group-average expression changes of specific gene groups (see Fig. 2 in [Tsuchiya et al., 2025] and the technical definition in [Tsuchiya et al., 2024]). We define the peripheral expression system (PES) as PES = Super ∪ Sub, that is, all genes outside the CP (see **Table 1** for MCF-7 and HL-60 cancer cells).

**Table 1:** *ln(nrmsf*)-Based Domain Partitioning of Genome-Wide mRNA Expression in MCF-7 and HL-60 Cells under HRG, EGF, atRA, and DMSO Stimulation. Note: MCF-7 data include open reading frame (ORF) expression.

| Cell type | Sub-domain | CP region | Super-domain |
| --- | --- | --- | --- |
| HRG<br>MCF-7 cells<br>(22,277 mRNAs) | $\ln(nrmsf) < -2.6$<br>(7,917 mRNAs) | $-2.6 < \ln(nrmsf) < -2.4$<br>(3,846 mRNAs) | $-2.4 < \ln(nrmsf)$<br>(10,514 mRNAs) |
| EGF<br>MCF-7 cells<br>(22,277 mRNAs) | $\ln(nrmsf) < -2.7$<br>(7,425 mRNAs) | $-2.7 < \ln(nrmsf) < -2.5$<br>(4,033 mRNAs) | $-2.5 < \ln(nrmsf)$<br>(10,819 mRNAs) |
| atRA<br>HL-60 cells<br>(12,625 mRNAs) | $\ln(nrmsf) < -2.7$<br>(6,594 mRNAs) | $-2.7 < \ln(nrmsf) < -2.5$<br>(1,674 mRNAs) | $-2.5 < \ln(nrmsf)$<br>(4,357 mRNAs) |
| DMSO<br>HL-60 cells<br>(12,625 mRNAs) | $\ln(nrmsf) < -2.7$<br>(6,895 mRNAs) | $-2.7 < \ln(nrmsf) < -2.5$<br>(1,719 mRNAs) | $-2.5 < \ln(nrmsf)$<br>(4,011 mRNAs) |

**Note: Critical states**. In our previous work, we used super-, near-, and sub-critical states defined by gene-expression distributions [Tsuchiya et al., 2020]. Here we refer to the CP region simply as the CP unless domain-level distinctions are required.

This *ln*(*nrmsf*) coordinate is essential because it provides a time-independent genome-wide ordering axis for resolving both the thermodynamic organization (Section 2.3) and the effective expression-derived (EED) dynamics of the WES (Section 2.4). In both ITA and CSA, the *ln*(*nrmsf*) coordinate orders the CP, Super, and Sub domains. This coordinate enables quantification of CP-guided information coupling, entropy-related synchronization, and chromatin-state changes, thereby serving as an operational coarse-grained proxy coordinate for chromatin remodeling [Zimatore et al., 2021; Tsuchiya et al., 2025]. In the genome-engine framework, the same coordinate also defines the domain attractors and their flux-balance network, thereby providing the coordinate basis for evaluating domain-attractor EED work in expression space.

Thus, *ln*(*nrmsf*) is not merely the SOC metric, but also the essential coarse-grained coordinate that defines the macroscopic variables required to unify information thermodynamics with dynamical systems analysis in cell-fate control (**Sections 2.3-2.4**).

Along this common coordinate, system-wide coupling generates extended correlations across the WES. The CP region defines the critical interface at which homeostatic stability coexists with genome-wide transition susceptibility manifested as power-law criticality. Nonlinear higher-order mutual information between the CP and PES quantifies the coupling that supports this critical dynamic balance. At this interface, the CP drives the genome-wide transition through extended correlations and Maxwell’s demon-guided rewriting of chromatin memory, enabling coherent chromatin reorganization (**Sections 2.3-2.4**).

### 2.2 Coherent Stochastic Behavior (CSB): Statistical Validation of Convergence

CSB manifests as the convergence of unit-mass center-of-mass (CM) dynamics of a gene group defined along a coordinate such as *ln*(*nrmsf*) in two complementary ways:

- **Horizontal convergence**: Within a fixed group, the CM of a randomly sampled subset converges to the CM of the full group as sample size increases. Thus, by the law of large numbers, larger random subsets yield increasingly reproducible CM dynamics (e.g., Fig. 10 in [Tsuchiya et al., 2016]; Fig. 3 in [Tsuchiya et al., 2020]). Coordinate-based grouping further reveals CSB (e.g., **Figure 1**; Figs. 5-6 in [Tsuchiya et al., 2009]; Fig. 1A,D in [Tsuchiya et al., 2024]).
- **Vertical convergence**: For a fixed, large sample size, repeated random sampling gives a distribution of CM estimates. As the number of bootstrap resamples increases, the bootstrap ensemble average of the resamples approaches the group CM dynamics, and the remaining bootstrap-to-bootstrap variation provides a practical Monte Carlo uncertainty scale for the CM estimator (e.g., Fig. 1C in [Tsuchiya et al., 2024]; Fig. 4 in [Tsuchiya et al., 2025]).

**Figure 3.**
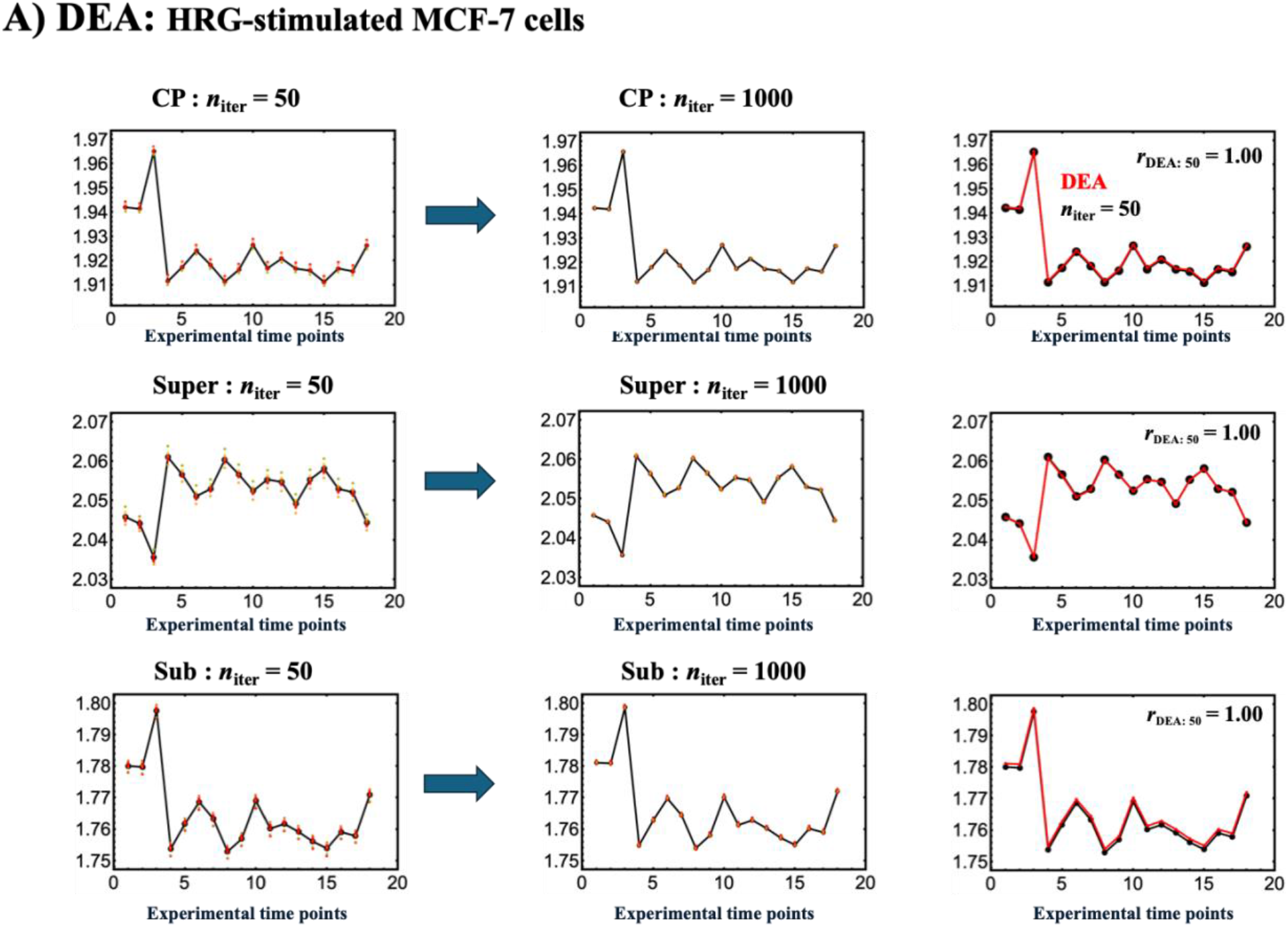
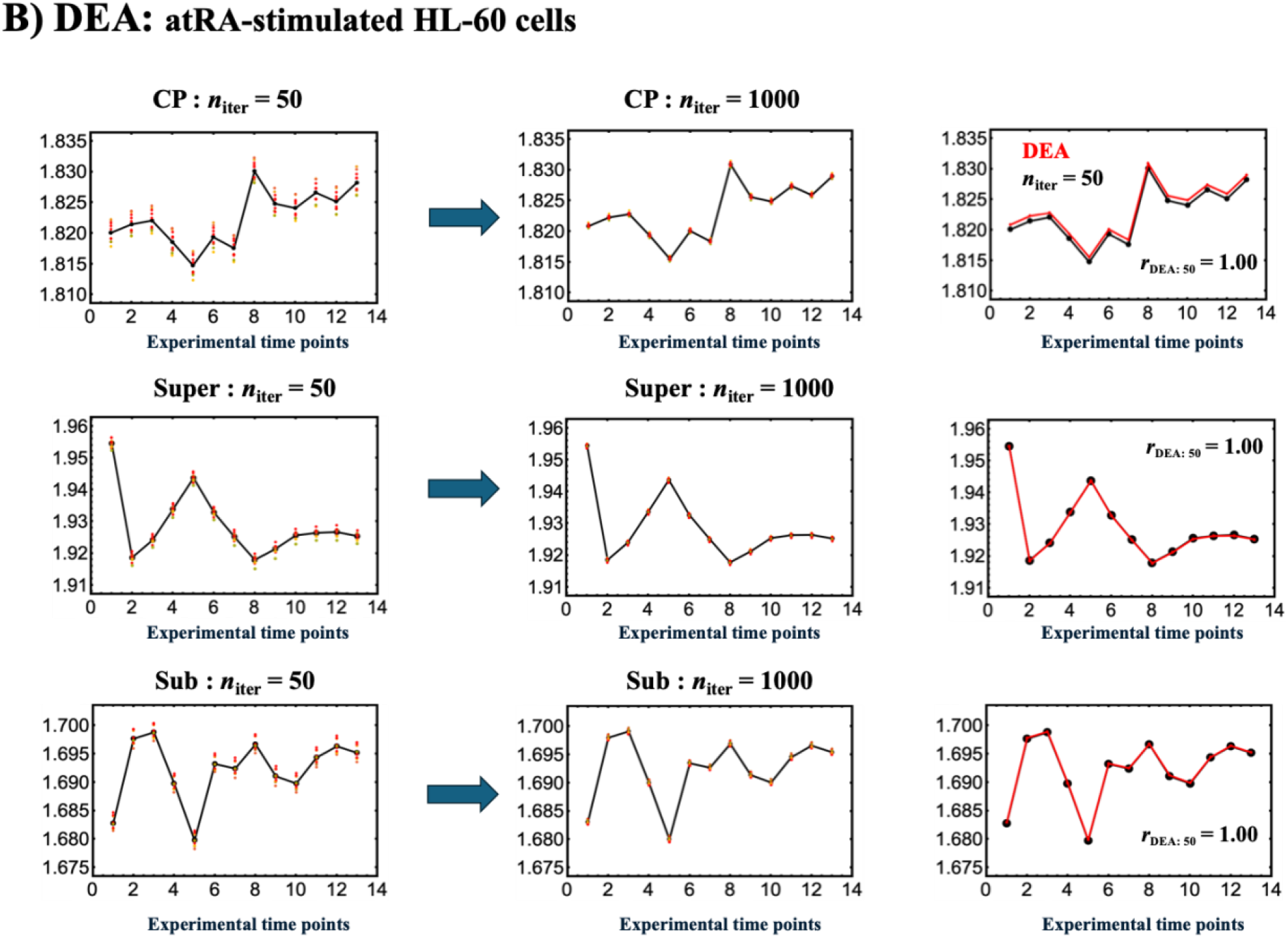
Statistical validation of coherent stochastic behavior (CSB) by bootstrap convergence of domain ensemble-average (DEA) trajectories. **(A)** HRG-MCF-7 cells and **(B)** atRA-HL-60 cells. Rows correspond to the CP, Super-domain, and Sub-domain trajectories, respectively. **Left panels**: the bootstrap ensemble-average trajectory (CM with unit mass; black line) computed from *n*_boot_ = 10 independent bootstrap runs at *n*_iter_ = 50, with the corresponding bootstrap scatter (red points) at each time point. **Middle panels**: the same display at *n*_iter_ = 1000, showing reduced bootstrap scatter while retaining the stabilized ensemble-average trajectories. **Right panels**: convergence of the bootstrap ensemble-average trajectory to the corresponding domain ensemble-average trajectory (domain attractor) is already evident at *n*_iter_ = 50. This is quantified by a temporal Pearson correlation of approximately unity ( *r*_DEA:50_ ≈ 1.0), despite the larger bootstrap scatter at *n*_iter_ = 50. This convergence supports the use of domain ensemble-average trajectories as stable low-dimensional state variables for expression-flux analysis (EFA).

**Figure 4.**
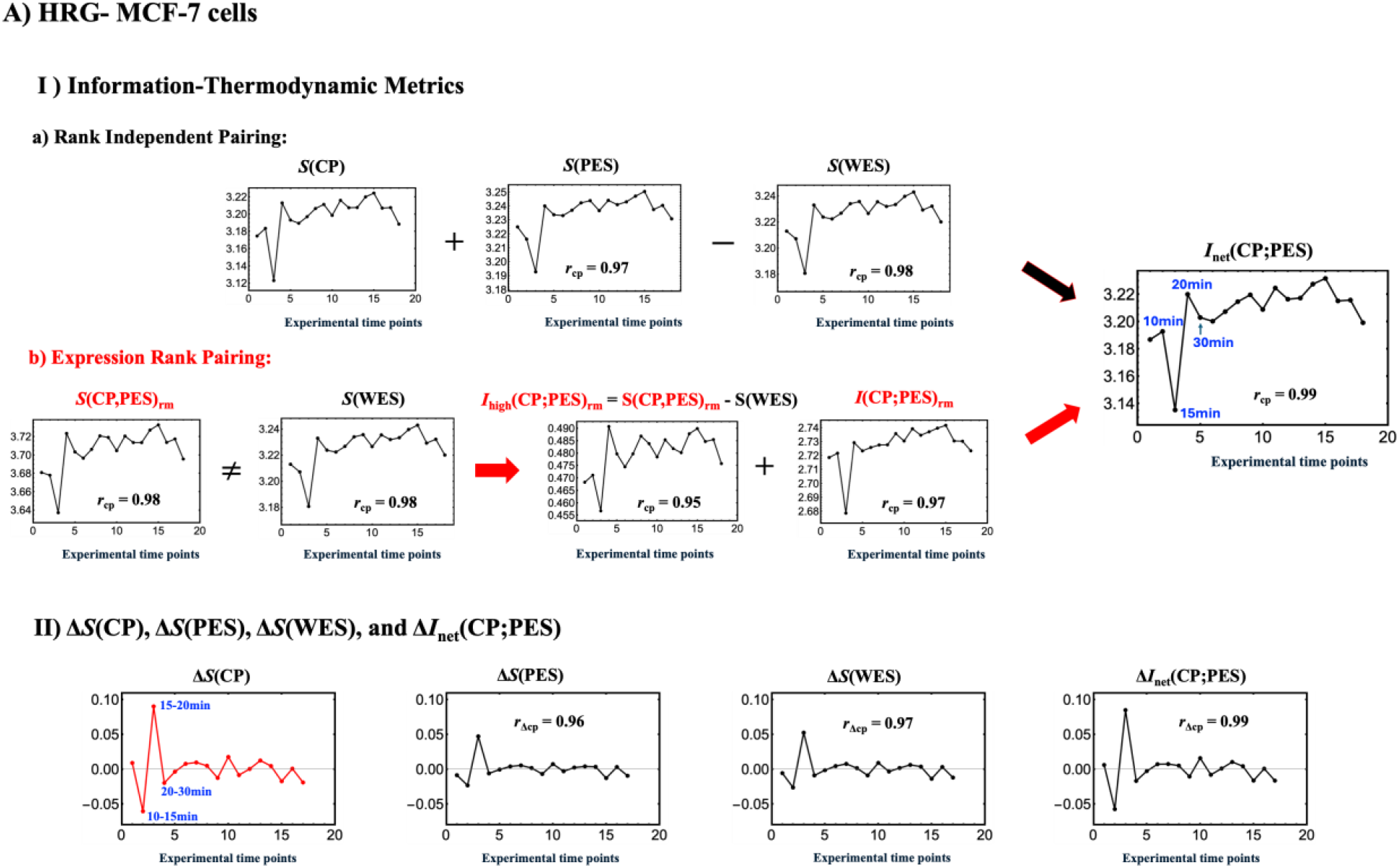
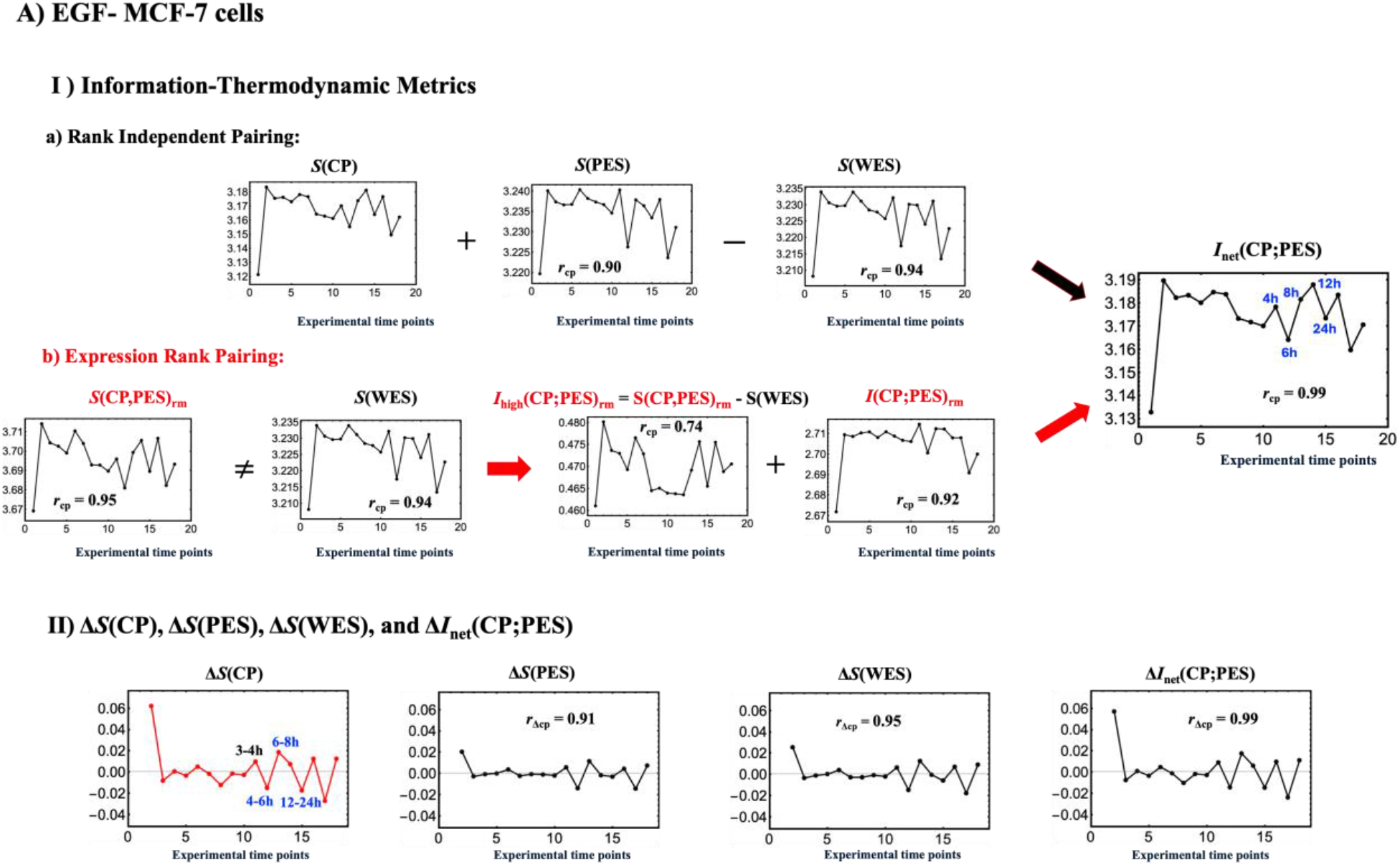
Information-thermodynamic comparison of fate-guiding and non-fate trajectories in MCF-7 cells. **(A)** HRG (ErbB3/ErbB4 ligand; differentiation) and **(B)** EGF (ErbB1/EGFR ligand; proliferation). Blue text indicates the two-time-point windows of the Maxwell’s demon (MD) cycle, ordered by phase (Preparation, Measurement, and Feedback; Table 3). Temporal Pearson correlations with CP entropy, *S*(CP), and CP entropy change, Δ*S*(CP), are denoted by *r*_CP_ and *r*_ΔCP_, respectively.

**Figure 5.**
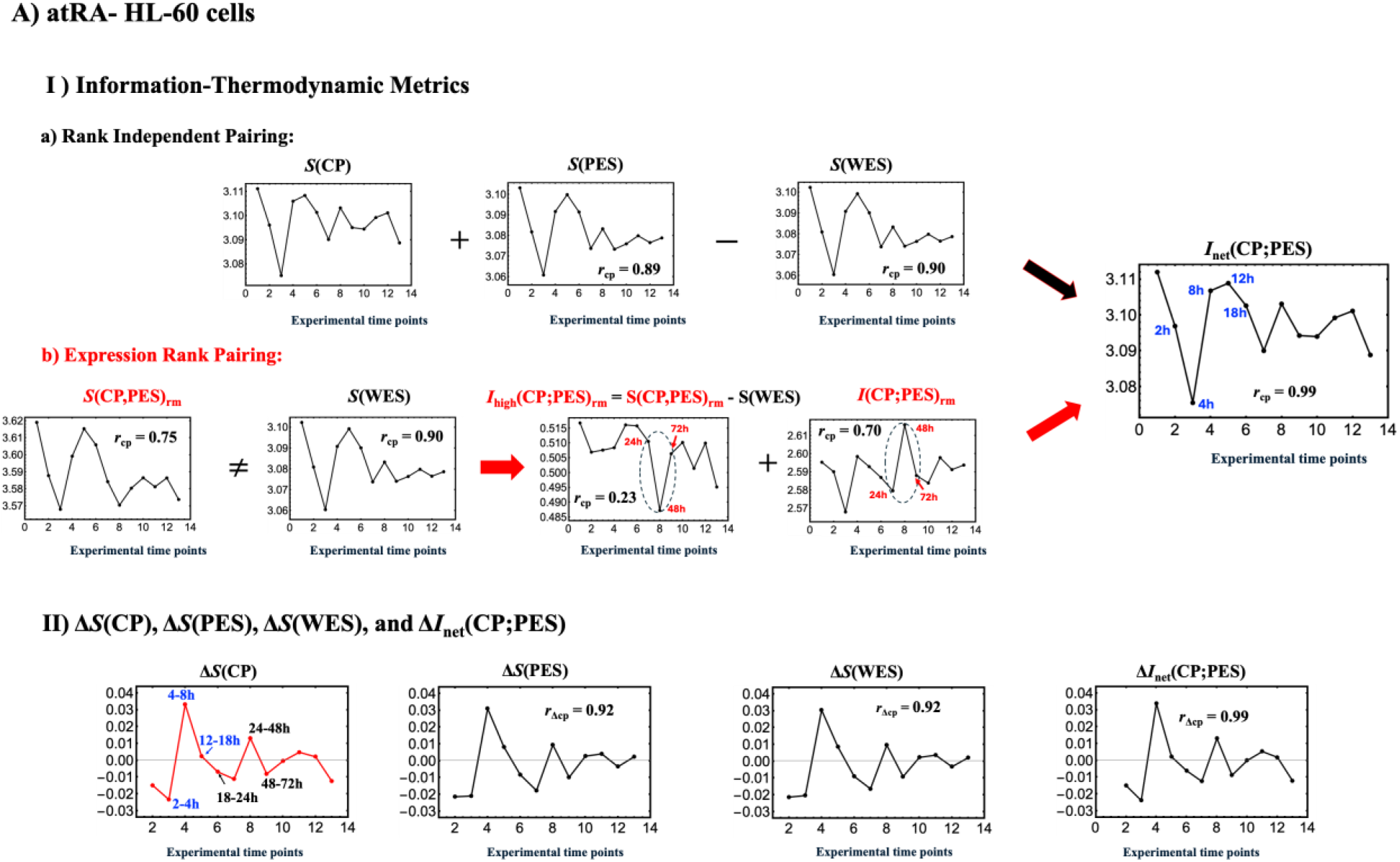
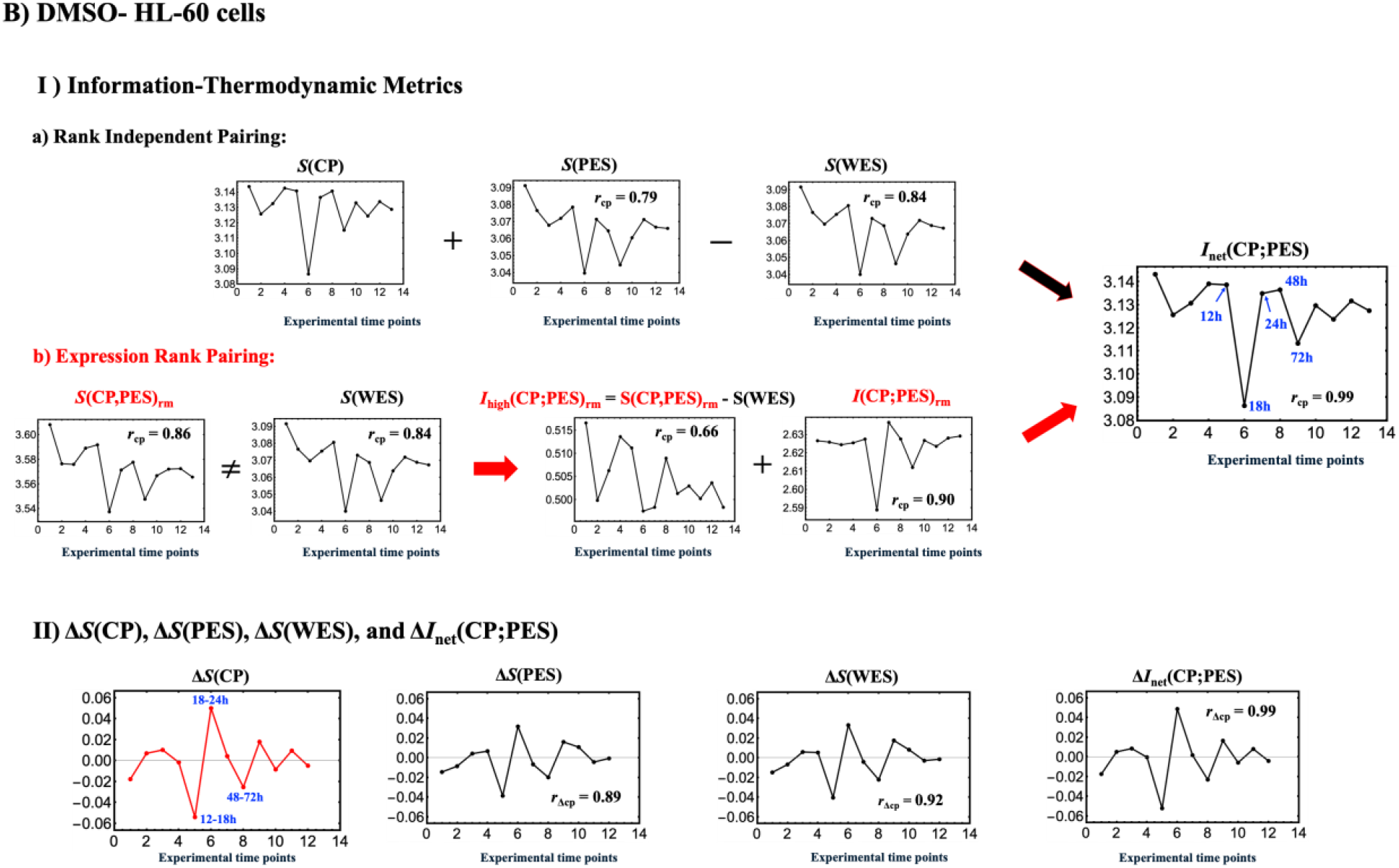
Information-thermodynamic comparison of fate-guiding trajectories in HL-60 cells. **(A)** atRA (nuclear-receptor ligand) and **(B)** DMSO (polar-solvent inducer). Blue text, MD-cycle phase ordering, and temporal Pearson correlation labels are as defined in **Figure 4**.

**Figure 6.**
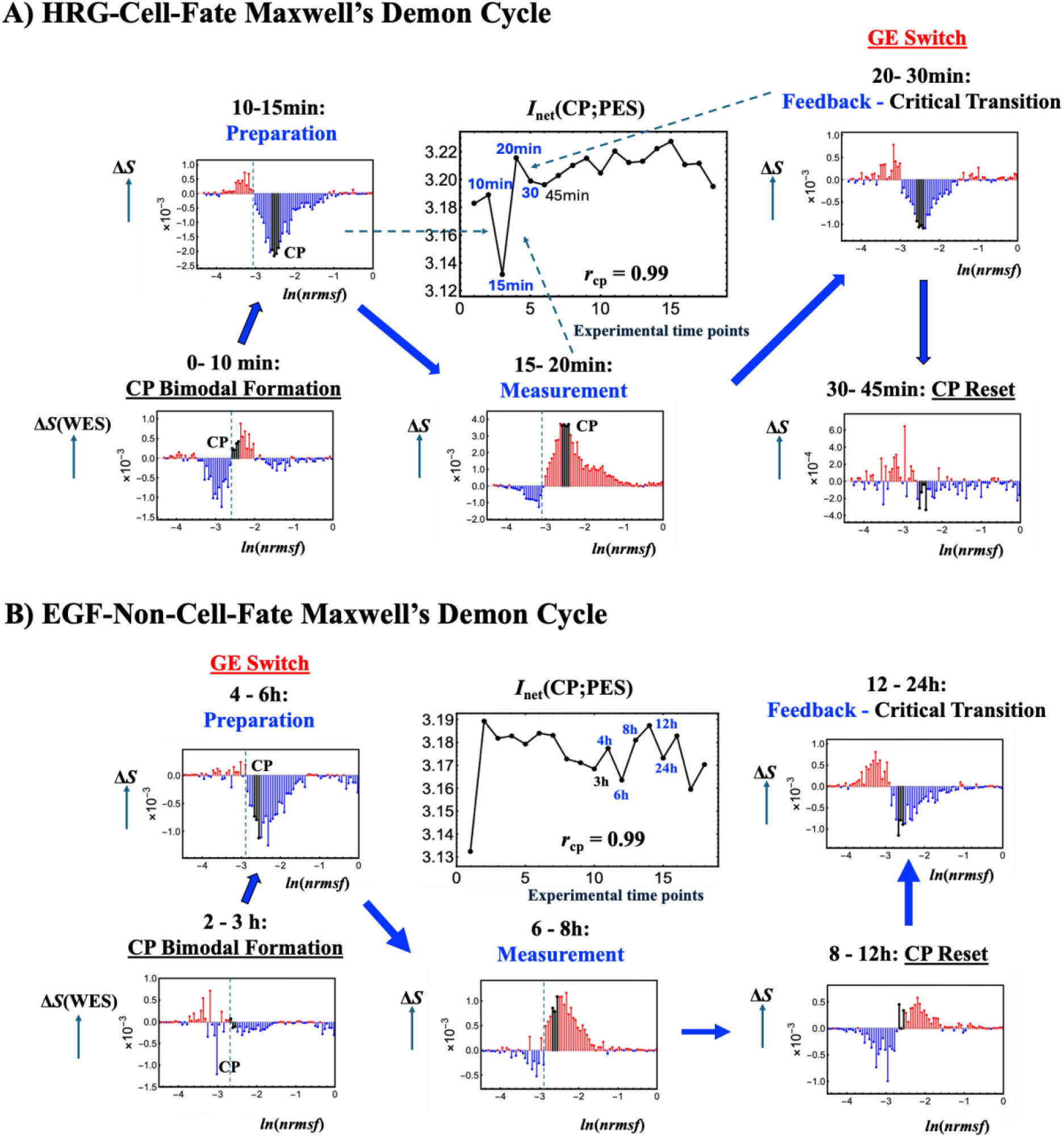
Stimulus-dependent MD-cycle timing in MCF-7 cells: HRG (fate) versus EGF (non-fate). Panels A and B display the temporal trajectory of *I*_net_(CP;PES) in the center, with the surrounding plots showing CSA-derived, bin-resolved entropy increments of the whole expression system, Δ*S*(WES), plotted along the *ln*(*nrmsf*) coordinate. *ln*(*nrmsf*) serves as a self-organized proxy for chromatin remodeling dynamics. Black lines indicate the CP region. The x-axis shows the *ln*(*nrmsf*) coordinate discretized into 80 fixed bins (logical chromatin states), and the *y*-axis reports the corresponding Δ*S*(WES) in each bin. In CSA, coherent positive Δ*S*(WES) > 0 is consistent with coherent chromatin opening (red lines), whereas coherent negative Δ*S*(WES) < 0 indicates increased ordering of the expression distribution (blue lines). The Δ*S*(WES) profiles for both HRG and EGF conditions exhibit a CP-driven coherent ON/OFF boundary near *ln*(*nrmsf*) values between −3.15 and −3.00 in HRG and between −2.96 and −2.85 in EGF, indicating the presence of a CP-driven Maxwell’s demon (MD) double-well potential profile (see the main text for CP-driven bistable switching). The central panel shows that *I*_net_(CP;PES) remains tightly synchronized with *S*(CP) (temporal Pearson correlation: *r*_cp_ = 0.99) in both HRG and EGF. Blue labels mark the MD boundary times in the *I*_net_(CP;PES) panel. The genome engine (GE) switch, a dynamical marker associated with the Maxwell’s demon cycle (Section 2.4.5; Table 3), is shown by underlined red labels. With the GE-switch timing included, **(A) HRG (cell-fate change)** shows 1) CP-bimodal formation before the labeled Preparation window and CP reset at 30-45 min after the Feedback phase, reflecting formation and re-initialization of CP-driven bistable switching; and 2) a clear three-phase sequence of Preparation (10-15 min), Measurement (15-20 min), and Feedback (20-30 min), with the GE switch occurring in this Feedback interval. **(B) EGF (non-cell-fate change)** shows 1) CP-bimodal formation before the labeled Preparation window and CP reset after the Measurement phase, reflecting an early re-initialization route before Feedback; and 2) an early GE switch at 4-6 h in the Preparation phase, before the main Measurement window (6-8 h). During Measurement, the Δ*I*_net_ gain is about fivefold smaller than in HRG (**Figure 4**), indicating insufficient information-drive accumulation to support CP-guided fate control, followed by CP reorganization (8-12 h) and subsequent Feedback (12-24 h), consistent with incomplete stabilization of a new state.

In this section, using the vertical bootstrap convergence principle, we evaluate the statistical convergence of the information-thermodynamic metrics in ITA [Tsuchiya et al., 2025] through repeated Monte Carlo resampling of fixed-size gene sets and ensemble averaging of the resulting time-resolved trajectories, and statistically validate coherent stochastic behavior (CSB) as the foundation for the genome engine dynamical system in EFA [Tsuchiya et al., 2020] and chromatin state analysis (CSA) [Tsuchiya et al., 2025].

**(i) Information-Thermodynamic Analysis (ITA): Bootstrap convergence of IT metric trajectories**. We statistically validate convergence of information-thermodynamic (IT) metrics using the vertical CSB bootstrap principle. MCF-7 cells under HRG and EGF conditions and HL-60 cells under atRA and DMSO conditions all show clear convergence behavior (HRG, Fig. 4 in [Tsuchiya et al., 2025]; atRA and DMSO, **Figure 2**; EGF, not shown). We compute time-resolved trajectories with *n*_boot_ = 10 independent bootstrap runs while increasing the resampling iterations per bootstrap run, *n*_iter_ = 50-1000.

Here, *n*_iter_ denotes the number of resampling iterations performed within one bootstrap run. In each resampling iteration, we randomly sample a fixed-size gene set (1000 genes) from the target domain without replacement and compute the IT metric trajectory across all time points. For the joint metrics *S*(CP,PES) and *I*(CP;PES), sampled CP and PES expression values were independently ranked and paired at equal relative ranks.This procedure enables the construction of a rank-matched joint distribution between the two gene ensembles without requiring gene-to-gene correspondence (see Section 2.3.1 for details). The bootstrap-run trajectory is then obtained by averaging these *n*_iter_ iteration-level trajectories at each time point. Increasing *n*_iter_ reduces resampling noise, which is quantified by the bootstrap standard deviation (SD) across independent bootstrap runs at each time point, and stabilizes the bootstrap ensemble-average trajectory. The bootstrap SD at time point *t* is computed as

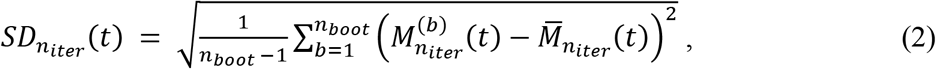

Where 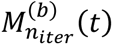 is the IT metric value from bootstrap run *b* and 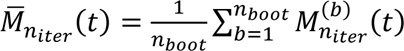 is the bootstrap ensemble average across bootstrap runs at time point *t*. Both the temporal average of the metric level and the temporal average of the bootstrap SD show systematic stabilization as *n*_iter_ increases from 50 to 1000 in both HRG-MCF-7 and atRA-HL-60 cells (see **Figure 2** for temporal profiles and **Table 2** for bootstrap convergence).

**Table 2.** Bootstrap convergence summary for IT metrics (HRG-MCF-7 cells and atRA-HL-60 cells). Each entry reports the temporal average of the bootstrap ensemble mean ± the temporal average of the across-run SD, calculated from *n*_boot_ = 10 independent bootstrap runs. Results are shown for two settings of the resampling iterations per bootstrap run, *n*_iter_ = 50 and 1000, using a fixed resample size of 1000 genes in each iteration (cf. **Figure 2** for time-resolved trajectories and bootstrap-scatter convergence in atRA-HL-60 cells, and Fig. 4 in [Tsuchiya et al., 2025] for the corresponding results in HRG-MCF-7 cells). To examine convergence under controlled and reproducible sampling conditions, the random-number seed was changed for each of the 10 bootstrap runs, and within each run, the *n*_iter_ = 50 result corresponds to the first 50 iterations of the same random stream used through *n*_iter_ = 1000. This enables direct comparison between the two settings while preserving reproducibility and distinct random streams across runs.

| Cell Type | Metric | $n_{\text{iter}}=50$ | $n_{\text{iter}}=1000$ |
| --- | --- | --- | --- |
| HRG<br>MCF-7 Cells | $S(\text{CP})$ | $3.1986 \pm 0.0014$ | $3.1980 \pm 0.0004$ |
| HRG<br>MCF-7 Cells | $S(\text{WES})$ | $3.2254 \pm 0.0013$ | $3.2257 \pm 0.0004$ |
| HRG<br>MCF-7 Cells | $S(\text{CP}, \text{PES})$ | $3.7067 \pm 0.0058$ | $3.7057 \pm 0.0011$ |
| HRG<br>MCF-7 Cells | $I(\text{CP}; \text{PES})$ | $2.7270 \pm 0.0060$ | $2.7275 \pm 0.0012$ |
| atRA<br>HL-60 Cells | $S(\text{CP})$ | $3.0980 \pm 0.0015$ | $3.0976 \pm 0.0003$ |
| atRA<br>HL-60 Cells | $S(\text{WES})$ | $3.0821 \pm 0.0019$ | $3.0820 \pm 0.0006$ |
| atRA<br>HL-60 Cells | $S(\text{CP}, \text{PES})$ | $3.5891 \pm 0.0078$ | $3.5890 \pm 0.0013$ |
| atRA<br>HL-60 Cells | $I(\text{CP}; \text{PES})$ | $2.5913 \pm 0.0065$ | $2.5908 \pm 0.0014$ |

We use the ensemble-average trajectory at *n*_iter_ = 1000 as the practical baseline trajectory for convergence comparisons. Convergence is validated by two complementary numerical checks:

- **SD convergence**: The time-averaged SD decreases and stabilizes with increasing *n*_iter_. At each experimental time point, the bootstrap scatter, defined as the SD across *n*_boot_ = 10 runs, systematically contracts, with the characteristic scale decreasing from *O*(10^-3^) at *n*_iter_ = 50 to *O*(10^-4^) at *n*_iter_ = 1000 for *S*(CP) and *S*(WES). The metrics *S*(CP, PES) and *I*(CP; PES) converge more slowly because they are estimated from a two-dimensional contingency table.
- **Ensemble-average stabilization**: The ensemble-average trajectories of the information-thermodynamic metrics are already effectively invariant at *n*_iter_ = 50 (**Figure 2**). Under these criteria, the IT metric trajectories are reproducible within the bootstrap noise level, demonstrating convergence. Each information-thermodynamic metric trajectory represents an ensemble-average quantity.

**(ii) Statistical Validation of Coherent Stochastic Behavior (CSB)**. We statistically validate CSB by demonstrating convergence to the domain ensemble-average (DEA) trajectories for the CP region, the Super-domain, and the Sub-domain in both HRG-MCF-7 and atRA-HL-60 cells using the same bootstrap procedure as ITA (above).

**Figure 3** demonstrates rapid convergence with increasing *n*_iter_, consistent with the law of large numbers and verified by two complementary checks:

- **SD convergence**: The time-averaged SD decreases from *O*(10^-3^) at *n*_iter_ = 50 to *O*(10^-4^) at *n*_iter_ = 1000 and stabilizes with increasing *n*_iter_.
- **Ensemble-average stabilization**: The ensemble-average trajectory is already effectively invariant at *n*_iter_ = 50. This is supported by the temporal Pearson correlation (*r*_DEA:50_ ≈1.0) between the domain ensemble-average trajectory (DEA) and the bootstrap ensemble-average trajectory at *n*_iter_ = 50, despite the larger bootstrap scatter (**Figure 3**).

This stabilization demonstrates that the unit-mass center-of-mass (CM) embodies a robust expression-level law of large numbers. Consequently, each bootstrap domain ensemble-average trajectory converges to the DEA trajectory, which is a reproducible domain-attractor state variable. This convergence also supports modeling CM dynamics with unit mass (*m* = 1), i.e., treating each gene-expression contribution as an equal unit-mass element. Thus, the ensemble-average dynamics define domain effective forces. As a result, high-dimensional genome-wide expression dynamics across embryonic and cancer cell development collapse onto a one-dimensional trajectory along the domain coordinate, enabling expression flux analysis (EFA) [Tsuchiya et al., 2016, 2020, 2024]. This collapse provides the foundation for unifying genome engine dynamics with open-system, information-thermodynamic genome regulation (Section 2.4).

**(iii) Reconstructive Bootstrap for Chromatin State Analysis (CSA).** CSA reconstructs the gene-expression probability distribution at each time point using the bootstrap-converged entropy obtained in (i). CSA quantifies coherent chromatin state transitions by mapping stochastic gene-expression fluctuations onto a time-independent *ln*(*nrmsf*) coordinate. This coordinate serves as a coarse-grained proxy for chromatin remodeling [Zimatore et al., 2021; Krigerts et al., 2021; Erenpreisa et al., 2023; Tsuchiya et al., 2025] (see Section 2.3.2 for details). We separate a physical description from a logical description: the physical state at time *t*_j_ is the reconstructed gene-expression probability distribution, specifying the microstate statistics of stochastic expression; the logical state is the coarse-grained chromatin macrostate indexed by fixed *ln*(*nrmsf*) bins.

Using the bootstrap-converged entropy obtained in (i), we reconstruct the gene-expression distribution at each time point and normalize it so that its binned probabilities sum to one. Genes are then grouped into fixed *ln*(*nrmsf*) bins. For each reconstructed gene-expression distribution, we calculate the gene-level entropy contributions and map them onto the fixed *ln*(*nrmsf*) bins. This reconstruction yields stable, time-resolved chromatin state trajectories along *ln*(*nrmsf*).

Thus, CSA converts each reconstructed gene-expression distribution at *t*_j_ into a time-resolved entropy, S(WES)(*t_j_*) = − ∑*_k_ p_k_*(*t_j_*)*ln* (*p_k_*(*t_j_*)), where *p_k_*(*t_j_*) is the reconstructed probability in bin *k* with ∑*_k_ p_k_*(*t_j_*) = 1. The method maps the interval change Δ*S*(WES)[*t*_j_ → *t*_j+1_] onto the fixed *ln*(*nrmsf*) coordinate to visualize coherent chromatin state transitions across the Maxwell’s demon cycle. Within this *ln*(*nrmsf*)-based chromatin-remodeling proxy (Section 2.3.2), Δ*S*(WES) > 0 is consistent with chromatin unfolding, whereas Δ*S*(WES) < 0 indicates increased ordering of the expression distribution; its structural correlate is condition- and phase-dependent. This two-level structure follows information thermodynamics, in which microstates are described by probability distributions and logical states are coarse-grained macrostates used to track chromatin state changes [Maroney, 2009].

### 2.3 Genomic Maxwell’s Demon (MD) Cycle on MCF-7 Cells and HL-60 Cells

In this section, higher-order nonlinear interactions between the CP and PES are elucidated using the ITA bootstrap-convergence approach. The CP is then shown to act as a Maxwell’s demon guiding rewritable chromatin memory, thereby establishing a generalized MD cycle across the four stimuli.

#### 2.3.1 Thermodynamic Phase Synchronization as a Basis for CP-PES Pairing

Given that the CP and PES genes are distinct, CP-PES pairing is not unique; the two-dimensional probability distribution depends on how the CP and PES expression values are paired [Sklar, 1959; Größer and Okhrin, 2022]. Therefore, we address how these expression values can be paired to construct the CP-PES joint distribution.The convergence-based approach in Section 2.2 statistically validates the IT metrics; varying the histogram bin number produced only time-independent constant offsets, while preserving the temporal dynamics (not shown). The results show that the entropy of the CP, *S*(CP), is temporally phase synchronized with *S*(PES) and *S*(WES) at corresponding time points. Furthermore, net mutual information, defined as

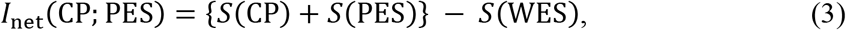

is also temporally phase synchronized with *S*(CP) (**Figures 4-5**).

These pairing-independent phase synchronizations establish that the CP and PES evolve as coordinated collective ensembles at the sampled time points (i.e., no observable lag), thereby supporting correspondence of their collective states at the same time point. Furthermore, EFA independently supports CP-PES synchronization through phase-synchronized expression-derived dynamical work of the CP and PES (Section 2.4.4).

Under coherent stochastic behavior, CP and PES are treated as collective gene ensembles rather than as individually matched genes. Because their expression values are defined on the same centered coordinate relative to the genome attractor (GA), their collective states can be matched by equal relative rank in the same direction. We therefore use expression rank matched (rm) to construct the joint distribution for *S*(CP,PES)_rm_ and the corresponding pairwise information term, *I*(CP,PES)_rm_, without assuming gene-to-gene correspondence.

We independently estimate the whole expression entropy, *S*(WES), from the WES probability distribution, and the rank-matched joint entropy, *S*(CP,PES)_rm_, from the two-dimensional CP-PES probability distribution constructed by expression-rank matching. **Figures 4**-**5** show that these two independently evaluated entropies do not coincide:

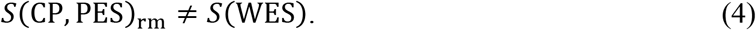

Here, the subscript rm stands for rank matching used to construct the two-dimensional CP-PES joint probability distribution. The persistent nonzero difference is robust to the finite-sampling procedure used in the ITA estimation (see Section 2.2) and therefore motivates an operational decomposition of system-level information into a rank-matched pairwise term and a higher-order contribution.

The rank-matched pairwise information term is defined as

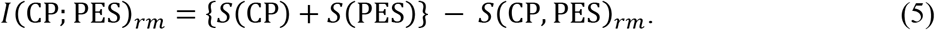

This quantity is not strictly a Shannon mutual information estimated from a two-dimensional joint probability table because *S*(CP) is calculated over the native CP expression range, whereas the CP marginal of the joint table is defined over the broader PES range.

**Note: Rank-Matched Decomposition**. Within each bootstrap iteration as described in Section 2.2, the same sampled CP and PES gene sets are used for the corresponding entropy estimates, and the sampled WES set is their union. Importantly, the rank-matched decomposition exactly reproduces the pairing-independent *I*_net_(CP,PES) at every time point: *I*_net_(CP,PES) = *I*(CP,PES)_rm_ + *I*_high_(CP,PES)_rm_ = {S(CP) + S(PES)} – S(WES) as given by Eq. 3 and illustrated in **Figures 4-5**. Thus, the rank-matched decomposition preserves the pairing-independent ITA quantity in Eq. (3), which is defined from Shannon entropies within the stochastic information-thermodynamic framework [Seifert, 2012; Van den Broeck and Esposito, 2015; Parrondo et al., 2015].

The higher-order contribution is defined as the difference between *I*_net_(CP;PES) and *I*(CP;PES)_rm_:

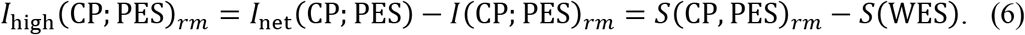

#### Both MCF-7 and HL-60 transcriptome data (**Figures 4**-**5)** show that

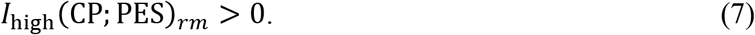

Thus, *I*_high_ > 0 directly quantifies, in expression space, a higher-order CP-PES information contribution not captured by the rank-matched pairwise term, *I*(CP,PES)_rm_. CP-GA dynamical phase synchronization (Section 2.4.2) indicates coupling beyond the rank-matched CP-PES pair, because the GA represents the collective state of the WES.

##### 2.3.2. Transcriptome-Derived Proxy versus Direct Structural Evidence

*ln*(*nrmsf*) is the natural logarithm of the normalized root mean square fluctuation defined in Eq. 1. This coordinate and the associated entropy dynamics are coarse-grained transcriptomic quantities derived from time-series transcriptome data. Because transcriptomic states provide a coarse-grained representation that integrates over detailed chromatin and DNA configurations, these quantities characterize collective structural organization rather than microscopic detail. Chromatin remodeling changes transcriptional access to regulatory regions and can thereby reshape gene-expression distributions, providing a mechanistic basis for changes in transcriptomic entropy.

Our previous study [Zimatore et al. (2021)] statistically validated the role of chromatin remodeling as the material basis of SOC control of genome expression. This validation was based on recurrence quantification analysis (RQA) and principal component analysis (PCA) of gene-expression data,together with the temporal correspondence between expression fluctuations and PAD structural reorganization [Krigerts et al., 2021]. RQA showed a robust exponential decay of gene co-regulation with increasing separation in chromosomal gene order, consistent with local chromatin folding and flexibility. PCA showed that *ln*(*nrmsf*) quantitatively captures the major temporal gene-expression fluctuations around the genome attractor (GA), including the singular behavior of the CP in *ln*(*nrmsf*) space. The temporal correspondence of these expression fluctuations with PAD structural reorganization further links this expression-space organization to chromatin structural dynamics. Thus, the *ln*(*nrmsf*) coordinate provides a quantitative expression-derived proxy for chromatin remodeling at the coarse-grained level of collective genome-expression dynamics.

In HRG-stimulated MCF-7 cells, Krigerts et al. (2021) experimentally observed PAD unraveling together with unfolding of H3K4me3-positive chromatin around the split PADs at 15 min, together with an acridine orange (AO) fluorescence signal indicating a more open DNA conformation at 20 min. They further related these structural changes to the first critical transcription avalanche reported by Tsuchiya et al. (2015, 2016). These experimentally observed structural changes temporally coincide with the critical transition, identified by ITA and CSA, as shown in Section 2.4.5, supporting the chromatin-remodeling interpretation of the transcriptome-derived transition in HRG.

Within this structural interpretation, Δ*S*(WES) > 0 is proposed as a coarse-grained transcriptomic signature consistent with chromatin unfolding. Negative Δ*S*(WES) indicates increased ordering of the transcriptome-derived expression distribution, whereas its chromatin structural correlate depends on the condition and dynamical phase, particularly during critical transitions (see Section 2.3.3), and cannot be assigned from the entropy sign alone. Structural interpretation therefore requires integration with the corresponding transition dynamics and available structural measurements. For DMSO, EGF, and atRA, condition-specific structural validation remains to be established.

Thus, the analysis establishes collective transcriptomic transitions supported by independent structural evidence in HRG. The physical interpretation of these findings, their limitations, and their connections to chromatin dynamics and EED work are discussed in **Sections 3.1 and 3.2**.

##### 2.3.3 Generalized CP-Driven Genomic Maxwell’s Demon (MD) Cycle

The net MI *I*_net_(CP; PES), including higher-order terms, serves as an order parameter of CP-PES coupling and resolves the three characteristic operational phases of a genomic Maxwell’s demon (MD), with the CP functioning as the MD operator [Tsuchiya et al., 2025]. Here, we generalize the MD cycle to reveal a common operational structure across the four conditions. By analyzing time-dependent changes in stochastic information-thermodynamic metrics, we identify when and how thermodynamic constraints shift (see Section 2.4.5 for details), thereby quantitatively resolving the organization, stability, and critical transitions within the genomic Maxwell’s demon cycle:

- **Thermodynamic phase synchronization between the CP and PES.** Across all four trajectories, *I*_net_(CP;PES) and Δ*I*_net_(CP;PES) are tightly synchronized with *S*(CP) and Δ*S*(CP), respectively (*r*_CP_ = *r*_ΔCP_ = 0.99; **Figures 4-5**), demonstrating strong CP-PES coupling in both fate-guiding and non-fate responses. As the order parameter of CP-PES coupling, *I*_net_ synchronization with *S*(CP), together with Δ*S*(WES) ∝ Δ*S*(CP), identifies the CP as the information-thermodynamic controller of the WES. This thermodynamic phase synchronization underlies the coupling between Δ*I*_net_(CP;PES) and incoming CP work in expression space described in Section 2.4.4.
- **Emergent genome-level Maxwell’s demon cycle**. Under CP-guided SOC, local enzymatic Maxwell’s demon actions [Ichii et al., 2026] are integrated along the *ln*(*nrmsf*) coordinate into a coherent genome-wide non-equilibrium MD mechanism. Figures 6–7 show that the MD cycle comprises Preparation, Measurement (active sensing), and Feedback, during which condition-specific genome-wide reorganization occurs, and Table 3 summarizes the timing of these phases across conditions. **Figures 6-7** show, across all conditions, the timing of the genome-engine (GE) switch [Tsuchiya et al., 2016, 2020, 2024], which is defined by a topological change in the genome-engine interaction network. The GE switch serves as a condition-specific temporal landmark within or after the MD cycle, as detailed in Section 2.4.5.
- **Formation of CP-driven bistable chromatin switching**. The convergence-validated reconstructive CSA established in Section 2.2 resolves coherent chromatin-state transitions as Δ*S*(WES) profiles along the time-independent *ln*(*nrmsf*) coordinate. As discussed above, positive and negative domains correspond to chromatin opening and genome-wide ordering, respectively (**Figures 6, 7B**). In HRG, EGF, and DMSO, the Δ*S*(WES) profile exhibits switchable bistable chromatin states with a coherent ON/OFF chromatin-state boundary near *ln*(*nrmsf*) values between −3.15 and −3.00 in HRG and DMSO, and between −2.96 and −2.85 in EGF. These bistable states are organized by a CP-driven MD double-well potential, suggesting CP-driven bistable switching of the chromatin-state landscape. Through this switching, the CP organizes CP-PES information into ON/OFF chromatin states, consistent with bistable-switch principles in cell-fate and cell-cycle control [Xiong and Ferrell, 2003; Stallaert et al., 2019]. In contrast, in atRA, the negative Δ*S*(WES) profile shows an all-or-none compaction pattern across the *ln*(*nrmsf*) coordinate under CP-PES phase synchronization (**Figure 7A**). This pattern does not form the CP-driven MD double-well profile required for coherent bistable switching.
- **CP-enhancing and CP-braking modes of CP-driven chromatin-state regulation**. **Figures 6, 7B** show that, in HRG, EGF, and DMSO, transient CP bimodality appears along the *ln*(*nrmsf*) coordinate as CP-bimodal formation and CP reset. CP-bimodal formation occurs before the MD cycle, whereas CP reset occurs after the Measurement phase in EGF and DMSO, and after the Feedback phase in HRG. CP reset denotes reconfiguration and re-initialization of the CP bimodal state, not its disappearance. This transient sequence marks the formation or re-initialization of the CP-driven MD double-well potential profile, revealing the CP-enhancing mode. In contrast, sustained CP bimodality in atRA prevents coherent ON/OFF chromatin-state switching. This suppresses net-driving CP actuation during the MD cycle and delays fate commitment until after the MD cycle, revealing the CP-braking mode (**Figure 7A**; see Section 2.4.5 for details).

**Table 3.** Timing-window summary of the genomic Maxwell’s demon (MD) cycle, genome-engine switch, and CP EED work across stimuli and cell models. For the four conditions, HRG- and EGF-stimulation of MCF-7 cells and atRA- and DMSO-stimulation of HL-60 cells, the table reports the phase windows for Preparation, Measurement (active sensing), and Feedback, including the critical transition, based on time-resolved information-thermodynamic metrics (**Figures 6-7**). Green text denotes peak CP EED-work work windows (**Figure 12**), whereas red text denotes the genome engine switch window, a dynamical marker associated with the Maxwell’s demon cycle (**Figure 13**).

| Stimulus & Cell Type | Fate | MD Cycle Phase Window | Genome Engine Switch Window | CP Mechanical Work Window |
| --- | --- | --- | --- | --- |
| HRG<br>(MCF-7 cells) | Cell-fate change<br>(Differentiation) | Preparation: 10–15 min;<br>Measurement: 15–20 min;<br>Feedback: 20–30 min<br>(Critical Transition) | 20–30 min<br>Feedback Phase | CP spike-type<br>Peak at 20–30 min<br>Feedback Phase |
| atRA<br>(HL-60 cells) | Cell-fate change<br>(Differentiation) | Preparation: 2–4 h;<br>Measurement: 4–8 h;<br>Feedback: 12–18 h<br>(Critical Transition) | Switch 1: 4–8 h<br>Measurement Phase;<br>Switch 2: 18–24 h<br>Post-feedback Phase | CP spike-type<br>Peak at 48–72 h<br>Post-feedback Phase |
| DMSO<br>(HL-60 cells) | Cell-fate change<br>(Differentiation) | Preparation: 12–18 h;<br>Measurement: 18–24 h;<br>Feedback: 48–72 h<br>(Critical Transition) | 18–24 h<br>Measurement Phase | CP spike-type<br>Peak at 18–24 h<br>Measurement Phase |
| EGF<br>(MCF-7 cells) | Non-cell-fate change<br>(Proliferation) | Pre-preparation: 0–4 h;<br>Preparation: 4–6 h;<br>Measurement: 6–8 h;<br>Feedback: 12–24 h | 4–6 h<br>Preparation Phase | CP basin-type<br>Local peak at 6–8 h (Measurement Phase)<br>Initial rise at 0–15 min (Initial Signaling)<br>End peak at 48–72h (Proliferation Phase) |

**Figure 7.**
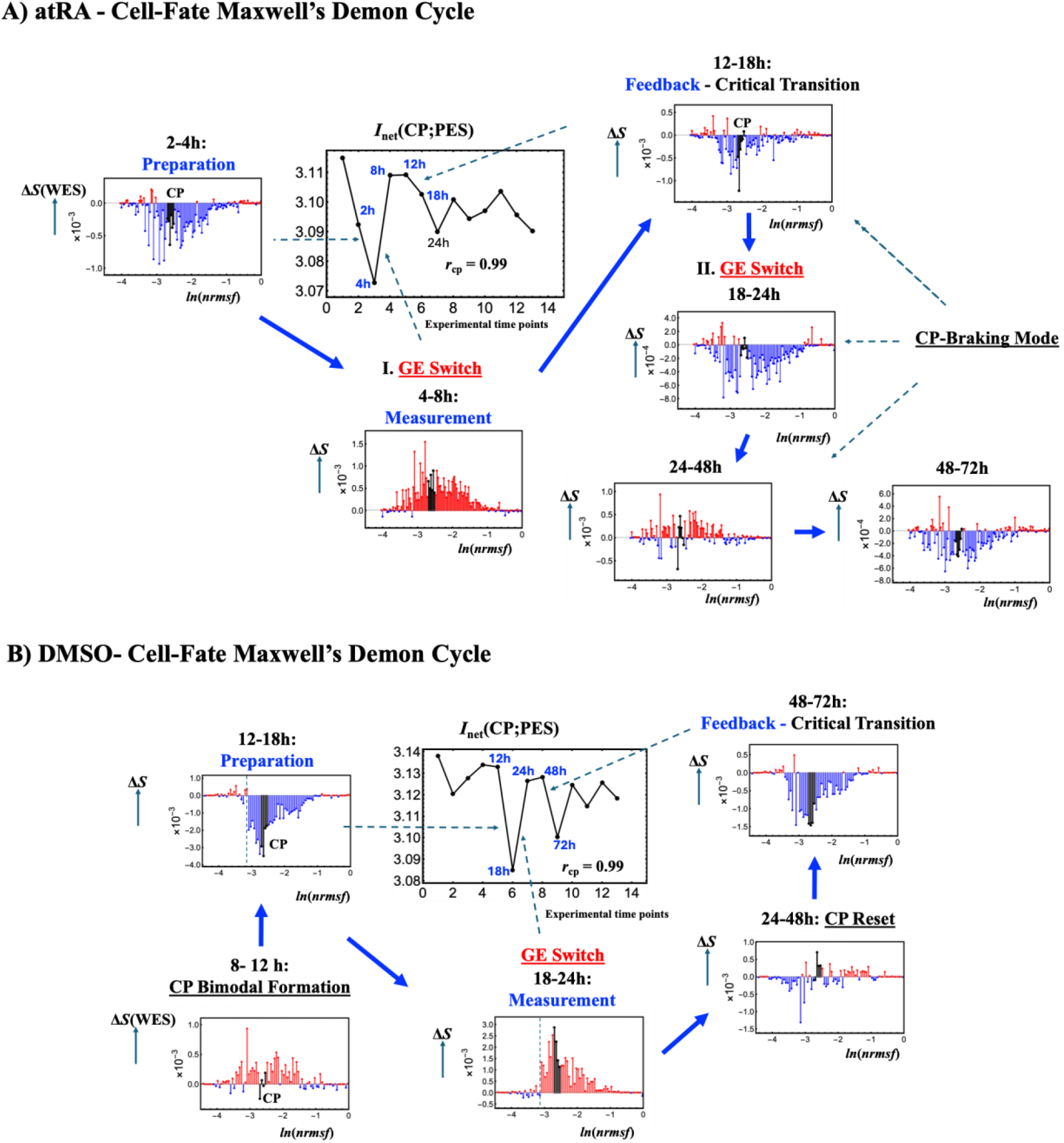
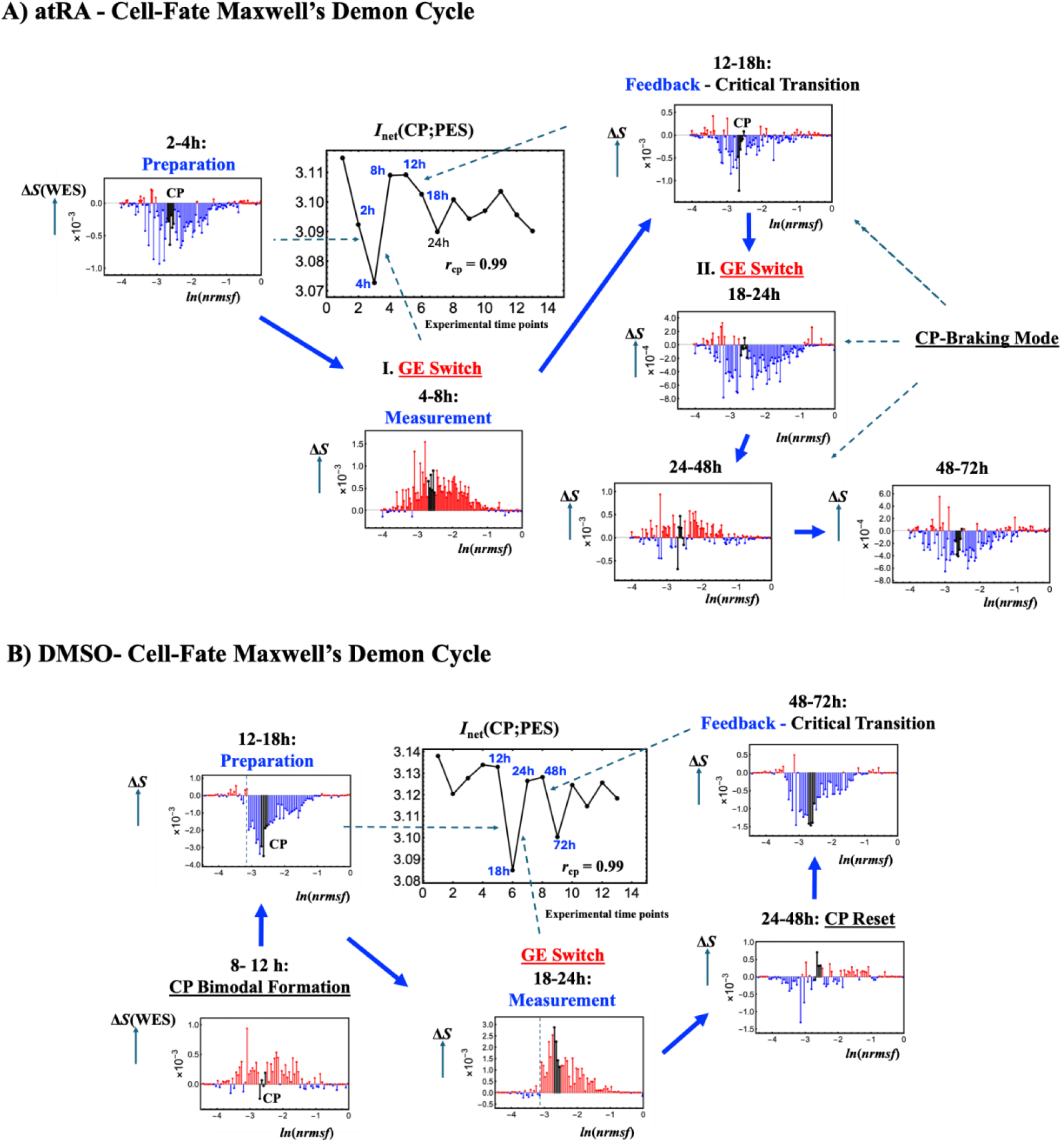
Response-speed spectrum in HL-60 fate trajectories: atRA (stepwise, two-event) versus DMSO (damped, delayed). Plotting conventions are the same as in **Figure 6** except for the CSA-derived bin resolution, with a fixed bin number of 120 for atRA to identify the CP-braking mode and 80 for DMSO. In both trajectories, *I*_net_(CP;PES) remains tightly synchronized with the CP state (*r*_cp_ = 0.99), indicating preserved baseline CP-PES coupling, while stimulus-dependent differences appear in event timing and the speed of stabilization. **(A) atRA (cell-fate change)** exhibits two GE switches (Section 2.4.5) at 4-8 h (Measurement phase) and 18-24 h (Table 3: Post-Feedback), the latter following the Feedback/critical-transition window (12-18 h). Notably, the Δ*S*(WES) profile indicates an all-or-none expression-state pattern consistent with chromatin remodeling across the *ln(nrmsf*) coordinate, and at 2-4 h, the CP region (black lines) shows only a modest negative braking signature, does not form coherent CP-driven bistable switching and therefore does not establish coherent ON/OFF expression-state switching (see the main text). In contrast, at 12-18 h (Feedback phase), the CP region starts to show clear bimodal behavior in entropy change, which later develops into a strong CP-braking mode or repressive state during 24-48 h. At 48-72 h, this bimodal pattern is largely resolved, and the CP acts coherently. The GE switch at 18 h supports the peak in CP work at 48-72 h (Table 3; Section 2.4.5). **(B) DMSO (cell-fate change)** shows 1) CP-bimodal formation before the labeled Preparation window and CP reset after the Measurement phase, reflecting an early re-initialization route before Feedback; and 2) a delayed Feedback route, with CP-driven coherent chromatin remodeling under an MD double-well potential profile and a damped, oscillatory *I*_net_ response. The GE switch at 18-24 h (Table 3: Measurement phase) is followed by genome-wide reorganization through the 48-72 h Feedback critical transition.

Across conditions, these common features of CP, the genomic Maxwell’s demon, are summarized as the MD cycle:

**(i) Preparation phase, initialization and ordering.** During Preparation, HRG, atRA, and DMSO show a deep negative Δ*S*(WES) excursion, indicating genome-wide chromatin ordering and the lowest CP entropy configuration within the analyzed time window. This defines the initial synchronized CP, PES, and WES state under CP-PES phase synchronization. The same window also shows the deepest negative Δ*I*_net_(CP;PES), marking a signature of cell-fate-changing trajectories (**Figures 6**-**7**). In contrast, EGF shows only shallow local minima in Δ*S*(WES) and Δ*I*_net_(CP;PES), with an approximately fivefold smaller Δ*I*_net_ magnitude than HRG, consistent with incomplete genome-wide reorganization (**Figure 4B**).
**(ii) Measurement phase, active sensing**. **Figures 4**-**5** show that the CP and PES are actively driven under CP-PES phase synchronization, producing concurrent positive entropy increments in the CP and PES (Δ*S*(CP) > 0 and Δ*S*(PES) > 0) together with a net increase in *I*_net_(CP;PES) (Δ*I*_net_(CP;PES) > 0). Thus, CP-driven active sensing is defined by increased CP-PES information coupling under synchronized entropy expansion. Importantly, information gain does not require PES entropy reduction. Instead, it occurs together with Δ*S*(PES) > 0.

In HRG, atRA, and DMSO, the Measurement window reaches trajectory-wide maxima of Δ*S*(CP), Δ*S*(PES), and Δ*I*_net_(CP;PES), indicating maximal entropy expansion and information gain before Feedback. In EGF, the same pattern appears only as a submaximal local maximum, indicating insufficient accumulation for effective actuation.

Biophysically, as shown later (Section 2.4.4 and **Note 2)**, in CP-enhancing trajectories, large positive (incoming) CP work in expression space supports positive entropy increments in both CP and PES, together with the associated information gain, Δ*I*_net_(CP;PES) > 0. This work drives coordinated chromatin opening, yielding Δ*S*(WES) > 0.

**Note 1: CP-driven active sensing and Measurement phase.** Tsuchiya et al. [2025] proposed that environmental energy influx sustains CP-PES phase synchronization during Measurement. Here, active sensing is characterized by synchronized entropy expansion, Δ*S*(CP) > 0 and Δ*S*(PES) > 0, together with information gain, Δ*I*_net_(CP;PES) > 0. As shown in Section 2.4.4, under CP-PES thermodynamic phase synchronization, this information gain is coupled to the incoming work received by the CP in expression space. In CP-enhancing responses, positive incoming CP work supports the concurrent entropy increases during Measurement. Thus, active sensing is mediated by information-work coupling rather than requiring environmental energy influx.

**(iii) Feedback phase: information decrease and ordering.** During Feedback, the information gained during Measurement decreases, with Δ*I*_net_(CP;PES) < 0, together with genome-wide ordering characterized by Δ*S*(WES) < 0 (**Figures 6**-**7**). Its coupling to positive incoming CP work and net work is condition dependent and determines whether Feedback proceeds to fate execution, as detailed in Section 2.4.5.

In summary, the MD cycle retains a common three-phase structure but produces condition-specific chromatin-remodeling responses (Figures 6–7). HRG and DMSO show CP-driven double-well remodeling, EGF shows a weaker CP-enhancing response, and atRA shows all-or-none remodeling with CP braking. Section 2.4.5 details the condition-specific commitment and execution windows.

**Note 2: Physical meaning of Δ*I*net(CP;PES) ≈ Δ*S*(CP) in expression space.** The CP drives rewritable chromatin memory through mode-specific state rewriting: CP-enhancing switching in HRG, DMSO, and EGF, and CP-braking dynamics in atRA (**Figures 6-7**). Across all four conditions, Δ*S*(WES) tracks Δ*S*(CP) (**Figures 4-5**). From Eq. 3, Δ*I*net ≈ Δ*S*(CP) implies Δ*S*(WES) ≈ Δ*S*(PES). Thus, changes in CP-PES information coupling are directly reflected in CP entropy changes, while PES and WES entropy changes remain approximately matched. Together with the mode-specific state rewriting in **Figures 6-7**, this relation shows that CP-driven chromatin-memory rewriting remains synchronized with WES entropy dynamics, providing an operational basis for coherent MD feedback.

**Panel I** shows the information-thermodynamic metrics using (a) rank-independent pairing and (b) expression-rank pairing. The net mutual inoformation, *I*_net_(CP;PES) is identical under rank-independent and expression-rank pairing and remains tightly synchronized with *S*(CP) in both HRG and EGF (*r*_CP_ = 0.99). This demonstrates CP-PES thermodynamic phase synchronization across all experimental time points. Expression-rank pairing decomposes *I*_net_ into the rank-matched pairwise component, *I*_rm_, and the positive higher-order information contribution, *I*_high,rm_ = *I*_net_ − *I*_rm_ > 0 (see main text). Notably, in HRG, even the higher-order information contribution exhibits strong phase synchronization with CP entropy (*r*_CP_ = 0.95).

Panel II compares Δ*S*(CP), Δ*S*(PES), Δ*S*(WES), and Δ*I*_net_, plotted on the same vertical scale within each condition. Their temporal correlations with Δ*S*(CP) are indicated by *r*_ΔCP_. HRG exhibits pronounced, coordinated increases in entropy and net mutual information during the 15-20 min Measurement phase (active sensing). EGF also exhibits increases during its 6-8 h Measurement phase. Comparing these respective Measurement windows, however, the amplitudes of Δ*I*_net_ and Δ*S*(CP) in EGF are approximately fivefold smaller than in HRG (factors of 4.7 and 4.8, respectively), while Δ*S*(WES) is approximately fourfold smaller (factor of 4.1). These differences distinguish the strong information-thermodynamic response in HRG from the weaker response in non-committing EGF.

**Panel I** shows (a) pairing-independent information-thermodynamic metrics and (b) their expression-rank-matched decomposition. As in MCF-7 cells, *I*net(CP;PES) remains tightly synchronized with *S*(CP) in both conditions (*r*_CP_ = 0.99). In atRA, the rank-matched pairwise information term, *I*_rm_(CP;PES), and the higher-order information contribution, *I*_high,rm_(CP;PES), exhibit opposite-phase dynamics during the post-MD period (24-72 h), marked by dashed lines. During 24-72 h, the rank-matched pairwise information term acts as a local braking term, whereas the increase in net CP-PES information coupling is driven by the higher-order information contribution. In contrast, DMSO exhibits damped oscillatory coupling with progressively reduced amplitude.

**Panel II** compares Δ*S*(CP), Δ*S*(PES), Δ*S*(WES), and Δ*I*net(CP;PES), with correlations indicated as in **Figure 4**. The two fate-guiding HL-60 conditions exhibit distinct MD-cycle timing: atRA undergoes Preparation at 2-4 h, Measurement at 4-8 h, and Feedback at 12-18 h, whereas DMSO undergoes the corresponding phases at 12-18 h, 18-24 h, and 48-72 h.

### 2.4 Unified Genome Engine and Maxwell’s Demon Cycle via Their Convergence

We demonstrate that the CP drives the WES both as an information-thermodynamic MD operator through rewritable chromatin memory and as the SOC organizing center of dynamic criticality (see Section 3.1 for physical interpretation and limitations). The convergence of these two functions, with condition-specific timing relative to the MD cycle, supports a biophysical interpretation of cancer-fate control under open thermodynamic constraints.

#### 2.4.1 Genome Engine Dynamical System

Since 2016, we have developed Expression Flux Analysis (EFA) [Tsuchiya et al., 2016], with major extensions in [Tsuchiya et al., 2020, 2023]. Here, we further update EFA to integrate it with expression-derived information thermodynamics in an open-system framework. EFA is based on coherent stochastic behavior (CSB), in which the law of large numbers drives convergence of ensemble-averaged gene-group trajectories despite substantial single-gene noise, even in an open, non-equilibrium genomic system. As statistically validated in Section 2.2, this CSB convergence supports treating each gene-expression contribution as a unit-mass (*m* = 1) element.

In this representation, the unit-mass CM expression of each domain attractor serves as the effective expression-derived (EED) position, its first-order difference as the EED momentum, and its negative second-order difference as the self-flux. The coarse-grained representation organizes these interacting domain fluxes into the one-dimensional genome-engine network, which captures when and how genome-wide transitions occur through criticality, specifically self-organized criticality (SOC) [Tsuchiya et al., 2016, 2020, 2023a]. At this coarse-grained level, the use of Δ*t* = 1 as a unit time forward step is motivated by the following three observations:

**1. Sandpile-type critical transition profile as a signature of state change** [Tsuchiya et al., 2016, 2020, 2023]. The sandpile-type critical transition profile observed between experimental states reveals a hallmark of SOC at the coarse-grained level, with the CP at its summit. A change in the CP propagates throughout the whole expression system (WES) as a genome avalanche. Partial or complete loss of the sandpile profile is observed in cancer and immune cell populations and, at the single-cell level, mouse and human embryos. These losses mark erasure of initial-state information and identify genome-wide state transitions, together with their associated transition-time windows (cf. Figs. 3, 5, 7 in [Tsuchiya et al., 2016]), as discussed in relation to Landauer erasure in Section 3.1. Thus, each transition can be treated as one unit interval, Δ*t* = 1, independent of the actual elapsed clock time.
**2. Embryonic development directly realizes discrete state transitions**. Development proceeds through successive biological states such as the zygote, 2-cell, and 4-cell stages. These states are defined by biological events rather than equal elapsed clock times. Each transition therefore represents one biological state change and can naturally be assigned Δ*t* = 1, independent of the actual duration of the developmental interval.
**3. Convergence between self-flux and Δ*S*(CP)**. In our study (Section 2.4.3), the CP self-flux, the EED force acting on the CP, shows temporal phase convergence with the CP entropy change, Δ*S*(CP). Entropy is a thermodynamic state function and depends only on the initial and final CP states. Here, Δ*t* = 1 represents one forward step along the observed sequence of genome states, independent of the actual elapsed clock time. Thus, this convergence supports the use of Δ*t* = 1 as the coarse-grained forward step between successive genome states.

Together, these results support setting Δ*t* = 1 in EFA. Thus, Δ*t* = 1 and *m* = 1 define the coarse-grained representation of genome-state dynamics.

Taking the genome attractor (GA) trajectory as the baseline, the genome engine (GE) is formulated as a one-dimensional dynamical network of domain attractors that incorporates the cellular environment [Tsuchiya et al., 2024]. The GE detects critical transitions and quantifies inter-attractor coupling, including cyclic flux and phase synchronization.

To illustrate the temporal structure, consider a ball striking a wall at *t_j_*: it arrives over [*t*_j-1_, *t*_j_] and departs over [*t*_j_, *t*_j+1_], so its momentum change is determined by the difference between its outgoing and incoming momenta. Likewise, each domain attractor is treated as an effective unit-mass particle in expression space, with IN flux representing its incoming expression momentum and OUT flux its outgoing expression momentum.

This analogy illustrates the temporal ordering of the two intervals; the effective expression-derived (EED) force acting on the *i*^th^ domain attractor at *t*ⱼ, *F_i_* (*t_j_*), is defined by the difference between the IN and OUT fluxes. A positive EED force denotes the incoming direction toward the domain attractor, whereas a negative EED force denotes the outgoing direction.

Accordingly, for domain attractor *i* ∈ {CP, Super, Sub}, the EED force, *F_i_*(*t_j_*), termed self-flux, acting on the *i*^th^ domain attractor at *t*_j_ is defined as

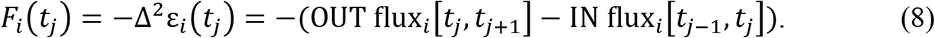

Here *ε_i_*(*t_j_*) is the expression of the *i*^th^ domain attractor, and the second order expression difference is Δ^2^ε*_i_*(*t_j_*) = *ε_i_*(*t_j_*_+1_) − 2*ε_i_*(*t_j_*) + *ε_i_*(*t_j_*_−1_). IN flux*_i_*[*t_j_*_−1_, *t_j_*] = *ε_i_*(*t_j_*) − *ε_i_*(*t_j_*_−1_) represents the incoming flux over the preceding interval, whereas OUT flux*_i_*[*t_j_*, *t_j_*_+1_] = *ε_i_*(*t_j_*_+1_) − *ε_i_*(*t_j_*) represents the outgoing flux over the forward interval. Thus, *F_i_*(*t_j_*) > 0 is interpreted as incoming EED force, with IN flux exceeding OUT flux, whereas *F_i_*(*t_j_*) < 0 indicates the opposite direction.

For the GE network, the origin of the IN flux and the destination of the OUT flux do not affect the definition of EED force at the *i*^th^ domain attractor. Thus, the ball-striking-a-wall analogy extends directly to the interacting EED force, defined by the difference between the IN flux and the OUT flux using the same sign convention as the self-flux.

This interacting EED force is denoted by the bra-ket notation ⟨*k*|*i*⟩(*t_j_*) (see **Note 1** below), with its positive direction defined as incoming from *k* → *i*. With this notation, the self-flux of the *i*th domain, *F_i_*(*t_j_*), is written as ⟨*i*|*i*⟩(*t_j_*). Note that this directional convention is opposite to the conventional bra-ket reading: *i* → *k*.

Accordingly, the directed interaction-flux term between domains *k* and *i* is defined as

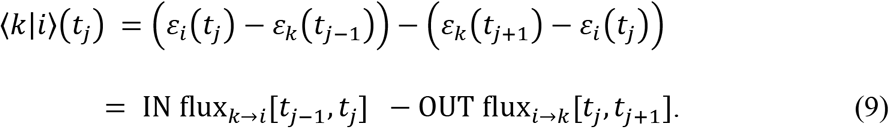

Here IN flux*_k_*_→*i*_[*t_j_*_−1_, *t_j_*] is the incoming flux from domain *k* to domain *i* over [*t*_j-1_, *t*_j_], and OUT flux*_i_*_→*k*_[*t_j_*, *t_j_*_+1_] is the outgoing flux from domain *i* to domain *k* over [*t*_j_, *t*_j+1_]. Thus, ⟨*k*|*i*⟩(*t_j_*) > 0 denotes that the incoming flux from *k* to *i* exceeds the outgoing flux from *i* to *k*, whereas ⟨*k*|*i*⟩(*t_j_*) < 0 denotes the reverse inequality.

Hereafter, we use “domain” or “attractor” in place of “domain attractor” unless the distinction needs to be made explicit.

**Note 1. Temporal separation of IN and OUT fluxes and meaning of bra-ket notation.** EED force is defined from the IN and OUT fluxes over [*t*_j-1_, *t*_j_] and [*t*_j_, *t*_j+1_], respectively. In a later section, this temporal separation provides the basis for defining EED work and analyzing the irreversible time arrow of cancer-fate change. The ket | *i*⟩ is used only as a domain label; self-flux and interaction-flux bra-kets denote flux quantities, not vector-space inner products.

Consider a multidomain GE network consisting of the CP, Super, and Sub domains. Each domain self-flux is decomposed into interaction fluxes with the other domains and a common external flux. In the three-domain case, labeling CP, Super, and Sub as 1, 2, and 3 gives the following decomposition of the CP self-flux, ⟨1|1⟩(*t_j_*), using an add-and-subtract construction:

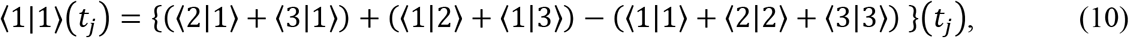

where ⟨1|2⟩, ⟨1|3⟩, ⟨2|1⟩, and ⟨3|1⟩ represent directed pairwise interaction fluxes coupling domain 1 with domains 2 and 3. In each bra-ket, the right-hand index identifies the reference domain. Thus, ⟨1|2⟩ is referenced to domain 2, whereas ⟨2|1⟩ is referenced to domain 1. For instance, ⟨1|2⟩(*t_j_*) = IN_1→2_[*t_j_*_−1_, *t_j_*] − OUT_2→1_[*t_j_*, *t_j_*_+1_], using the same sign convention as the self-flux in Eq. 8. The other interaction terms are defined analogously. The same add-and-subtract construction extends to a network of *N* domain attractors, as derived in **Note 2**.

The sum of the three domain self-fluxes appearing in Eq. 10 is

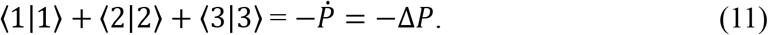

Here, the EED force, *F_i_*(*t_j_*) = ⟨*i*|*i*⟩(*t_j_*) = −Δ^2^ε*_i_*(*t_j_*) = −*ṗ_i_* for *i* ∈ {1, 2, 3}, where *p_i_*is the unit-mass EED momentum of domain *i*. Because *F_i_*is defined as positive in the incoming direction, opposite to the forward EED-momentum change *ṗ_i_*, *F_i_* = −*ṗ_i_*. The dot denotes the discrete rate of momentum change, and *ṗ_i_* = Δ*p_i_* with Δt =1. Thus, 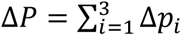, is the total EED momentum change of the three domains.

Eq. 10 shows that the external flux is common to all domain attractors. Accordingly, the common external flux is defined as

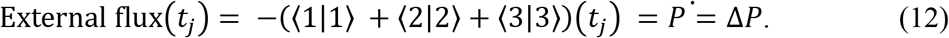

The external flux closes the expression-flux balance of the open genome system. Positive external flux is interpreted as incoming flux from the environment into the WES, whereas negative external flux is interpreted as outgoing flux from the WES to the environment.

### Note 2. Reciprocal interaction fluxes and self-flux decomposition

Eq. 9 identifies the reciprocal interaction-flux pair ⟨CP|Super⟩ and ⟨Super|CP⟩ (**Figure 8**). Their anti-phase relation is interpreted as a signature of cyclic flux between CP and Super. From Eqs. 8 and 9, ⟨*i*|*k*⟩ + ⟨*k*|*i*⟩ = −Δ^2^*ε_i_* − Δ^2^*ε_k_* = ⟨*i*|*i*⟩ + ⟨*k*|*k*⟩. All quantities are evaluated at *t*_j_. Thus, reciprocal interaction fluxes cancel exactly when the two domain self-fluxes sum to zero. The magnitude of this sum measures incomplete cancellation, not the amplitude of cyclic exchange or coupling strength. These terms describe expression-flux balances and need not obey Newtonian action-reaction symmetry. For *N* domains, summing this identity over *k* ≠ *i* gives

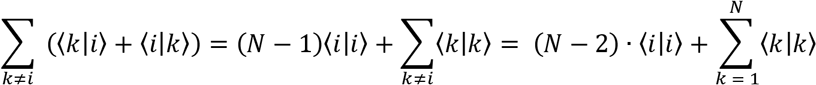

For *N* = 3, subtracting the total self-flux recovers Eq. 10.

**Figure 8.**
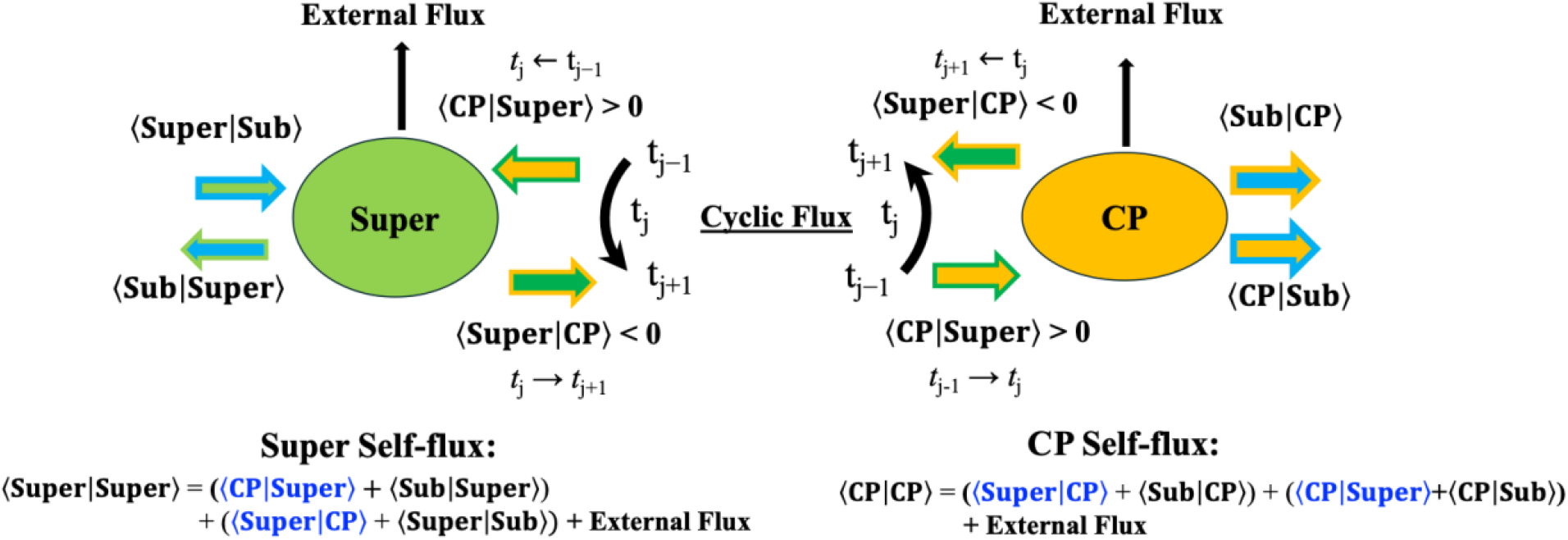
Schematic illustration of cyclic flux formation through decomposition of the CP and Super self-fluxes. This schematic shows how the interaction fluxes ⟨CP|Super⟩ and ⟨Super|CP⟩ exhibit an anti-phase relation within the CP and Super self-flux decompositions. Near cancellation requires nearly equal magnitudes and opposite signs, ⟨CP|Super⟩ + ⟨Super|CP⟩ ≈ 0. At *t*_j_, a positive ⟨CP|Super⟩ indicates that incoming flux from CP to Super over [*t*_j-1_, *t*_j_] exceeds outgoing flux from Super to CP over [*t*_j_, *t*_j+1_]. A negative ⟨Super|CP⟩ indicates that outgoing flux from CP to Super over [*t*_j_, *t*_j+1_] exceeds incoming flux from Super to CP over [*t*_j-1_, *t*_j_]. These signs describe complementary balances associated with the CP-to-Super direction under Eq. 9. A closed cyclic sequence additionally requires the reciprocal temporal pattern. The degree of cancellation characterizes the balance between the reciprocal interaction fluxes; their amplitudes and temporal organization are also needed to assess cyclic coupling.

#### 2.4.2 Dynamic CP-GA Phase Synchronization During Maxwell’s Demon Cycle

We examine how domain self-fluxes and CP-GA phase synchronization enable the genome engine to maintain cell-ensemble homeostasis while driving a critical transition, as summarized below:

**(i) Homeostatic stability. Figure 9** shows that the temporal average of each domain self-flux is *O*(10⁻⁴), compared with directed interaction-flux signals of *O*(10⁻¹), so that, for fixed *i*,

**Figure 9.**
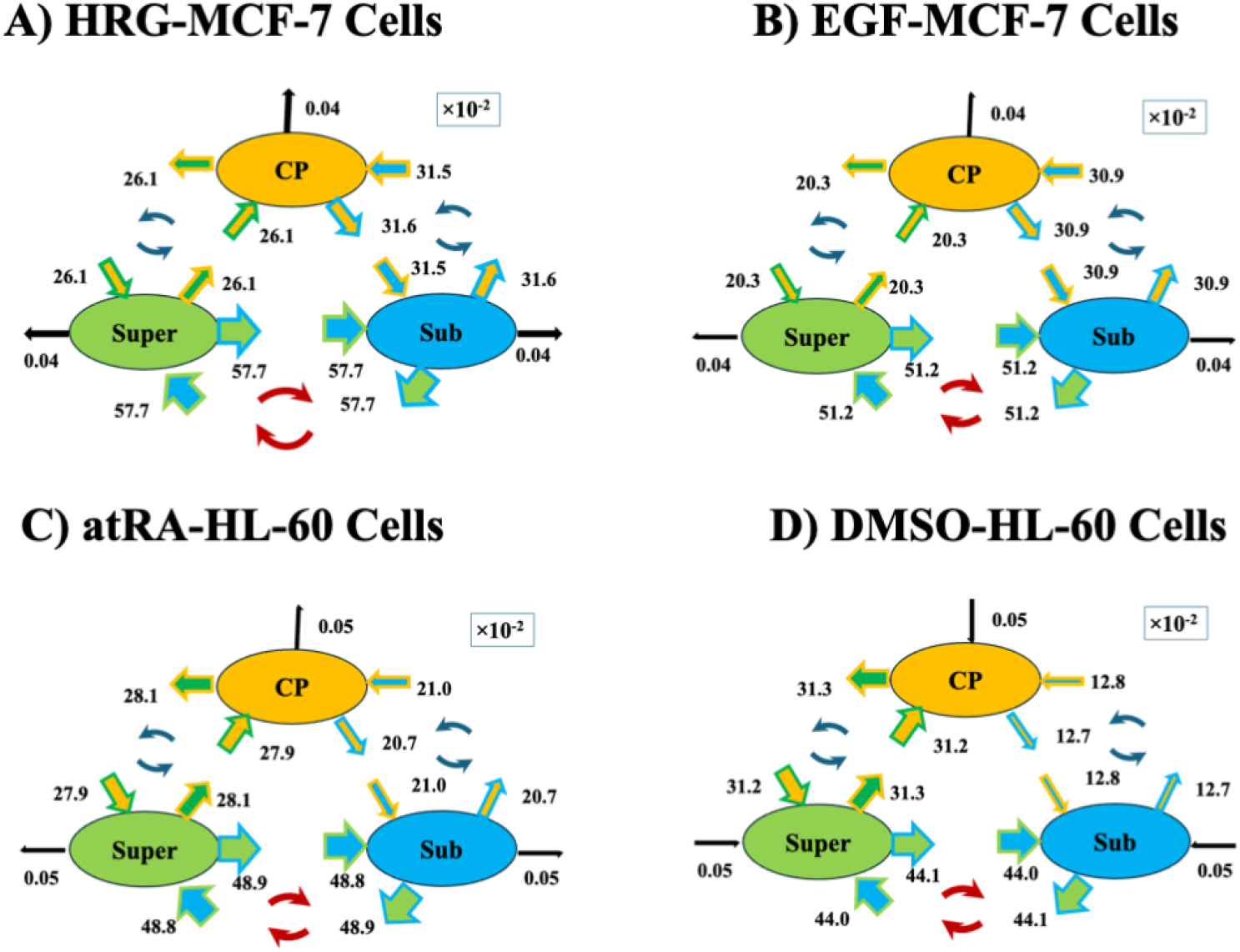
Temporally averaged expression-flux network for MCF-7 and HL-60 cancer cells. The network summarizes the temporally averaged interaction fluxes among the three domains and the external environment in the open cell-culture system. An arrow represents an interaction flux. For example, the interaction flux <CP|Sub> from the CP (orange) into the Sub domain (blue) is represented by the arrow with the same CP orange fill and the blue contour of the Sub domain. Whether the flux is incoming or outgoing is determined by the sign of the corresponding interaction flux, as described in the main text. Across all four stimuli, the temporally averaged domain self-fluxes are nearly balanced, the temporal average of the cell-culture environmental flux is negligible (*O*(10^−4^)), and a temporally averaged cyclic flux network forms among the CP-Super, CP-Sub, and Super-Sub domains. Numerical results (×10^−2^) are rounded to three significant figures.

**Figure 10.**
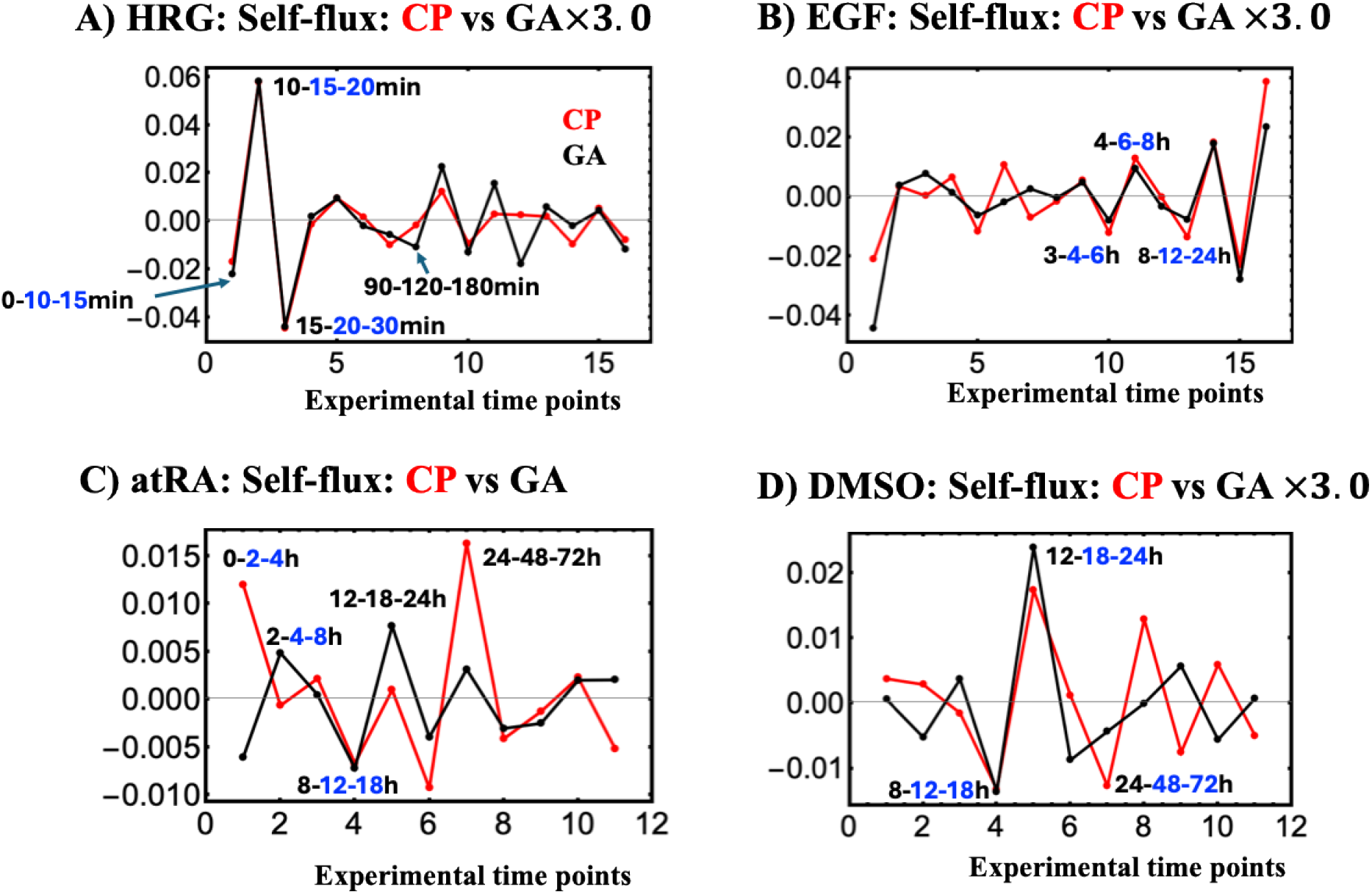
Synchronization of the CP with the Genome Attractor (GA). **A)**: HRG- and **B)** EGF-stimulated MCF-7 cells. **C)**: atRA- and **D)** DMSO-stimulated HL-60 cells. Black and red solid lines denote the GA and CP self-fluxes (effective forces acting on the GA and CP), respectively. Black and blue text indicate the self-flux three-time-point window [*t*_j-1_,*t*_j_,*t*_j+1_] at the indicated time point. Blue text identifies the Maxwell’s demon (MD) time windows: Preparation (first blue window), Measurement (second), and Feedback (third).

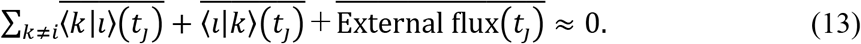

This indicates homeostatic EED-force balance for each domain through compensation among internal directed interaction fluxes that are large relative to the temporally averaged external flux, supporting the interpretation that EED force functions internally as a force-like dynamical quantity in coarse-grained expression space. However, the external flux becomes large at the critical transition [Tsuchiya et al., 2020], accompanying cyclic EED-force exchange among the CP, Super, and Sub domains.

**(ii) Critical transition**. **Figure 10** shows that the CP self-flux synchronizes with the genome attractor (GA) during the MD cycle, except during the atRA MD cycle. This CP-GA phase synchronization is the dynamical counterpart of the thermodynamic synchronization in Section 2.3, linking active sensing during Measurement to CP-driven EED actuation in expression space during Feedback, which induces the critical transition. The resulting CP changes propagate through the GA and WES as a genome-wide expression avalanche [Tsuchiya et al., 2020, 2023], the dynamical signature of criticality.
**(iii) atRA exception**. During the Preparation-to-Measurement window, CP and GA are anti-phase, undergoing opposite self-flux sign changes and thereby exhibiting directed CP-GA asymmetry before phase synchronization emerges after the MD cycle (**Figure 10C**). The corresponding information-thermodynamic mechanism is addressed in the following subsections.
**(iv) Unified role of CP**. The CP controls the genome through two coupled functions: dynamically, as the GE control center through EED force-driven actuation in expression space, and information-thermodynamically, as the MD through rewritable chromatin memory. These functions converge during the MD cycle, coupling CP-driven information gain to EED actuation in a single genome-wide regulatory event (see the next section).

**For HRG, EGF, and DMSO**, the figures show synchronization of the CP self-flux (red) with the genome attractor (GA) self-flux (black, scaled×3.0) during the Maxwell’s demon (MD) cycle. This CP-GA phase synchronization supports the interpretation that changes in the CP propagate through the GA as a genome-wide expression avalanche [Tsuchiya et al., 2023a]. This timing match between CP-GA synchronization and the Maxwell’s demon cycle supports EED actuation in expression space, with condition-specific transition and execution timing. This timing is consistent with CP-dominated mechanical work actuation during the MD cycle (Section 2.4.4). HRG and DMSO trigger cell-fate guiding transitions, whereas EGF triggers a non-cell-fate guiding transition.

**For atRA**, the behavior differs. During the Preparation and Measurement phases, anti-phase CP-GA coupling is observed, consistent with the CP-braking mode. CP and GA already share a negative trough at 12 h during Feedback, with further aligned sign changes after the MD cycle. The pronounced CP self-flux at 48 h (evaluated over the 24-48-72 h window) is consistent with CP-dominated EED work actuation at 48-72 h emerging after the MD cycle (Section 2.4.4).

#### 2.4.3 Forward Time Alignment: Convergence of Self-Flux with CP Entropy Change

In previous sections, Shannon entropy is calculated from the probability distribution of stochastic gene-expression variation at each time point along the genome attractor trajectory. Its temporal trajectory is statistically validated by the ITA bootstrap-convergence analysis in Section 2.2. In stochastic thermodynamics, Shannon entropy is used to describe uncertainty about a system’s physical state within a model that specifies its dynamics and exchanges with the environment [Seifert, 2012; Van den Broeck and Esposito, 2015; Parrondo et al., 2015].

Note that the entropy calculated here is not a direct measurement of thermodynamic entropy because a quantitative mapping between the gene-expression probability distributions and thermodynamic state variables has not been established, as discussed in **Section 3.1**.

We treat the WES within a dimensionless EED dynamical framework, with *m* = 1 and unit time Δ*t* = 1. Self-flux, −Δ^2^*ε_i_*, and Δ*S*(CP) are dimensionless. Their convergence is evaluated through forward temporal alignment and signed phase scaling.

**Figure 11** shows condition-specific convergence of domain-attractor self-flux, ⟨*i*|*i*⟩(*t_j_*), with ΔS(CP) during the MD cycle and, for atRA, after the cycle. Under CP-PES synchronization, Δ*S*(CP) scales with Δ*I*_net_(CP;PES) (**Figures 4**-**5**). These relations are summarized by

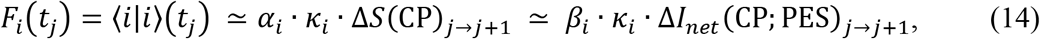

which captures the leading signed phase relation. Here, *α_i_*and *β_i_* are positive scaling constants for domain *i*; *κ_i_* = +1 and *κ_i_* = −1denote the in-phase and anti-phase relations, respectively, over [*t*_j_, *t*_j+1_], between the self-flux of domain *i* and Δ*S*(CP). Because Δ*S*(CP) and Δ*I*_net_(CP;PES) are in phase under CP-PES synchronization, the same *κ_i_* applies to the second scaling relation.

**Note 1: Average effective expression-derived (EED) force.** The domain-attractor self-flux ⟨*i*|*i*⟩(*t_j_*) is the negative difference between the OUT flux over [*t*_j_, *t*_j+1_] and the IN flux over [*t*_j-1_, *t*_j_] (Eq. 8). With unit mass *m* = 1 and time step Δ*t* = 1, these fluxes represent the average momenta over the two adjacent unit intervals, and their difference gives the average EED force, ⟨*i*|*i*⟩(*t_j_*) at *t*_j_.

**Note 2: CP-GA dynamical phase synchronization and self-flux-CP entropy convergence**. Here, “dynamical” distinguishes CP-GA phase synchronization from CP-PES thermodynamic phase synchronization. For HRG, EGF, and DMSO, Eq. 14 holds during the MD cycle: the CP and Sub self-fluxes converge with Δ*S*(CP) under CP-GA dynamical phase synchronization. In atRA, CP-GA anti-phase behavior during Preparation and Measurement is consistent with the CP-braking mode, whereas CP-GA dynamical phase synchronization emerges after the MD cycle, when clear in-phase convergence with ΔS(CP) appears only for the Super self-flux (**Figure 11C**).

This convergence supports the forward time alignment between domain self-flux and Δ*S*(CP): the domain self-flux at *t*_j_, computed from the three-point sequence [*t*_j-1_, *t*_j_, *t*_j+1_], shows convergence with the entropy change Δ*S_j_*_→*j*+1_over the interval [*t*_j_, *t*_j+1_]. This sets the forward time alignment over [*t*_j_, *t*_j+1_]. For example, the CP self-flux at 15 min, derived from the 10-15-20 min sequence corresponds to Δ*S*(CP) over the 15-20 min interval, denoted by Δ*S*(CP)_15→20 min_.

With this alignment, a positive effective force, ⟨*i*|*i*⟩(*t_j_*) > 0, corresponding to net incoming flux, is associated with Δ*S*(CP) > 0 under in-phase alignment and Δ*S*(CP) < 0 under anti-phase alignment. A negative effective force, ⟨*i*|*i*⟩(*t_j_*) < 0, corresponding to net outgoing flux, reverses these relations.

Applying this alignment rule reveals the condition-dependent convergence architectures between domain-attractor self-fluxes and CP entropy change, as summarized in **Figure 11**:

**For the HRG and EGF conditions**, within the MD-cycle window, the leading in-phase relations are ⟨CP|CP⟩(*t_j_*) ≃ Δ*S*(CP)*_j_*_→*j*+1_ and ⟨Sub|Sub⟩(*t_j_*) ≃ Δ*S*(CP)*_j_*_→*j*+1_, whereas the leading anti-phase relation is ⟨Super|Super⟩(*t_j_*) ≃ −Δ*S*(CP)*_j_*_→*j*+1_.

**For the DMSO condition**, the same phase architecture as the HRG and EGF conditions is observed, with Δ*S*(CP)*_j_*_→*j*+1_ scaled by approximately 0.4 for comparison with the CP and Sub self-fluxes, whereas the Super self-flux shows a clear anti-phase relationship with unscaled Δ*S*(CP)*_j_*_→*j*+1_ during the MD cycle.

**For the atRA condition**, within the MD-cycle window, the domain-attractor self-fluxes and Δ*S_j_*_→*j*+1_(CP) exhibit mixed-phase behavior associated with the CP-braking mode (**Figure 7A**). After the MD-cycle window, the leading in-phase relation is ⟨Super|Super⟩(*t_j_*) ≃ Δ*S*(CP)*_j_*_→*j*+1_, whereas ⟨CP|CP⟩(*t_j_*) remains mixed-phase, shifting from anti-phase to in-phase behavior, while ⟨Sub|Sub⟩(*t_j_*) exhibits weak anti-phase behavior.

Overall, under CP-GA phase synchronization, HRG, DMSO, and EGF share a CP-enhancing convergence architecture during the MD-cycle window, whereas atRA exhibits mixed-phase behavior during the MD cycle, followed by leading Super–CP entropy convergence after the cycle.

**Figure 11.**
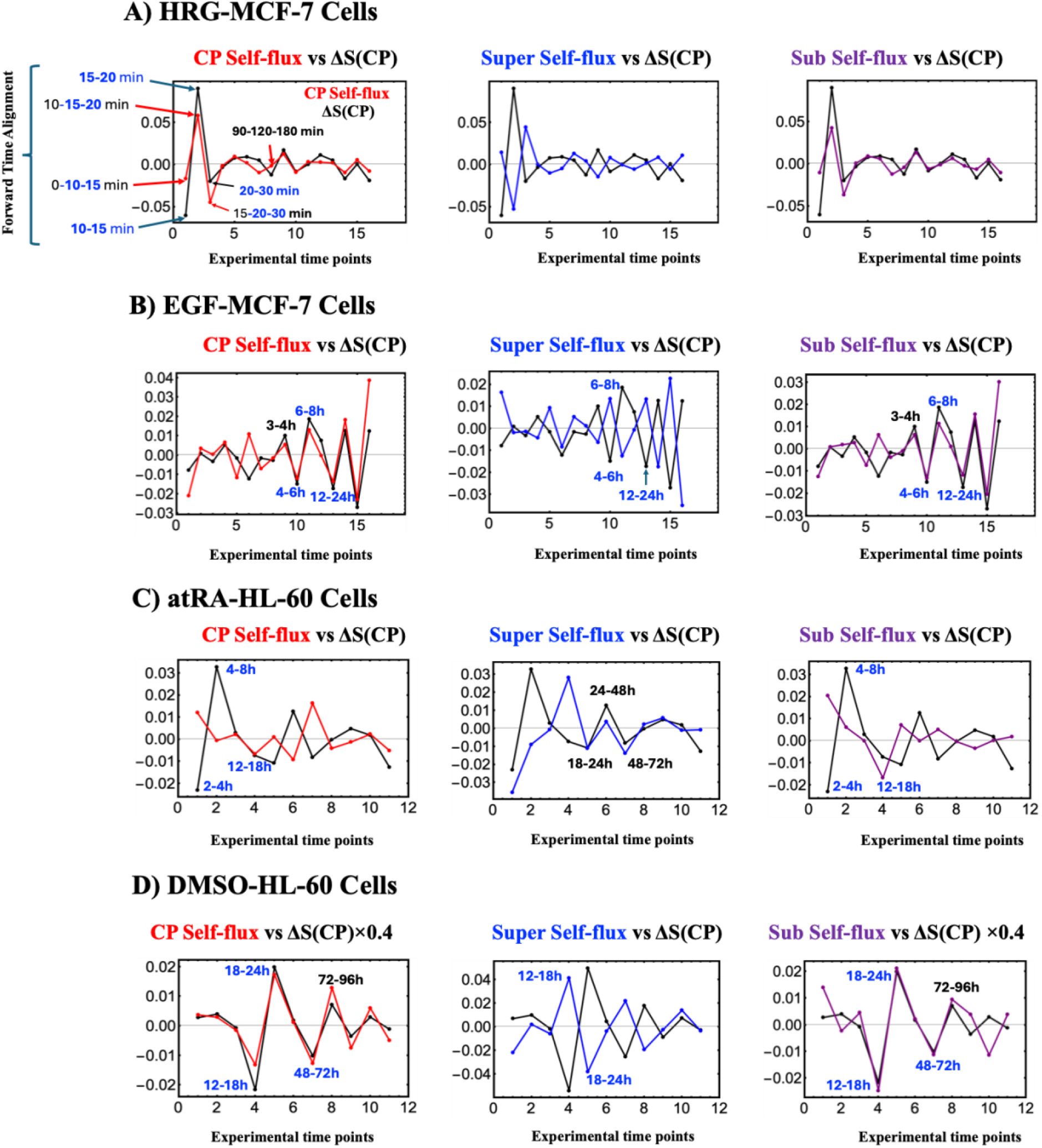
Convergence of self-flux and entropy change Δ*S*(CP) during and after Maxwell’s demon cycles. This figure illustrates the temporal phase convergence between the dimensionless self-flux signals for CP (red), Super (blue), and Sub (purple) and the CP Shannon entropy change Δ*S*(CP) (black line). The time intervals marked in blue text indicate the Preparation, Measurement, and Feedback intervals of each MD cycle. The *x*-axis uses Δ*t* = 1 between successive experimental time points. Self-flux at *t*_j_ is evaluated from the three-point sequence [*t*_j-1_, *t*_j_, *t*_j+1_], whereas Δ*S*(CP) is assigned by forward alignment to the interval [*t*_j_, *t*_j+1_]. The panels show **(A)** HRG- and **(B)** EGF-stimulated MCF-7 cells and **(C)** atRA- and **(D)** DMSO-stimulated HL-60 cells. **(A) HRG**: The CP self-flux, Sub self-flux, and Δ*S*(CP) show a sharp in-phase convergence during the MD cycle (10-30 min) under CP-GA phase synchronization (**Figure 10**); thereafter, they remain phase-aligned. In contrast, the Super self-flux exhibits a sharp anti-phase convergence during the MD cycle and remains anti-phase-aligned thereafter. Thus, HRG-stimulated MCF-7 cells exhibit a highly organized genomic SOC-control architecture. **(B) EGF:** Before the MD-cycle onset (< 4 h), the CP self-flux, Sub self-flux, and Δ*S*(CP) exhibit a mixed-phase relationship, alternating between in-phase and anti-phase. From MD-cycle onset (> 4 h), they show in-phase convergence under CP-GA phase synchronization (**Figure 10B**). The Super self-flux displays the same mixed-phase behavior before 4 h and shows clear anti-phase convergence after 4 h. (C) **atRA**: ΔS(CP) shows a clear in-phase convergence only with the Super self-flux after the MD cycle, under CP-GA phase synchronization. In contrast, Δ*S*(CP) exhibits a mixed-phase relationship with the CP self-flux, particularly after 18 h, where the relationship transitions from anti-phase to in-phase. The Sub self-flux shows only a weak anti-phase relationship with ΔS(CP) after 18 h. (D) **DMSO**: The CP and Sub self-fluxes show a clear in-phase convergence with the scaled Δ*S*(CP) (× 0.4) during the MD cycle (12–72 h), under CP-GA phase synchronization, whereas the Super self-flux exhibits a clear anti-phase convergence over the same interval. After 72 h, the Super and Sub self-fluxes continue to show distinct phase-dependent behavior.

#### 2.4.4 Unifying Effective Expression-Derived Work with CP Entropy Change

Here we employ the forward-aligned step [*t*_j_ → *t*_j+1_] to define effective expression-derived (EED) work for each domain attractor, aligning with the direction of EED force, using a sign convention in which positive EED force denotes the incoming direction toward the domain attractor.

For each domain attractor *i* ∈ {CP, Super, Sub}, the coarse-grained EED work *δW_i_* is evaluated as the negative product of two components: (i) the average incoming EED force ⟨*i*|*i*⟩(*t_j_*) acting on the *i*^th^ domain attractor at *t*_j_ and (ii) the forward interval displacement over [*t*_j_, *t*_j+1_], quantified using the OUT flux[*t*_j_, *t*_j+1_] rather than the IN flux[*t*_j-1_, *t*_j_]. Using the self-flux–entropy convergence relation in Eq. 14,

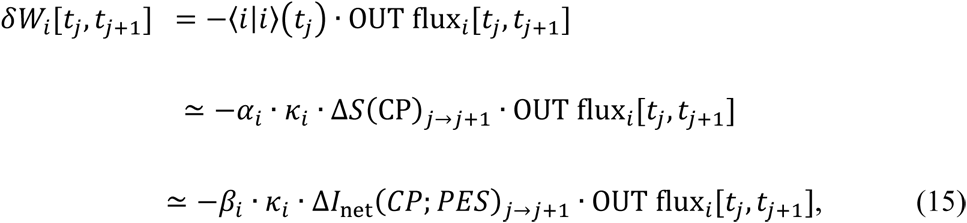

where *δ* is used because work is path-dependent, whereas Δ is used for state-function changes such as entropy change, which depend only on the endpoints. The minus sign is required because self-flux represents the average EED force on the domain attractor in the incoming-positive direction (Eq. 8), whereas the forward displacement is expressed in the outward-positive direction, analogous to a ball striking a wall as illustrated in Section 2.4.1. Thus, EED work is the negative product of incoming EED force and forward OUT displacement.

Under this sign convention, δ*W_i_* > 0 denotes incoming work to domain attractor *i*, whereas δ*W_i_* < 0 denotes outgoing work from it.

The first line gives EED work. The second and third lines give its leading signed phase scaling representations in the condition-specific windows where the convergence relation in Eq. 14 holds. Under CP-PES thermodynamic phase synchronization, Δ*S*(CP) scales with Δ*I*_net_(CP;PES). The constants *α_i_* and *β_i_* are positive scaling constants, while *κ_i_* = +1 and *κ_i_* = −1 denote in-phase and anti-phase alignment, respectively, between domain self-flux and CP entropy change. The sign of EED work also depends on the forward OUT flux.

#### Old Note 1 is removed

**Note 1: Information-to-work coupling through CP-incoming EED work**. Where the scaling relation in Eq. 14 holds, Eq. 15 operationally links net MI change, Δ*I*_net_, to CP-incoming EED work, *δW_CP_*, through the CP OUT flux. Thus, Δ*I*_net_ serves as an operational information variable for genome-engine information-work coupling. The sign of the work depends on both the signed scaling relation and the forward CP OUT flux, while, for a given CP OUT flux, the magnitude of CP EED work scales approximately linearly with |Δ*I*_net_|, which is an important marker of non-commitment in the EGF condition.

By treating the environment as an external domain, Eq. 12 defines the external flux as the environment-coupled EED force acting on the WES. Because the GA represents the CM of the WES with the unit mass, *m* = 1, its forward displacement is used to define the external EED work:

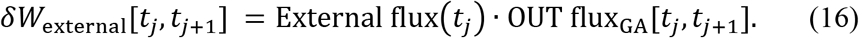

Here, OUT flux_GA_[*t_j_*, *t_j_*_+1_] = *ε*_GA_(*t_j_*_+1_) − *ε*_GA_(*t_j_*) is the forward GA displacement. This pairing defines external EED work at the coarse-grained level. *δW*_external_ > 0 indicates incoming work from the environment to the WES, whereas *δW*_external_ < 0 indicates outgoing work from the WES to the environment.

The net incoming EED work is defined as the sum of the CP, PES, and external work contributions:

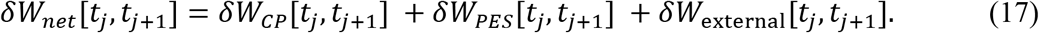

Hereafter, the time window is omitted unless explicitly required.

**Figure 12** shows that for all four conditions, the magnitude of CP EED work is more than an order of magnitude greater than that of PES EED work: δ*W*_CP_ ≫ δ*W*_PE*S*_. Strong positive Pearson correlations support CP-PES phase synchronization in EED work dynamics. Moreover, *δW*_net_ closely tracks *δW*_CP_, consistent with the CP contribution being the leading term:

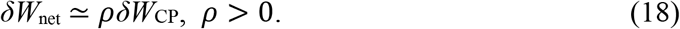

Here, *ρ* is a condition-specific positive scaling constant. This leading approximation applies to CP-dominated intervals. Maxwell’s demon represents the CP’s operational information-thermodynamic role, whereas dominant CP EED work represents its dynamical role (Section 2.4.5).

Therefore, the net coarse-grained EED work *δW*_net_ quantifies the net forward-aligned actuation in expression space for the open genome system, including both the internal domain work and the external EED work contribution:

- *δW*_net_ > 0 (net-driving mode): Net positive EED work arises from coupling among the CP, PES, and environment. This drives genome-wide expression change, supports CP-driven actuation, and suggests work-coupled outward entropy export under the thermodynamic assumptions discussed in Section 3.1 (v).
- *δW*_net_ < 0 (net-braking mode): Negative net EED work opposes forward-aligned genome actuation, indicating net braking associated with CP-braking dynamics (see below).
- *δW*_net_ ≈ 0 (execution-suppressing mode): The combined CP, PES, and external contributions yield little net EED work. Information acquisition can occur without substantial net EED actuation of the open genome system.

Furthermore, Figures 10 and 12 identify two CP actuation modes. HRG and DMSO show CP-enhancing actuation, whereas atRA shows CP-braking actuation during the early MD-cycle phases, followed by delayed positive EED actuation. EGF shows weak CP-enhancing actuation during the MD cycle, consistent with its non-committing response. Thus, the timing and magnitude of CP EED work distinguish the observed fate-changing and non-committing responses:

**(i) CP-enhancing mode**. In HRG and DMSO, CP-enhancing actuation produces positive CP work peaks and therefore positive net work peaks, defining the net driving mode during the MD cycle. Together with the CP-GA synchronization shown in **Figure 10**, this CP-dominated net actuation supports CP-GA phase synchronized dynamics during positive EED work. In EGF, during the MD cycle, the local CP work peak at 4-6 h is approximately 20-fold smaller than the HRG peak at 15-20 min. This weaker response is associated with non-commitment; a quantitative CP-work threshold for commitment has not been established.
**(ii) CP-braking mode**. In atRA, the CP-braking mode suppresses CP EED work, *δW*_CP_ and keeps Net work, *δW*_net_ ≈ 0 during the Measurement phase. Net work is slightly negative during this interval, mainly because of the negative external contribution, *δW*_external_. Although Δ*S*(CP) and Δ*I*_net_(CP;PES) reach positive peaks in the Measurement phase (**Figure 5A**), this information gain is not yet accompanied by substantial positive CP EED work. Strong positive CP and net-work peaks emerge later, at 24-48 h after the MD cycle, identifying delayed EED actuation.

Although positive net work, *δW*_net_ > 0, is observed over most time intervals in both MCF-7 and HL-60 cells, negative net work *δW*_net_ < 0 occurs during several intervals in the atRA condition within the CP-braking mode (**Figure 12C**).

In summary, EED work is the dimensionless, coarse-grained work performed by the EED force along the forward displacement of domain attractors in the open WES, linking CP-PES information-work dynamics to Maxwell’s demon actuation. Here, effective genome actuation denotes displacement and reorganization in expression space, not directly measured physical displacement of chromatin or DNA, as discussed in Section 3.1. Within this framework, the CP-enhancing and CP-braking modes define the key SOC control modes for the cell-fate-guiding critical transition. In the subsequent subsection, we address these modes through three timelines for fate commitment and the resulting time arrow of cancer fate control.

**Note 3: Stochastic resonance-like mechanism.** For cell-fate-guiding conditions (HRG, atRA, and DMSO), the pronounced positive peaks in CP work (**Figure 12**) arise from the **negative product** of the CP self-flux and the forward OUT flux in Eq. 15. Both quantities describe coarse-grained stochastic expression dynamics relative to the genome attractor (GA) trajectory used as the baseline. Positive CP work occurs when the self-flux and OUT flux have opposite signs. The emergence of pronounced CP-work peaks from these stochastic expression dynamics, together with CP-associated bimodal state transitions, suggests a stochastic resonance-like mechanism [Gammaitoni et al., 1998; McDonnell and Abbott, 2009]. In this interpretation, stochastic expression fluctuations interact with the threshold-like CP structure to enhance EED work within SOC-controlled genome dynamics. This motivates the hypothesis of “self-organized critical stochastic resonance.” Whether stochastic fluctuations facilitate CP bimodal-state reconfiguration and enhance actuation coherence remains to be tested.

**Figure 12.**
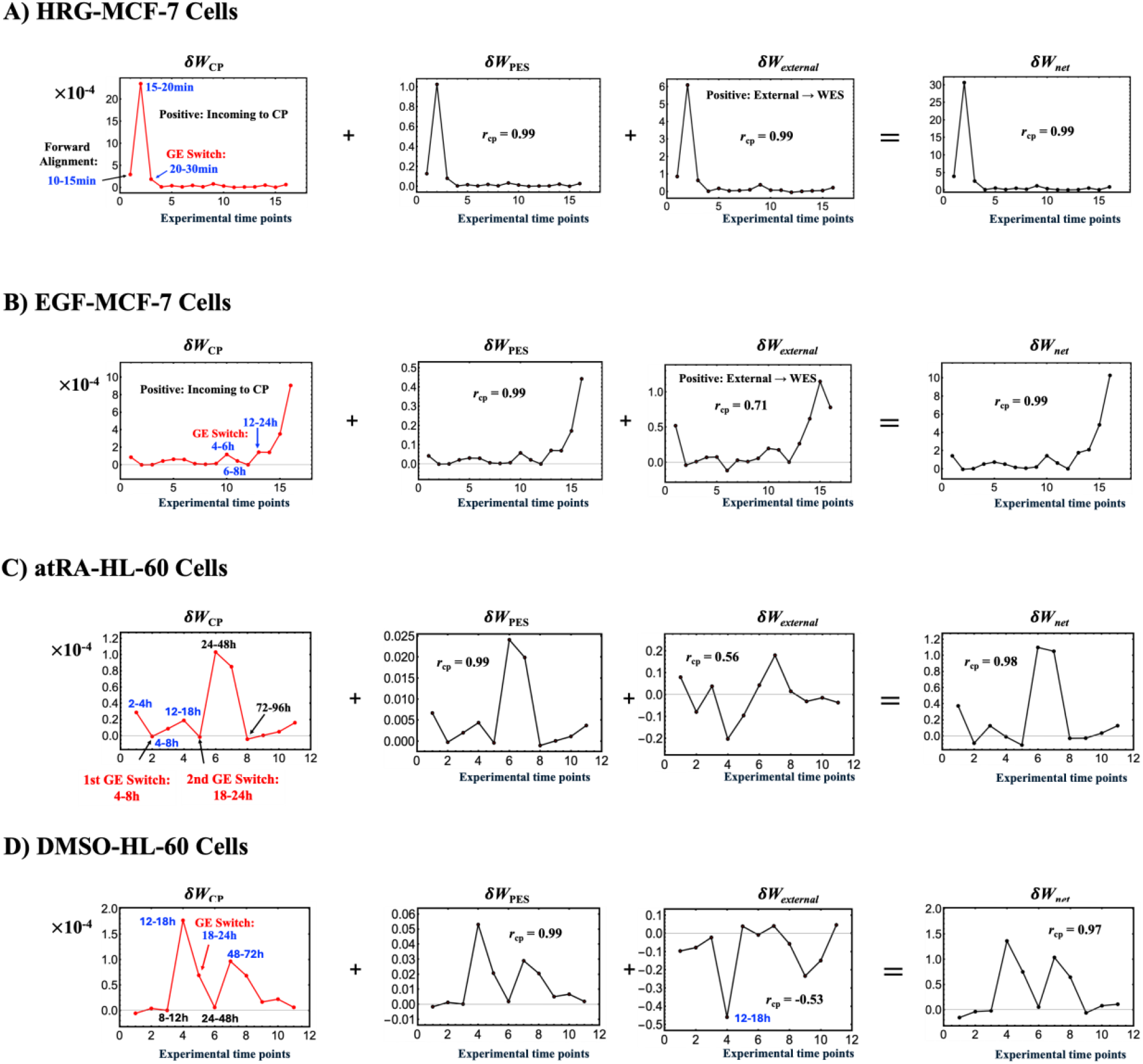
Unified temporal dynamics of EED work (δ*W*) and the Maxwell’s demon (MD) cycle. (A) HRG- and (B) EGF-stimulated MCF-7 cells; (C) atRA- and (D) DMSO-stimulated HL-60 cells. All *y*-axis values of δ*W* are shown with a multiplier of 10^-4^. Black text: *δW*timing; blue text: MD-cycle time window; red text: genome engine (GE) switch time window (Table 3; Section 2.4.5). The *x*-axis shows experimental time-point intervals. EED work over [*t*_j_, *t*_j+1_] is calculated from the self-flux at *t_j_*, evaluated from [*t*_j-1_, *t*_j_*, t*_j+1_], and the forward OUT flux over [*t*_j_, *t*_j+1_]. Net EED work is the sum of the CP, PES, and external contributions: δ*W*_CP_, δ*W*_PES_, δ*W*_external_, giving δ*W*_net_. The correlation coefficients in the PES, external, and net EED work panels are calculated relative to the CP EED work; the PES correlations in all four conditions show clear CP-PES phase synchronization, while net EED work also scales well with CP EED work. Notably, the magnitude of δ*W*_CP_ generally far exceeds that of δ*W*_PES_ in all four conditions, showing that CP provides the dominant internal work contribution. **(A) HRG-MCF-7 (fast actuation)**: During the MD cycle, δ*W*_CP_ exhibits a sharp positive peak at 15-20 min, within the Measurement phase, followed by a rapid decline during the 20-30 min Feedback interval, where the GE switch occurs. Furthermore, δ*W*_external_ and δ*W*_net_ are both almost perfectly correlated with δ*W*_CP_, consistent with CP-dominated EED work dynamics in the HRG condition. **(B) EGF-MCF-7 (cell proliferation)**: During the MD cycle, a local δ*W*_CP_ peak occurs during the 4–6 h Preparation phase, where the GE switch occurs, and is about 20-fold smaller than the HRG peak. δ*W*_CP_ then approaches zero during the Measurement phase and increases during the 12-24 h Feedback interval. After the MD cycle, δ*W*_CP_ increases sharply in the post-cycle period (> 24 h), suggesting increased CP EED work during the later cell proliferative response (Section 2.4.5). **(C) atRA-HL-60 (delayed actuation):** δ*W*_CP_ decreases from the 2-4 h Preparation phase to near zero during the 4-8 h Measurement phase, then increases during the 12-18 h Feedback phase, consistent with CP-braking dynamics (**Figure 7A**). After the MD cycle, δ*W*_CP_ becomes negative at 18-24 h, where the second genome engine switch occurs, then rises sharply to a peak at 24-48 h and falls again to negative values at 72-96 h and then gradually increases. This reflects delayed EED work execution after the MD cycle (Section 2.4.5). δ*W*_net_ strongly correlates with δ*W*_CP_, showing CP-dominated net-work dynamics. (D) DMSO-HL-60 (intermediate actuation): δ*W*_CP_ rises sharply after 8-12 h to its largest peak during the 12-18 h Preparation phase. It then decreases sharply but remains positive during the 18-24 h Measurement phase, where the GE switch occurs, and approaches zero at 24-48 h. δ*W*_CP_ then rises again to a second peak during the 48-72 h Feedback phase. External EED work is predominantly negative during the Preparation phase (12-18 h) and partially offsets the positive CP contribution. *δW*_net_nevertheless closely tracks the CP trajectory.

#### 2.4.5 The Irreversible Time Arrow of Cancer Cell-Fate Change: Three Timelines Defining the Thermodynamic Commitment Window

We examine fate commitment by comparing three distinct timelines: (1) the Maxwell’s demon (MD) cycle, (2) the information-actuation timeline defined by CP effective expression-derived (EED) work coupled to net mutual-information change, and (3) the genome-engine (GE) switch timeline. The MD cycle defines the Preparation, Measurement, and Feedback phases. The information-actuation timeline tracks the temporal coupling between CP EED work and net mutual-information change. The GE switch marks coordinated reorganization of the genome-engine network. Fate commitment is identified from the condition-specific temporal relati cell fate onships among these three timelines. Based on these three timelines, we propose four criteria for the irreversible time arrow of cell-fate change.

Hereafter, EED work is termed simply work because the analysis is entirely within expression space, unless the EED designation is needed for clarity.

The first two timelines are established by the following findings:

**(1) CP as the dominant internal EED-work contributor through infromation-EED work coupling. Figure 12** shows that, within the EED-work space, CP and PES exhibit clear phase synchronization with δ*W*_CP_ ≫ δ*W*_PE*S*_ and that net work closely tracks the CP work. Notably, this CP-PES work synchronization is consistent with CP-PES thermodynamic phase synchronization (Section 2.3.3), reflecting their dynamical-thermodynamic coupling in the coarse-grained WES representation. These relations identify the CP as the dominant internal work contributor. Furthermore, as the leading term, CP work is coupled to net mutual-information change, Δ*I*_net_(CP, PES), defining information-work coupling (Eq. 15).
**(2) CP as the genomic Maxwell’s demon (MD) in expression space**. The expression-derived net mutual information *I*_net_(CP, PES), is the order parameter of CP-PES thermodynamic coupling. Its temporal change, Δ*I*_net_(CP, PES), maps the Preparation, Measurement, and Feedback phases of the MD cycle and is phase-aligned with Δ*S*(CP) (**Figures 4-5**). Along the chromatin-remodeling proxy coordinate, *ln*(*nrmsf*), CP-associated chromatin-state rewriting occurs through CP-enhancing and CP-braking modes (**Figures 6-7**). In the CP-enhancing mode, CP bimodal formation and reset can support switchable ON/OFF chromatin-state reconfiguration; in the CP-braking mode, CP-dominated execution is delayed until after the MD cycle.

Together, these findings suggest that the CP functions operationally as the genomic Maxwell’s demon through information-work coupling and mode-specific chromatin-state rewriting. The possible correspondence to Landauer erasure and the physical scope of this framework are discussed in Section 3.1.

The third timeline is the GE switch. It is determined from the fluctuation of each interaction flux around its temporal mean:

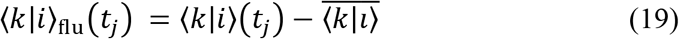

The temporal mean is taken over all time points at which the interaction flux is defined. Coordinated sign reversals among interaction-flux fluctuations locate the GE switch within [*t*_j_, *t*_j+1_].

**Figure 13** identifies the GE switch as reorganization of interaction fluxes among the CP, Super, and Sub domains. This is important because it marks when the genome-engine network changes its coordinated state, allowing the GE-switch timeline to be aligned with CP work and information dynamics in defining the commitment window. In HRG, EGF, and DMSO, this GE-switch reorganization occurs within the same CP-driven phase architecture identified in **Figure 11**, linking the dynamical GE switch to the CP thermodynamic dynamics. atRA deviates from this architecture through mixed-phase CP-braking dynamics and delayed CP-GA dynamical phase synchronization.

Therefore, these findings link the information-thermodynamic and dynamical roles of the CP to the corresponding net-work mode (**Section 2.4.4**) and subsequent condition-specific transition, as follows.

**(i) HRG condition (rapid fate commitment and Feedback execution**). In MCF-7 cells, HRG and EGF activate distinct receptors within the shared ErbB signaling network. HRG induces sustained cytoplasmic ERK signaling and stabilization of c-Fos protein associated with differentiation, whereas EGF induces transient cytoplasmic ERK signaling associated with proliferation [Nagashima et al., 2007; Nakakuki et al., 2010].

Under the HRG condition, all-or-none (biphasic) genome-wide expression dynamics emerge around 10-20 min along the nrmsf coordinate [Tsuchiya et al., 2014]. Chromatin structural reorganization occurs over the same 15-30 min interval [Krigerts et al., 2021]. By about 20 min, the ligand-specific signaling divergence also becomes evident in biphasic c-Fos protein induction [Nagashima et al., 2007].

**Figure 12A** shows that, under HRG, *δW*_PES_ and *δW*_external_ are phase-correlated with *δW*_CP,_ while *δW*_net_, closely tracks *δW*_CP_ with near-perfect Pearson correlations. During the 15-20 min Measurement phase, *δW*_CP_ shows a sharp positive peak, indicating incoming work to the CP. Positive external work further increases net work, producing a strong positive net-work peak. In the same interval, Δ*I*_net_(CP, PES) reaches its maximum positive value within the HRG MD cycle, indicating peak active information gain (**Figure 4A**). Strong CP-GA dynamical phase synchronization spans the Measurement phase and the transition into Feedback (**Figure 10A**).

##### Fate-guiding critical transition initiated at 20 min

**Figure 6A** identifies the 20-30 min Feedback phase as the fate-guiding critical-transition interval, initiated at the 20 min boundary following the 15-20 min Measurement phase. Across this boundary, Δ*I*_net_(CP, PES) changes from positive during Measurement to negative during Feedback. During 20-30 min Feedback, the GE switch occurs, while Δ*S*(CP) < 0 and Δ*S*(WES) < 0 indicate genome-wide ordering (**Figures 6A, 13A**). Strong CP-GA dynamical phase synchronization spans the Measurement-to-Feedback transition (**Figure 10A**).

Thus, the 15-20 min Measurement phase defines the HRG commitment window, and the critical transition initiated at the 20 min boundary marks the transition into synchronized Feedback execution.

This Measurement-to-Feedback transition is temporally aligned with independent protein and structural observations: biphasic c-Fos protein after about 20 min [Nagashima et al., 2007] and PAD/chromatin opening from 15-20 min, with the unraveled PAD state persisting to 30 min [Krigerts et al., 2021]. For further details, see Section 2.3.2 for structural correspondence and Section 3.1 for physical interpretation.

Regarding reproducibility, the two replicates (Rep1 and Rep 2) [Saeki et al., 2009] reproduce the same fate-commitment dynamics: (1) the CP-work response across the 15-30 min MD transition and (2) the GE switch during 20-30 min Feedback. Thus, the 15-20 min commitment window and subsequent Feedback execution are reproducible across replicates (**Supplementary File S1**).

**(ii) EGF condition (no fate commitment with proliferation)**. EGF activates EGFR/ErbB1, producing transient cytoplasmic ERK signaling associated with proliferation [Nagashima et al., 2007; Nakakuki et al., 2010]. EGF also fails to establish the HRG-specific late-transcriptional regulatory network, which contributes to ERK feedback control and irreversible differentiation [Saeki et al., 2009].

EGF exibits CP-PES work synchronization and undergoes a GE switch during the 4-6 h Preparation phase together with a local positive CP-work peak (**Figures 12B and 13B**). Its magnitude is approximately 20-fold smaller than the corresponding HRG peak in MCF-7 cells. The MD cycle exhibits CP-GA dynamical synchronization (**Figure 10B**). In the 6-8 h Measurement phase, Δ*I*_net_(CP, PES), Δ*S*(CP), and Δ*S*(WES) increase (**Figure 4B**), but only to about one-fifth of the HRG amplitudes. After Measurement, the CP undergoes reset and reorganization (**Figure 6B**). During the 12-24 h Feedback phase, the commitment-to-execution signature is not established (see criterion (ii) below). Together, these findings show that information gain and positive CP work occur during the MD cycle but do not progress to fate commitment or execution. Thus, EGF exhibits a CP-enhancing mode without fate commitment or execution.

After 24 h, CP work increases substantially, whereas Δ*I*_net_(CP, PES) remains of the same order of magnitude as during the MD cycle. The post-24 h increase in CP work may reflect increased forward CP OUT flux associated with the EGF-induced proliferative response, which is evident by 24 h and sustained thereafter [Dong et al., 1992; Hopkins et al., 2016].

Regarding reproducibility, EGF Rep 1 and Rep 2 [Saeki et al., 2009] show the same non-commitment pattern: CP work rises at 4-6 h, whereas the information/entropy response peaks at 6-8 h, indicating that work and information/entropy amplification are not temporally coordinated (see proposed criteria (iv)). The pronounced post-CP-work responses in Rep 1 after 24 h and in Rep 2 at 4-6 h may reflect stronger proliferative activity rather than fate commitment [Nagashima et al., 2007] (**Supplementary File S1**).

**(iii) atRA condition (fate commitment following CP braking)**. In HL-60 cells, atRA induces granulocytic differentiation through RARα, a nuclear receptor-type transcription factor [Collins et al., 1990], yielding a more mature state with reduced proliferation [Ghonyan et al., 2025]. atRA-induced differentiation is accompanied by specific chromatin conformational changes that alter gene-regulatory interactions and are correlated with differentiation-associated gene expression [Li, Y. et al., 2018].

Two GE switches occur at 4-8 h during Measurement and at 18-24 h after the MD cycle. They define two distinct transition routes: CP-braked sensing and compaction without irreversible execution, and post-Feedback CP-enhancing actuation leading to fate commitment.

**(a) Active sensing and the first transition under CP braking**. Positive incoming CP work during 2-4 h Preparation precedes active sensing. During 4-8 h Measurement, *δW*_CP_ approaches zero, while Δ*S*(CP), Δ*S*(PES), Δ*S*(WES) and Δ*I*_net_(CP, PES) reach their largest positive values under CP-PES thermodynamic synchronization (**Figure 5A**). Net work is negative, mainly because of the negative external contribution (**Figure 12C**).

During the 12-18 h Feedback phase, *δW*_CP_ becomes positive, but negative external work approximately offsets the positive internal contributions, leaving net work near zero (**Figure 12C**). At the same time, Δ*S*(CP) < 0 and Δ*S*(WES) < 0, indicating genome-wide ordering (**Figure 5A**).

Thus, genome-wide ordering occurs without positive net-driving work, and this transition does not establish commitment. A recent study [Ghonyan et al., 2025] suggests that reduced proliferation and a more mature granulocytic phenotype may be associated with the atRA-specific CP-braking trajectory.

**(b) Second critical transition at 48 h.** After the second GE switch during 18-24 h **(Figure 13C**), both *δW*_CP_and *δW*_net_rapidly rise during 24-48 h from near-zero and negative values, respectively (**Figure 12C**). During 24-48 h, Δ*I*_net_(CP; PES) reaches its second-largest positive value, identifying the second-largest information gain, and then becomes negative during 48-72 h (**Figure 5A**). This change coincides with *δW*_net_decreasing from its largest positive value during 24-48 h (**Figure 12C**). At the 48 h boundary, the second critical transition occurs together with strong CP-GA dynamical phase synchronization in the 24-48-72 h sequence (**Figure 10C**). During 48-72 h, Δ*S*(CP) < 0 and Δ*S*(WES) < 0 (**Figure 5A**), while CP incoming work and net work remain positive (Figure 12C), indicating genome-wide execution after resolution of CP braking with the CP bimodal profile resolving across the 48 h boundary (**Figure 7A**).

Thus, 24-48 h is the commitment window, and the critical transition at the 48 h boundary marks the transition into execution during 48-72 h.

**(iv) DMSO condition (measurement-phase fate commitment).** DMSO, a polar solvent inducer of differentiation, drives HL-60 cells from proliferation toward granulocytic differentiation through c-myc transcriptional repression accompanied by reorganization of its regulatory chromatin toward a more closed state [High et al., 1987; Siebenlist et al., 1988].

DMSO exhibits CP-PES work synchronization and shows its largest positive CP-work peak during the 12-18 h Preparation phase (**Figure 12D**). Negative external work partially offsets this contribution, but net work remains positive. During the 18-24 h Measurement phase, CP and net work remain positive, Δ*I*_net_(CP;PES) > 0 **(Figure 5B)**, the GE switch occurs (**Figure 13D**), and CP-GA dynamical synchronization emerges around 18 h (**Figure 10D**). Thus, 18-24 h defines the DMSO fate-commitment window.

During 24-48 h, CP and net work approach zero. Renewed positive CP and net work occur during the 48-72 h Feedback phase, accompanied by the fate-guiding critical transition and CP-GA dynamical synchronization. During this execution interval, Δ*I*_net_(CP;PES) < 0, and Δ*S*(CP) < 0 and Δ*S*(WES) < 0, indicating genome-wide ordering. Thus, DMSO separates Measurement-phase fate commitment from Feedback-phase execution.

Based on the four conditions, we propose four criteria for the irreversible time arrow of cell-fate change in expression space. Criteria (i) and (ii) describe signatures derived from measured gene expression; criterion (iii) states a conditional thermodynamic requirement; and criterion (iv) proposes an amplitude-coupling condition together with a mechanistic hypothesis that remains to be tested. These criteria are assessed together through the condition-specific sequence of commitment and execution.

**(i) CP-dominated net-driving commitment**: Positive (incoming) CP work, *δW*_CP_ > 0, positive CP-dominated net work, *δW*_net_ > 0, and net mutual-information gain, Δ*I*_net_(CP;PES) > 0, identify the commitment window, with the GE switch serving as a condition-specific temporal landmark and the subsequent execution sequence in criterion (ii) supporting the identification of commitment within the proposed framework.
**(ii) Dynamical execution**: Positive CP work, *δW*_CP_ > 0 and positive net work, *δW*_net_ > 0, accompany net mutual-information decrease, Δ*I*_net_(CP;PES) < 0, and genome-wide ordering, Δ*S*(CP) < 0 and Δ*S*(WES) < 0, at or after commitment. The timing of CP-GA synchronization is assessed across the commitment-to-execution sequence.
**(iii) Outward dissipation**: Under the thermodynamic mapping, with the WES defined as the system, when Δ*S*(WES) < 0, entropy export must exceed the entropy decrease within the WES, Δ*S*(OUT) > |Δ*S*(WES)|, to ensure positive total entropy production. This is a conditional requirement; entropy export is not directly measured here. We discuss its physical interpretation and the measurements needed to test this requirement in Section 3.1**(v)**.
**(iv) Coupled information-work amplification**. Within CP-PES synchronization windows, we hypothesize that fate commitment requires coordinated amplification of the synchronized information/entropy response and CP work within the commitment window identified by criterion (i) (**Figures 4A, 5, 12A,C,D**). This amplification may reflect stochastic-resonance-like behavior (**Note 3** in Section 2.4.4). Non-commitment is expected when information and work are not coordinately amplified (see EGF condition): information gain occurs without positive net-driving work, or work increases without a corresponding information/entropy amplification (**Figures 4B,12B** for EGF; **Figures 5A, 12C** for the first non-committing atRA transition at 12-18 h).

In summary, these condition-specific windows constitute “time-gated rules” for fate control: the relative timing of the three timelines, rather than any single information change, work peak, or GE-switching event, defines when the system progresses from commitment to execution.

**Figure 13.**
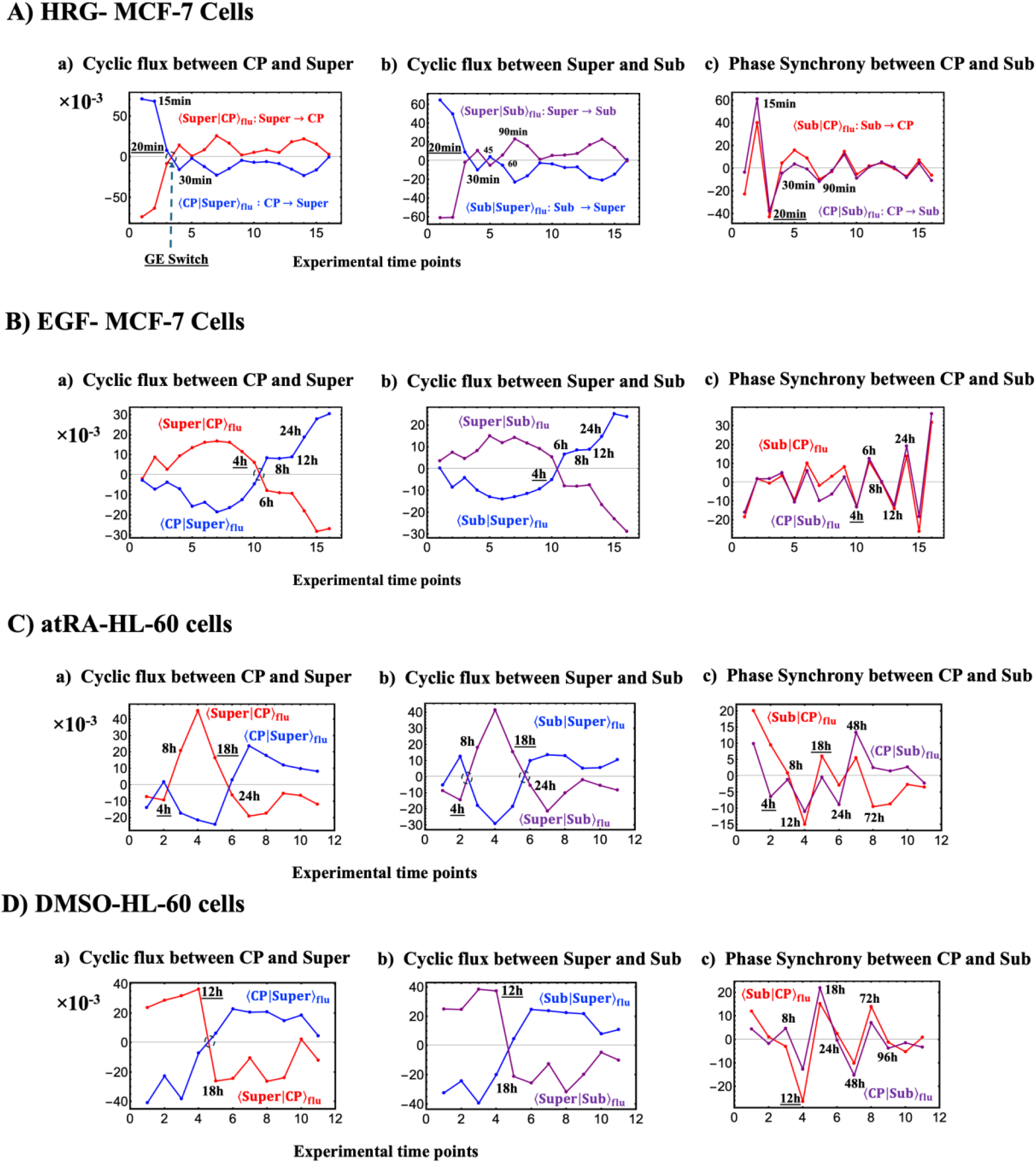
Fluctuation dynamics of interaction flux for MCF-7 and HL-60 cancer cells: cyclic flux and phase synchronization. **A)** HRG- and **B)** EGF-stimulated MCF-7 cells; **C)** atRA- and **D)** DMSO-stimulated HL-60 cells. The *x*-axis is expressed in discrete experimental time steps, with Δ*t* = 1 between consecutive measurements. Interaction flux fluctuation dynamics of the CP, Super, and Sub domains are shown relative to their temporal average baselines. Cyclic flux arises a) between the CP and Super domains and b) between the Super and Sub domains through anti-phase domain fluctuation dynamics, while panel c) shows phase synchronization between the CP and Sub domains. Interaction flux at *t*_j_ is evaluated within a three-time-point window [*t*_j-1_, *t*_j_, *t*_j+1_] (see main text), where *t*_j_, indicated by black text, denotes the corresponding actual experimental time point. All *y*-axis values are multiplied by 10^-3^, with actual signal levels of order *O*(10^-2^).

The summary of timings together with the GE switch, the CP-Maxwell’s demon (MD) cycle, and the peak CP work (see main text and **Table 3** for details) is as follows:

**A) HRG (CP-enhancing mode):** The GE switch is initiated at 20 min and occurs before 30 min; thus, the GE switch occurs at 20-30 min, matching the Feedback phase (**Figure 6A**) and following the sharp positive CP-work peak during the preceding 15-20 min Measurement phase (**Figure 12A**).
**B) EGF (CP-enhancing mode):** The GE switch occurs at 4-6 h, matching the Preparation phase (**Figure 6B**). This GE switch marks the onset of non-cell-fate-changing MD actuation. A much weaker local positive incoming CP-work peak than in HRG occurs during the same 4-6 h Preparation phase (**Figure 12B**).
**C) atRA (CP-braking mode):** Two GE switches occur at the 4-8 h and 18-24 h windows. These timings correspond to the Measurement phase and the post-MD-cycle period, respectively (**Figure 7A**).
**D) DMSO (CP-enhancing mode):** The GE switch occurs at 18-24 h, matching the Measurement phase (**Figure 7B**) and following the peak incoming CP work during the preceding 12-18 h Preparation phase (**Figure 12D**).

### 3.1 Physical Interpretation and Scope of the Genomic Maxwell’s Demon Framework within SOC-Controlled Genome Dynamics

Our previous studies have progressively extended our understanding of SOC-controlled genome dynamics across cell-fate transitions in cancer cell populations and T helper 17 differentiation, and at the single-cell level in mouse and human embryonic development [Tsuchiya et al., 2016, 2020]. The present study further develops the genomic Maxwell’s demon (MD) framework [Tsuchiya et al., 2025] under SOC-controlled genome dynamics by integrating information thermodynamics with effective expression-derived (EED) dynamics across HRG, DMSO, atRA, and EGF.

The following five points distinguish observed results, proposed physical mechanisms, and the measurements needed to test them: (i) the physical interpretation of transcriptomic information measures, (ii) the meaning and calibration of dimensionless work, (iii) the possible DNA-structural bridge to dynamics, (iv) the distinction between operational genome memory and Landauer erasure, and (v) the conditions linking commitment signatures to thermodynamic irreversibility.

**(i) Information-thermodynamic quantities and SOC: their physical interpretation.** Shannon entropy and mutual information enter formally into stochastic thermodynamics when the physical states, dynamics, and environmental exchanges are specified. The Preparation, Measurement, and Feedback phases defined here provide an operational cycle consistent with the measurement and feedback framework of information thermodynamics [Sagawa and Ueda, 2012; Parrondo et al., 2015]. Thus, the present information-thermodynamic quantities are experimentally grounded in transcriptomic dynamics, whereas their physical interpretation requires identifying the material states underlying these dynamics.

**1) *ln*(*nrmsf*) as a chromatin-remodeling proxy**. Chromatin organization provides a possible physical basis for the SOC-controlled genome dynamics. Zimatore et al. (2021) used PCA to statistically validate the quantitative relationship between principal-component scores and *ln*(*nrmsf*). RQA showed that the number of co-regulated gene pairs decreases exponentially with their separation along chromosomes, supporting local chromatin organization as a structural constraint on coordinated gene regulation. Together with the temporal correspondence between gene-expression dynamics and PAD remodeling in HRG-stimulated MCF-7 cells [Krigerts et al. 2021], these findings statistically validate the role of chromatin remodeling in SOC control of genome expression and support *ln*(*nrmsf*) as a quantitative expression-derived proxy for coarse-grained chromatin remodeling (Section 2.3.2).
**2) Scale-free PAD organization and chromatin structural evidence in HRG.** In MCF-7 cells, Krigerts et al. (2021) experimentally showed a scale-free relationship between PAD number and size, indicating the absence of a characteristic PAD size. Under HRG stimulation, PAD remodeling and chromatin/DNA unfolding occur during the 15-20 min window. Thus, scale-free PAD organization provides chromatin structural evidence consistent with SOC, while the time-matched HRG structural transition provides chromatin structural correspondence with the transcriptome-derived critical transition (Section 2.4.5). Equivalent structural validation is still needed for the other conditions.
**3) Calibration to physical thermodynamic quantities at constant temperature**. These structural correspondences provide a physical basis for connecting SOC-controlled transcriptomic dynamics with chromatin organization. Extending this framework to physical thermodynamics requires quantitative calibration of the associated dynamics and energy exchanges. The cell cultures are maintained at *T* = 310.15 K, with the incubator treated as an external heat bath (Section 2.1). Quantitative physical calibration would require time-resolved mapping of EED force and displacement onto measurable chromatin/DNA force-displacement dynamics, together with measurements of the associated energy exchanges, as discussed in (ii) and (iii) below.
**(ii) EED Work, dimensionless units, and energetic calibration.**

**1) Information-to-EED work coupling and condition-specific dynamics**. EED work (Section 2.4.4) is defined in expression space from the EED force acting on a domain attractor and its forward displacement. As established in Section 2.4.1, the EED force-entropy coupling supports the use of the forward OUT flux displacement to define EED work. Together with CP-PES thermodynamic phase synchronization, this coupling links Δ*I*_net_(CP;PES) to forward EED work (Eqs. 14-15). This provides the operational basis for information-to-work conversion during the fate-guiding critical transition in expression space (Section 2.4.5). Across the four conditions, the sign and timing of EED work and this information-work coupling distinguish CP-enhancing dynamics in HRG and DMSO, CP-braking followed by positive-work execution in atRA, and the weaker non-committing EGF response during the MD cycle. CP-dominated execution occurs during the MD Feedback phase in HRG and DMSO and after the MD cycle in atRA.
**2) Mechanical interpretation and energetic calibration of EED work**. EED work is dimensionless under the adopted normalization with unit mass (*m* = 1) and unit time step (Δ*t* = 1) (Section 2.4.1). Its interpretation as physical mechanical work requires energetic calibration against measurable chromatin/DNA force-displacement dynamics. Physical mechanical work can be obtained by integrating force over extension in single-molecule measurements of chromatin deformation, nucleosome unfolding, and DNA unwrapping [Cui & Bustamante, 2000; Bennink et al., 2001; Brower-Toland et al., 2002; Kruithof et al., 2009]. Converting EED work into mechanical energy units therefore requires a quantitative mapping between expression-space force-displacement trajectories and measurable chromatin/DNA force-displacement trajectories. In HRG, the time-matched chromatin/DNA changes discussed in point (i) provide structural support for such a connection, although quantitative energetic calibration remains to be established. The correspondence between CP-state rewriting and microscopic chromatin memory remains to be established experimentally. Likewise, the correspondence between CP-state rewriting and microscopic chromatin memory remains to be established experimentally.
**(iii) Possible physical bridge between EED work and DNA structural dynamics**. The DNA structural-scaling model links collective DNA open-state dynamics to mechanically influenced transitions across bistable, metastable, and critical regimes [Naimark, 2007; Nikitiuk et al., 2022, 2025]. Mechanical force enters the internal field acting on DNA open states and couples to their structural displacement, providing a possible physical counterpart to the force-displacement relation underlying EED work. Relating these structural states to measured chromatin conformations and EED trajectories would make this proposed connection quantitatively testable. Analysis of stochastic cellular dynamics using laser microscopy could complement this comparison. The broader mechanobiological framework is discussed in Section 3.2.
**(iv) Operational memory and its distinction from Landauer erasure**

**1) Operational memory and its thermodynamic signature.** “Rewritable chromatin memory” is defined operationally by CP-associated state rewriting synchronized with the whole expression system (WES). CP-enhancing switching occurs in HRG, DMSO, and EGF, whereas atRA exhibits a CP-braking mode with delayed execution. **Figures 4-5** show that the entropy changes, Δ*S*(CP) and Δ*S*(WES), together with the change in net mutual information, Δ*I*_net_(CP;PES), scale with Δ*S*(CP), consistent with CP-PES thermodynamic phase synchronization (Section 2.4.5). Furthermore, CP-PES phase synchronization occurs in EED work, where CP work dominates PES work and net work tracks CP work (**Figure 12**), and CP-GA dynamical phase synchronization (**Figure 10**). These results indicate that “rewritable memory” reflects CP-driven coordinated state changes across the WES in expression space.
**2) Bistable state rewriting and its possible structural basis.** Along the *ln*(*nrmsf*) coordinate, the MD double-well potential profile represents the inferred switchable bistable chromatin states in HRG, EGF, and DMSO, characterized by coherent ON/OFF Δ*S*(WES) profiles and transient CP-bimodal formation and reset (**Figures 6-7**). Here, *ln*(*nrmsf*) serves as a coarse-grained expression-derived proxy for chromatin remodeling. CP-associated bistable state rewriting and reinitialization provide the operational memory property within the Maxwell’s demon (MD) cycle. atRA shows no coherent bistable switching during the MD cycle. A possible structural basis for this operational memory is provided by bistable DNA dynamics. Experiments demonstrate hysteresis in DNA folding and unfolding [Shew et al., 2007], a bimodal nucleosome-density distribution in reconstituted chromatin [Nakai et al., 2005], and transcriptional ON/OFF switching associated with DNA conformational transitions [Tsuji and Yoshikawa, 2010]. These findings support a possible memory mechanism in which structural states retain information about prior conditions through hysteresis and influence transcriptional activity. Their direct connection to CP-associated memory remains to be established. Further DNA/chromatin structural mechanisms are discussed in Section 3.2.
**3) Initial-state loss and CP reset.** Two processes should be distinguished in relation to memory loss and rewriting:

- **Loss of the initial CP state.** Previous expression flux analysis (EFA) studies identified loss of the initial-state CP-centered sandpile-type critical organization in cancer cell populations and at the single-cell level in T helper 17 differentiation, and mouse and human embryonic development [Tsuchiya et al., 2016, 2020] (Section 2.4.1). This loss provides a coarse-grained signature of prior-state information loss.
- **CP reset.** CP reset denotes reconfiguration and reinitialization of the CP bimodal expression state. **Figures 6-7** show that the re-established CP bimodal state. Within the MD framework [Tsuchiya et al. 2025], this corresponds to a reconfigured expression-derived bistable ON/OFF chromatin-state landscape characterized by a CP-driven MD double-well potential profile. Thus, CP reset represents operational rewriting of the CP-associated memory state.
**4) Potential relation of initial-state loss and CP reset to Landauer erasure**. Initial-state loss and CP reset provide operational signatures of information loss and memory rewriting, respectively. Landauer erasure concerns a logically irreversible reset that maps distinct stored records to a common logical state [Landauer, 1961]. Establishing their relation to Landauer erasure requires identifying the physical states underlying the proposed chromatin memory, demonstrating a reset that maps distinct stored records to a common logical state, and establishing the associated energetic costs [Sagawa and Ueda, 2009; Parrondo et al., 2015]. The observed information-work coupling (Section 2.4.4) may provide a basis for examining these energetic costs once EED work is calibrated to physical energy.
**(v) Commitment signatures and thermodynamic irreversibility.**

**1) Commitment and execution signatures**. Entropy and information changes, under CP-PES thermodynamic phase synchronization, CP and net EED work, dynamical phase synchronization between the CP and the genome attractor (GA), and GE switching jointly identify commitment and execution windows within the proposed framework (Section 2.4.5). They do not directly measure physical entropy production or outward entropy export.
**2) Thermodynamic irreversibility and experimental tests.** Under the thermodynamic mapping assumed in Section 2.4.5, the requirement of positive total entropy production, together with the entropy balance in Eq 20, implies Δ*S*(OUT) > |Δ*S*(WES)| when Δ*S*(WES) < 0. Here, Δ*S*(OUT) denotes net outward entropy export. This relation provides a conditional lower bound on the entropy export required by the assumed thermodynamic mapping; its physical magnitude remains uncalibrated.

The identified windows guide tests of whether EED actuation is physically coupled to dissipation and outward entropy export. In HRG, matched expression and chromatin measurements at 10, 15, 20, and 30 min can test structural reorganization across the MD cycle, building on the structural observations of Krigerts et al. (2021). Equivalent measurements in DMSO, atRA, and EGF would test their distinct responses. Heat and metabolic-flux measurements, combined with a calibrated energy and entropy balance, could test entropy production and the proposed export bounds.

### 3.2 Higher-Order Chromatin and DNA Structural Dynamics as the Physical Basis for Genomic Control

Our study demonstrates that time-series transcriptomes encode time-gated control rules for fate change. However, although transcriptomes are sufficient to infer time-resolved control rules, they represent a many-to-one projection of the underlying highly ordered chromatin/DNA structure, so expression observables do not uniquely specify the memory-bearing structural state. At the coarse-grained level, this microscopic degeneracy is integrated over, allowing the collective-state dynamics relevant to genome control to be retained in the transcriptomic description. A complementary structural description is therefore needed to identify the physical basis of these collective states and the state variable that stores drive and implements switching.

DNA open-state dynamics provides such a structural coordinate. In a mesoscopic description of DNA open states, regulation is formulated in terms of an open-complex order parameter that reveals a specific type of criticality, namely structural-scaling transitions. The behavior of the open-complex ensemble is governed by a statistically predicted nonlinearity in the out-of-equilibrium free energy (representing the “epigenetic landscape” of an open-complex order parameter) and by an evolution equation in a generalized Ginzburg-Landau form [Naimark, 2007; Naimark et al., 2020a; Naimark et al., 2020b]. Two structural critical points were established in terms of the structural-scaling parameter *χ*, which plays the role of an effective temperature for the out-of-equilibrium DNA ensemble by setting its susceptibility to reorganize. Critical values of *χ* separate three regimes with distinct spatiotemporal dynamics, each associated with a distinct set of open-complex collective modes that correspond to self-similar solutions of the evolution equation. These modes can be associated with sets of attractor coordinates that encode memory of the current value of *χ* and support qualitatively different Maxwell’s demon scenarios [Naimark et al., 2020a; Nikitiuk et al., 2025] across the attractor set. At the transcriptomic level, *ln*(*nrmsf*) may serve as a counterpart of χ, providing an effective-temperature-like control axis for out-of-equilibrium chromatin organization, along which the CP region separates the Super and Sub domains, analogous to the critical values of χ that separate distinct dynamical regimes.

The pathway across the attractor set is characterized by independent “coarse-grained (mesoscopic) statistics” of the structural-scaling parameter, a characteristic feature of systems exhibiting so-called “slow dynamics.” The combination of generalized Boltzmann-Gibbs statistics with “coarse-grained statistics”, that is, superstatistics [Naimark, 2007], can be considered a statistical interpretation of “information thermodynamics.” The properties of the coarse-grained statistics, together with the nature of the critical points and the types of collective modes in an open complex system, provide a duality in the nature of the Maxwell’s demon, associated with the thermodynamic scenario of metastability decomposition along the CP pathway and the corresponding superstatistics. This superstatistics can be considered a key factor underlying slow dynamics, providing a qualitatively different scenario for epigenetic plasticity and the transition toward “epigenetic fragility” as a potential precursor of cancer.

The evolutionary path along the free-energy “epigenetic landscape” and the corresponding Maxwell’s demon scenario are coupled to slow mRNA expression dynamics, mediated by slow collective-mode dynamics as coherent mRNA expression modes. Consistent with these results, the critical dynamics of gene expression exhibits collective-mode-induced scaling, with attractor properties characteristic of the corresponding “epigenetic landscape”. The qualitative diversity of paths along the “epigenetic landscape” is determined by the current mean and standard deviation of *χ*, which characterize the corresponding domain-attractor region in chromatin phase separation. At a complementary polymer scale, coarse-grained modeling shows that heterogeneous chromatin interactions can generate dynamic three-dimensional genome organization through phase separation [Fujishiro and Sasai, 2022].

A more concrete genome-wide switching mechanism is the coil-compact transition of giant genomic DNA, in which transcription is active in the elongated coiled state and suppressed in the compact globule state [Nakai et al., 2005; Luckel et al., 2005; Takenaka et al., 2008; Nagahara et al., 2010; Tsuji et al., 2010; Nishio et al., 2021; Nishio et al., 2022; Ogawa et al, 2026]. A global transition in genome-sized DNA can therefore provide a physical mechanism for coordinating the expression of many genes and may underlie genome-wide reorganization during the Measurement-to-Feedback transition. Genome-sized DNA exhibits the characteristics of a semiflexible polymer whose conformational transitions exhibit strong hysteresis. DNA supercoiling can also generate entropy-stabilized coexistence of multiple plectonemic states, providing a complementary topological mechanism for metastable DNA structural switching [Segers et al., 2025]. Such intrinsic properties generate metastable conformations and direction-dependent switching, providing a physical basis for non-genetic memory and commitment [Yoshinaga et al., 2005].

These properties motivate coarse-grained stochastic models of slow genomic control, where a structural order parameter coupled to an effective environment variable is described by nonlinear Langevin-type dynamics that link microscopic noise to robust macroscopic switching and oscillation [Nagahara and Yoshikawa, 2010; Gillespie, 2000; van Kampen, 2007; Gardiner, 2009]. Finally, field-based models of DNA opening extend this view to explicit open-complex (“breathing bubble”) dynamics, connecting structural-scaling transitions to time-gated genome-wide reorganization [Naimark et al., 2020a; Nikitiuk et al., 2025].

## 4. Conclusion: The Genome as a Unified Information-Work Engine

We identify CP-driven SOC control as the organizing biophysical principle linking information acquisition and effective expression-derived (EED) work to genome-state transitions, cell-fate commitment, and execution. Within the present coarse-grained description, collective expression dynamics reveal this system-level control without explicit tracking of underlying molecular complexity. This principle unifies information thermodynamics and genome-engine dynamics within a single open, non-equilibrium whole expression system (WES), which operates as an information-work engine.

The critical point (CP) gene ensemble performs two distinct but integrated roles: an information-thermodynamic role and a dynamical role. As a genomic Maxwell’s demon (MD) operator, the CP links information acquisition to feedback-associated state rewriting; as an SOC controller, it coordinates genome-wide critical transitions through CP-GA synchronization and CP-dominated EED work. CP-PES thermodynamic phase synchronization and information-work coupling integrate these roles into a unified control process.

**(i) Temporal Control of Cancer Cell-Fate Commitment**

The analysis integrates three timelines: the MD cycle, CP EED work coupled to net mutual-information change, and GE switching, together with CP-GA synchronization. The three differentiation-inducing conditions realize this architecture through distinct temporal modes, while EGF provides the non-fate comparison:

- **HRG:** Maximal information gain and CP-dominated work coincide with commitment, followed by genome-wide execution.
- **DMSO:** A pronounced CP-work response precedes commitment, separating work magnitude from commitment timing.
- **atRA:** CP braking permits information acquisition while suppressing net forward work before commitment and execution.
- **EGF:** GE switching and phase synchronization occur without fate commitment; CP-work amplification is temporally separated from information and entropy amplification.

These comparisons distinguish genome-wide critical activity from fate commitment itself. Fate commitment is identified when information and entropy responses couple to CP-dominated net EED work within CP-PES synchronization windows. This forms a coordinated sequence leading to genome-wide execution. These condition-specific windows constitute time-gated rules for fate control: the relative timing of the three timelines, rather than any single information change, work peak, or GE switching event, defines progression from commitment to execution.

**(ii) Operational Rewritable Chromatin Memory and Irreversibility**

CP-PES synchronization in thermodynamics and EED work, together with CP-GA dynamical synchronization, provides the operational basis for rewritable chromatin memory through CP bimodal-state formation, reconfiguration, and reset. In the CP-enhancing mode, information dynamics couple to forward EED work; in the CP-braking mode, information acquisition can proceed while net forward work is suppressed. These operating modes are distinct from the two states of CP bimodality.

Operational memory captures the temporal separation between information acquisition and genome-wide execution. Independent experimental observations under HRG stimulation support correspondence with chromatin structural remodeling. Future structural studies under DMSO, atRA, and EGF could test this correspondence. Energetic calibration would enable quantitative comparison of EED work with physical mechanical work.

We propose four criteria for the irreversible arrow of cell-fate change in expression space (Section 2.4.5). In particular, criterion (iii) requires outward entropy export exceeding the magnitude of the WES entropy decrease to ensure positive total entropy production. Physical entropy calibration and direct measurement of outward entropy export would enable testing of this conditional thermodynamic requirement.

**(iii) Therapeutic Implications: Testable Time-Gated Interventions**

The time-gated rules extend intervention strategies by specifying not only which regulatory processes to target but also when to perturb them within the information-work sequence. Two complementary strategies follow:

- **Modulate CP-driven EED work through time-gated perturbation.** Identifying epigenetic and chromatin regulators associated with CP EED work would allow perturbations to be aligned with relevant MD-cycle and commitment windows, testing whether altered CP-driven work changes progression from information acquisition to commitment and execution.
- **Exploit the endogenous CP-braking mode.** The atRA response shows that information acquisition can proceed while net forward work remains suppressed. Identifying the molecular basis of this braking mode could provide a complementary strategy for controlling when CP-driven work and GE switching enter the commitment-to-execution sequence.

This framework integrates information processing, operational memory, and EED work through CP-driven SOC control of genome-wide fate transitions, defining testable time-gated intervention windows targeting CP actuation and braking.

## 5. Materials

i. Microarray data for the activation of ErbB receptor ligands in human breast cancer MCF-7 cells by EGF and HRG; Gene Expression Omnibus (GEO) ID: GSE13009 (N = 22,277 mRNAs; for experimental details, see [Saeki et al., 2009]) at 18 time points: *t*_1_ = 0, *t*_2_ = 10, 15, 20, 30, 45, 60, 90 min; and 2, 3, 4, 6, 8, 12, 24, 36, 48, *t*_18_ = 72 h. Each condition includes two replicates (rep 1 and rep 2). The analyses in this report use replicates 1 and 2 for both EGF and HRG (replicate 2 results are provided in **Supplemental File 1**).
ii. Microarray data for terminal differentiation induced in human leukemia HL-60 cells by DMSO and atRA; GEO ID: GSE14500 (N = 12,625 mRNAs; for details, see [Huang et al., 2005]) at 13 time points: *t*_1_= 0, *t*_2_ = 2, 4, 8, 12, 18, 24, 48, 72, 96, 120, 144, *t*_13_ =168 h.

The robust multichip average (RMA) method was used for background adjustment and normalization of the expression data and to reduce false positives [Bolstad et al., 2003; Irizarry et al., 2003; McClintick et al., 2006].

## Abbreviations (biology-related terms)

atRA: all-trans retinoic acid
C/EBPα: CCAAT/enhancer-binding protein alpha
c-Fos: cellular Fos proto-oncogene protein
DMSO: dimethyl sulfoxide
EGF: epidermal growth factor
EGFR (ERBB1): epidermal growth factor receptor
ErbB3 (HER3): human epidermal growth factor receptor 3
ErbB4 (HER4): human epidermal growth factor receptor 4
ERK: extracellular signal-regulated kinase (MAPK)
HER2: Human Epidermal Growth Factor Receptor 2
HL-60: human acute promyelocytic leukemia cell line
HRG: heregulin
MCF-7: human breast cancer cell line
PADs: pericentromere-associated domains
RARα: retinoic acid receptor alpha

## Abbreviations (physics-related terms)

CM: center of mass
CSA: chromatin state analysis
CSB: coherent stochastic behavior
CP: critical point
DEA: domain ensemble-average
EFA: expression flux analysis
EED: effective expression-derived
GE: genome engine
ITA: information-thermodynamic analysis
ln(nrmsf): natural logarithm of the normalized root mean square fluctuation
MD: Maxwell’s demon
MI: mutual information
PCA: principal component analysis
PES: peripheral expression system (the remainder of the genome excluding CP genes)
RQA: recurrence quantification analysis
SD: standard deviation
SOC: self-organized criticality
T-SLM: Thermodynamically gated sequential logic model
WES: whole expression system

## Author Contributions

M.T. conceived and designed the study; developed the theoretical framework and analysis pipeline; performed the analyses; interpreted the results; wrote the manuscript; and supervised the work.

K.Y. conceived and supervised the study based on his current research activity concerning the structure and dynamics of genomic DNA.

O.N. developed the concept of the structural-scaling transition to explain the diversity of Maxwell’s demon scenarios used to interpret the results.

All authors discussed the results and commented on the manuscript.

## Acknowledgments

The authors thank Alessandro Giuliani for insightful discussions and Midori Hashimoto for assistance with the MCF-7 cell data. M.T. is also grateful to his family, Takako Tsuchiya, Kimiko Tsuchiya, Kazumi Tsuchiya, and Harry Taylor, for their constant support. In particular, M.T. is deeply grateful to his late mother, who passed away from cancer; her experience guided him from theoretical physics to cancer research.

## Supplementary file

This supplementary file examines the reproducibility of the findings across independent experimental replicates.

Figure S1 shows that the principal conclusions are reproduced across the two MCF-7 replicates: the HRG replicates show the same 15-20 min commitment window and subsequent 20-30 min Feedback execution, whereas the EGF replicates retain the non-commitment architecture despite some quantitative differences. Notably, CP-PES phase synchronization in expression space occurs both in information-thermodynamic metrics and EED work:

- **HRG: Reproducibility of the commitment-to-execution sequence.** Replicate 2 reproduces the principal early information-thermodynamic and dynamical features observed in Replicate 1. Both replicates show CP-bimodal singular behavior at 15-20 min, the global minimum of *I*_net_(CP;PES) at 15 min, maximal positive Δ*S*(CP) and Δ*I*_net_(CP;PES) during the 15–20 min Measurement phase, the incoming CP-work maximum at 15-20 min, and GE switching during 20-30 min Feedback. CP-associated information and entropy dynamics are also strongly synchronized in both replicates, with *r*_CP_ ≈ 0.99 between *I*_ne_t(CP;PES) and *S*(CP), and their changes likewise showing *r*_ΔCP_ ≈ 0.99. Thus, the 15-20 min commitment window and the 20-30 min execution phase are reproduced across both replicates. Quantitative differences remain: the CP-work maximum is substantially smaller in Replicate 2, the Preparation phase begins at 0 min rather than 10 min, and the CP region is shifted along the *ln*(*nrmsf*) coordinate. These quantitative differences do not alter the reproduced timing of the Measurement phase, CP-work maximum, Feedback phase, or GE switch.
- **EGF: Reproducibility of the non-commitment pattern.** Replicates 1 and 2 both retain strong CP-associated information/entropy synchronization, with temporal Pearson correlations of *r*_CP_ ≈ 0.99 and *r*_ΔCP_ ≈ 0.99 within each replicate. However, they differ quantitatively in the amplitudes of information-thermodynamic (IT) metrics and CP work, as well as in their interaction-flux dynamics. Both replicates show a local CP-work maximum within the MD cycle and a work-associated GE switch at 4-6 h, followed by a sharp decrease in CP work at 6-8 h. Replicate 2 additionally shows an earlier interaction-flux reversal at 20-30 min. At 4-6 h, positive CP work coincides with decreasing WES entropy, whereas the subsequent increase in WES entropy at 6-8 h occurs after CP work has sharply decreased. Thus, both replicates reproduce the non-commitment pattern described in **criterion (iv)** in Section 2.4.5: CP-PES synchronization occurs, but information/entropy amplification and CP-work amplification are not coordinately coupled within the same MD-cycle interval.

The marked CP-work increase after 24 h in Replicate 1 and the larger 4-6 h CP-work response in Replicate 2 may reflect differences in proliferation-related dynamics. This interpretation is consistent with the proliferative EGF response reported by Nagashima et al. (2007), although replicate-specific differences in proliferation remain a hypothesis requiring direct measurement.

**Figure S1.**
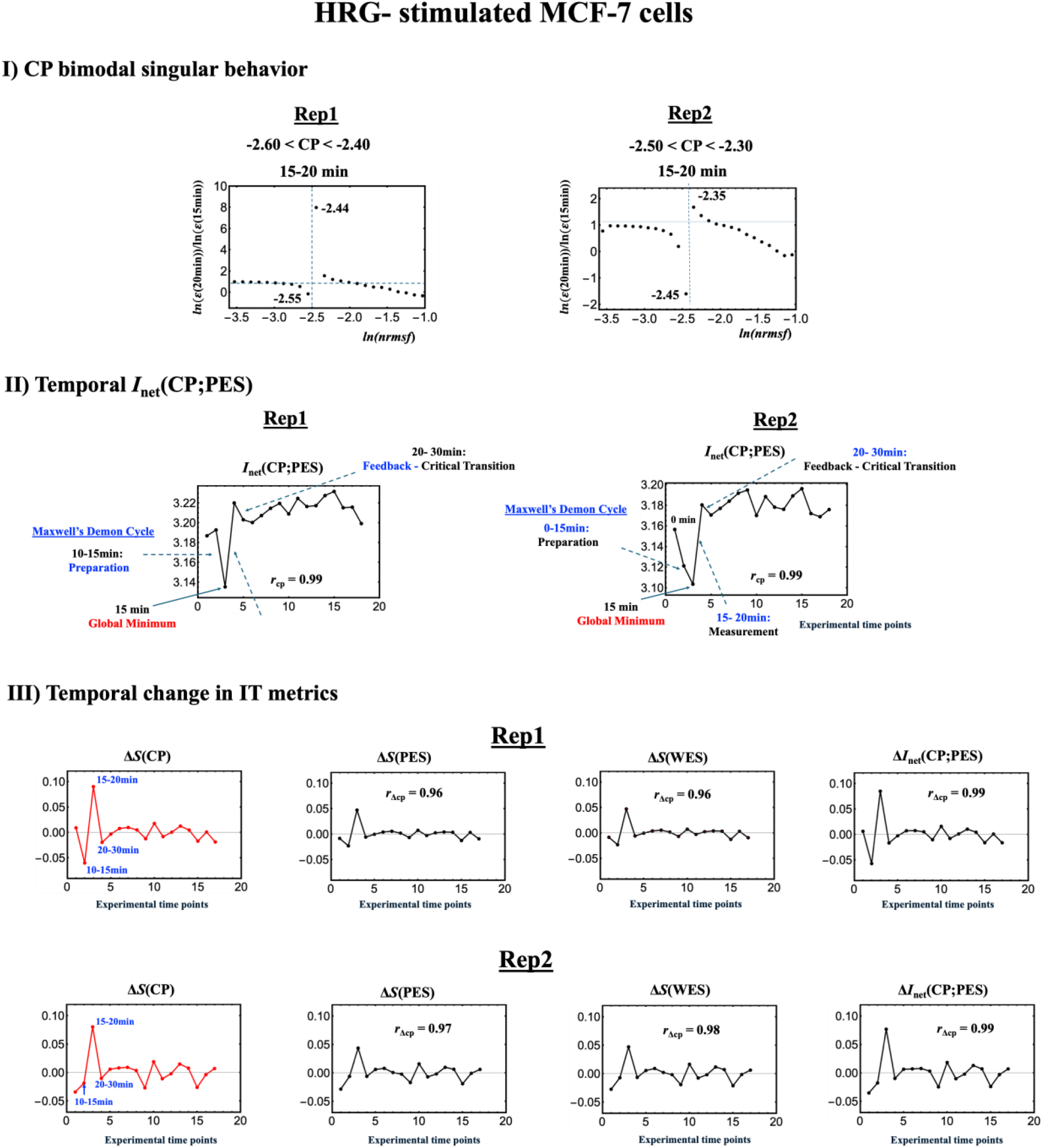
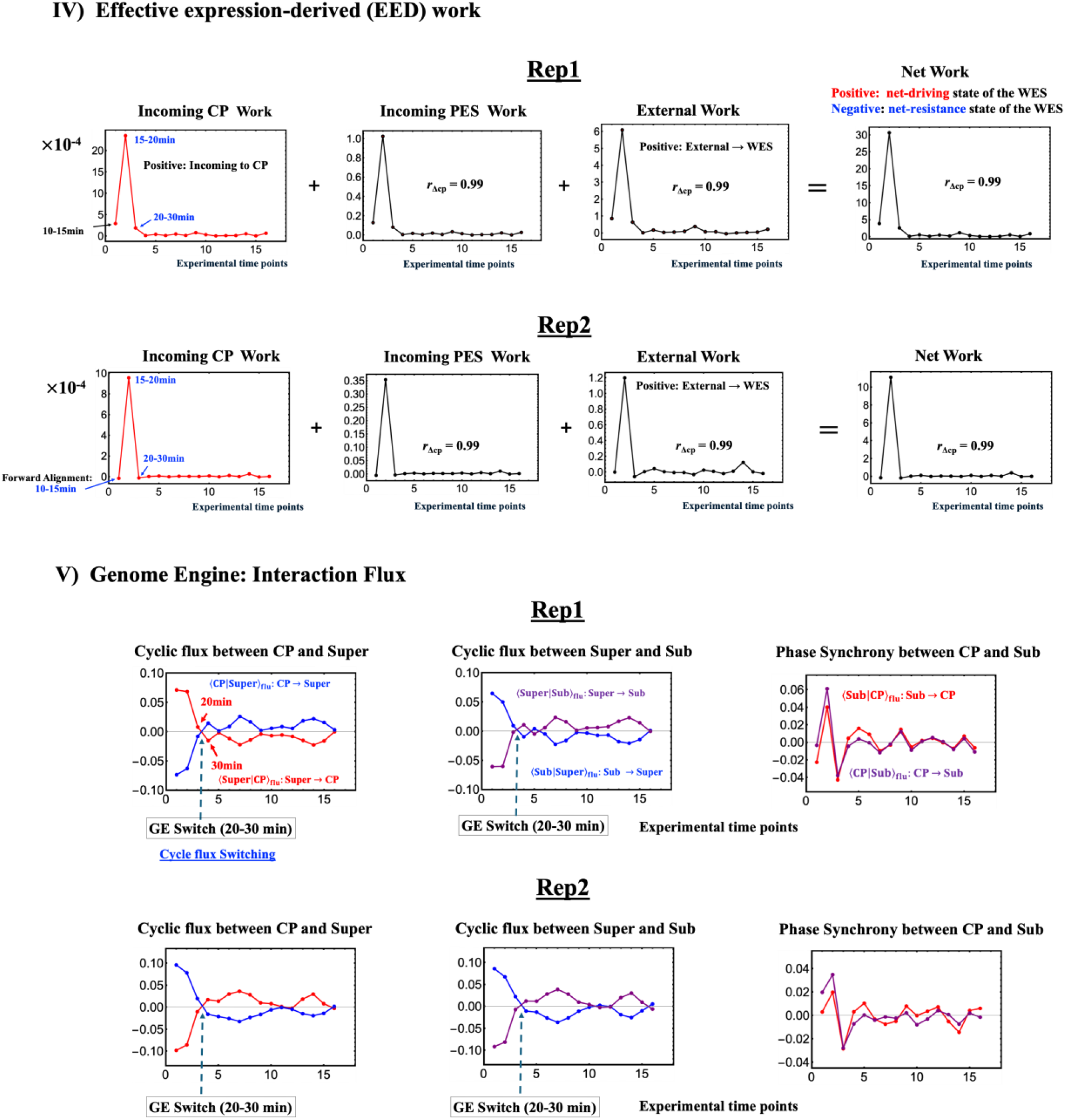
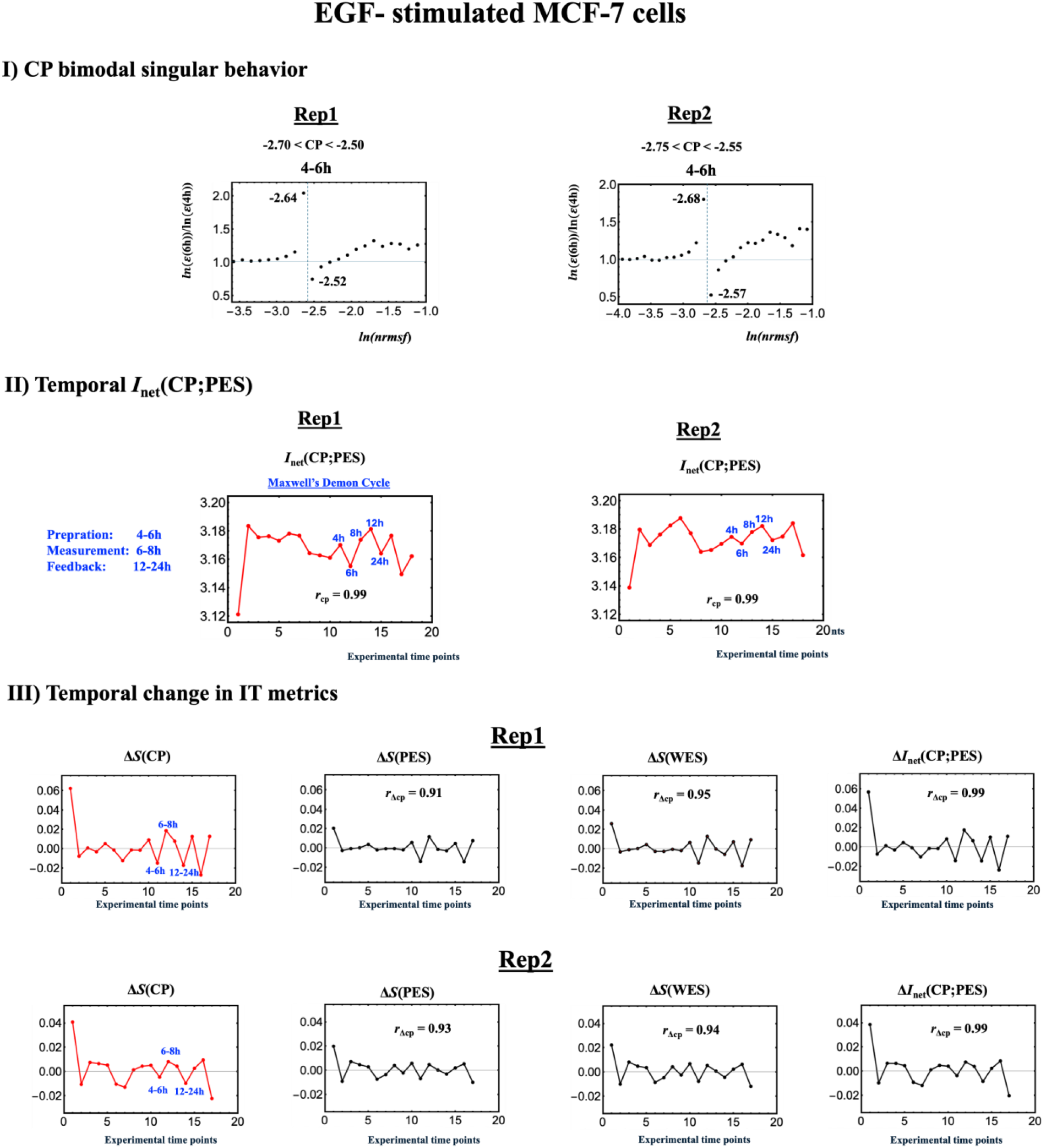
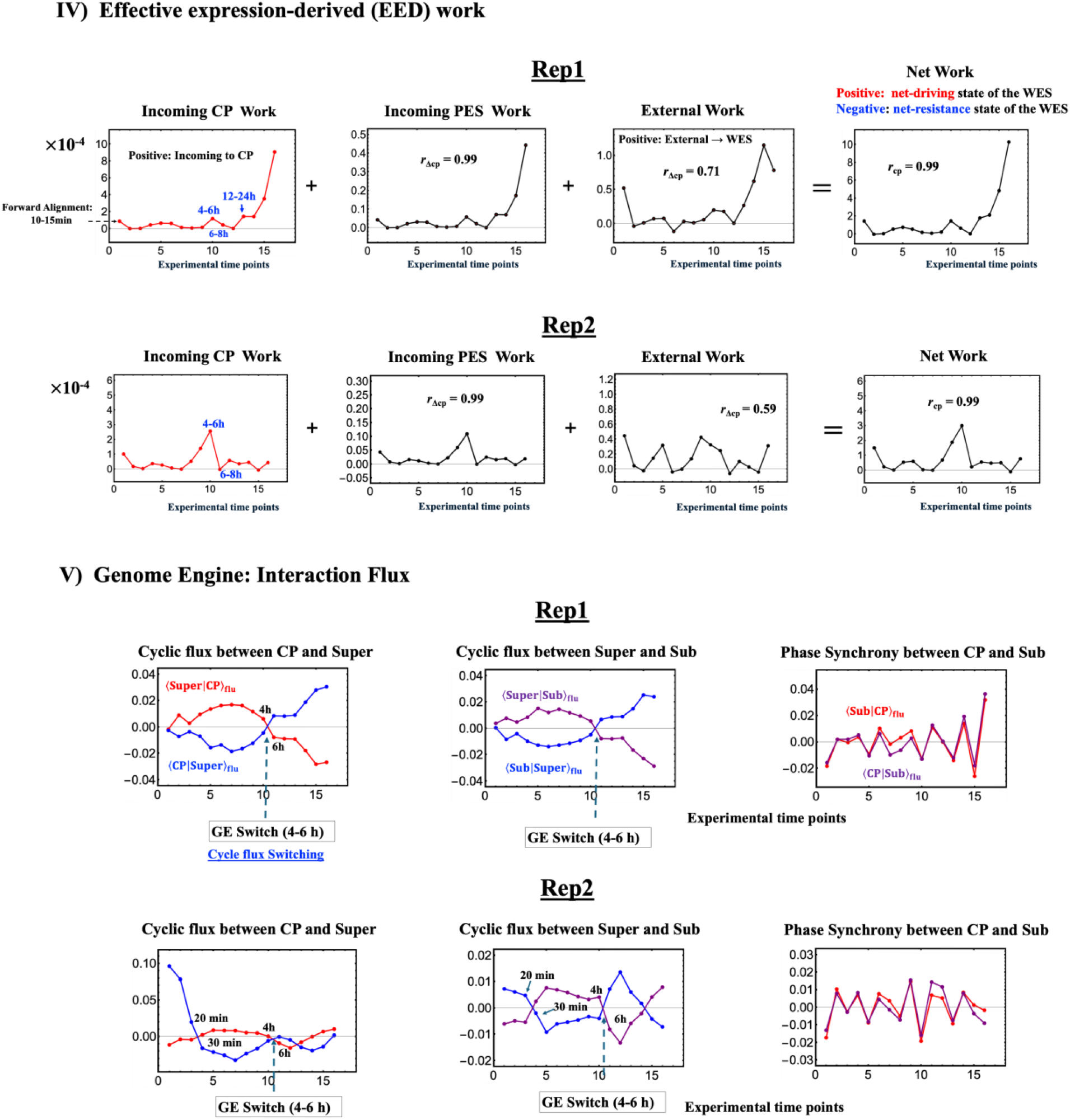
Reproducibility of information-thermodynamic and genome-engine dynamics across MCF-7 replicates. HRG and EGF Replicates 1 and 2 are compared using the same analyses. *r*_CP_ and *r*_ΔCP_ denote the temporal Pearson correlations with *S*(CP) and Δ*S*(CP), respectively. **I)** CP singular behavior along the *ln*(*nrmsf*) coordinate. **II)** Temporal dynamics of *I*_net_(CP;PES), including the Maxwell’s demon cycle. **III)** Temporal changes in information/entropy quantities; the upper and lower rows show Replicates 1 and 2, respectively. **IV)** Incoming CP and PES EED work, external EED work, and net EED work; the upper and lower rows show Replicates 1 and 2, respectively. **V)** Interaction-flux dynamics and GE switching; the upper and lower rows show Replicates 1 and 2, respectively. For HRG, both replicates reproduce the 15-20 min commitment window and 20-30 min Feedback execution. For EGF, both replicates retain the non-commitment architecture despite some quantitative differences.

